# Cell specialization and coordination in *Arabidopsis* leaves upon pathogenic attack revealed by scRNA-seq

**DOI:** 10.1101/2023.03.02.530814

**Authors:** Etienne Delannoy, Bastien Batardiere, Stéphanie Pateyron, Ludivine Soubigou-Taconnat, Julien Chiquet, Jean Colcombet, Julien Lang

**Affiliations:** Université Paris-Saclay, CNRS, INRAE, Université Evry, Institute of Plant Sciences Paris-Saclay (IPS2), 91190, Gif sur Yvette, France; Université Paris Cité, CNRS, INRAE, Institute of Plant Sciences Paris-Saclay (IPS2), 91190, Gif sur Yvette, France; UMR MIA Paris-Saclay, Université Paris-Saclay, AgroParisTech, INRAE, 91120 Palaiseau, France

**Keywords:** scRNA-seq, plant defense responses, plant immunity, plant susceptibility, *Arabidopsis* / *Pseudomonas* interactions, biotic stress

## Abstract

Plant defense responses involve several biological processes that allow plants to fight against pathogenic attacks. How these different processes are orchestrated within organs and depend on specific cell types is poorly known. Here, using scRNA-seq technology on three independent biological replicates, we identified 10 distinct cell populations in wild-type *Arabidopsis* leaves inoculated with the bacterial pathogen *Pseudomonas syringae* DC3000. Among those, we retrieved major cell types of the leaves (mesophyll, guard, epidermal, companion and vascular S cells) to which we could associate characteristic transcriptional reprogramming and regulators, thereby specifying different cell-type responses to the pathogen. Further analyses of transcriptional dynamics, based on inference of cell trajectories, indicated that the different cell types, in addition to their characteristic defense responses, can also share similar modules of gene reprogramming, allowing for instance vascular S cells, epidermal cells and mesophyll cells to converge towards an identical cell fate, mostly characterized by lignification and detoxification functions. Moreover, it appeared that the defense responses of these three cell types can evolve along a second separate path. As this divergence does not correspond to the differentiation between immune and susceptible cells, we speculate that this might reflect the discrimination between cell-autonomous and non-cell-autonomous responses. Altogether our data provide an upgraded framework to describe, explore and explain the specialization and the coordination of plant cell responses upon pathogenic challenge.

## Introduction

In presence of pathogens, plant responses are characterized by a combination of immune and susceptible processes whose balance ultimately determines the degree of plant resistance. Plant immunity is traditionally described as a two-layered system relying on two distinct perception machineries ^1^. In the first one, conserved Pathogen-Associated Molecular Patterns (PAMPs) are recognized by surface-localized Pattern-Recognition Receptors (PRRs), resulting in PAMP-Triggered Immunity (PTI). In the second layer, effectors that are secreted by the pathogens in the plant cells are recognized by cytoplasmic Nucleotide-Binding Leucine-rich Receptors (NLRs) leading to effector-Triggered Immunity (ETI). The arsenal of PRRs and NLRs are particularly rich in plants and allow them to mount effective defense responses against a large scope of pathogens with different trophic lifestyles ^2–4^. Recent advances also showed that PTI and ETI do not necessarily constitute independent types of immunity but can be interconnected at different nodes, reciprocally potentiating each other. Thus, PTI is needed for fully efficient ETI caused by various effectors, while ETI promotes the expression of key PTI actors ^5, 6^. Consistently PTI and ETI appear to converge towards a comparable gene reprogramming that is constantly fine-tuned or modified by the emergence and imbrication of new signaling like the ones mediated by the hormones salicylic acid (SA), jasmonates (JA), ethylene (Et), and abscisic acid (ABA) ^7–10^. In parallel, plant susceptibility is generally understood as the outcome of impaired immune responses, usually due to the action of unrecognized pathogen effectors able to counteract and sabotage key immune actors. In this case we talk of effector-Triggered Susceptibility (ETS) ^1^.

Despite the fact that PTI, ETI and ETS provide a powerful framework to explain plant defense responses, the temporal and spatial dimensions through which they exert their actions remain poorly characterized. Indeed, plant organs are composed of different tissues and cell types which therefore might respond differently to pathogen presence and give rise consequently to specific cell-type immunity or susceptibility. Moreover, pathogen colonization is a dynamic process that starts at specific entry points and then spreads throughout the host organs along possibly preferential paths, depending on how the various plant cell-types can integrate this progression ^11^. Also, communication between cells that are directly sensing pathogens and cells that are still safe might stimulate, even within a population of the same cell type, two different kinds of responses (cell-autonomous responses for those in direct contact with the pathogens and non-cell-autonomous for the others) ^12^. Till recently these questions could not be directly addressed due to technical limitations. Nonetheless cell-type immunity is a well admitted concept in the field. For instance, it is common to mention stomatal immunity as a guard cell-specific defense process preventing pathogens to penetrate tissues and colonize plant apoplast, even if the underlying molecular mechanisms are not totally elucidated so far ^13^. In line with this, a recent study focusing on *Arabidopsis* root epidermis, cortex, and pericycle cells, also showed that PTI responses were mostly driven by cell identity, a finding that reinforces the validity of cell-type immunity ^14^. Yet a comprehensive understanding of how the complex heterogeneity of cell types might affect defense response is still lacking. In this regard newly-developed single cell technologies constitute unprecedented tools to undertake such investigations ^15^.

The bacterial *Pseudomonas syringae* pv. *tomato* DC3000 (*Pst*) strain is routinely used as a model hemibiotrophic pathogen of the plant phyllosphere. Its virulence is mediated by 32 effectors that are injected in the plant host through a type III secretory system. Among those, 8 are of special importance to promote colonization, notably by creating an aqueous apoplast environment ^16, 17^. In addition, *Pst* produces the toxin coronatine, a JA mimic, that antagonizes SA defense pathways, and stomatal closure ^18, 19^. The presence of *Pst* in the plant leaves leads to what is termed basal resistance or basal immunity, i.e., a combination of PTI, ETI, and susceptibility due to the effects of *Pst* virulence factors. In the literature we can find the description of several actors and processes that support basal immunity, including hormone signaling ^20^, kinase signaling ^21^, ROS production ^22^, response to hypoxia ^23^, response to unfolded protein ^24, 25^, lignification ^26^ or production of defensive secondary metabolites like camalexin ^27^ and glucosinolates ^28^. Different time-course transcriptional analyses of plant leaves challenged with *Pst* are also available, offering a dynamic view of basal immunity ^29, 30^. However, all these studies were performed on bulk samples and therefore do not allow to determine whether the described defense responses take place in particular cell populations.

Here, using scRNA-seq technology, on *Arabidopsis* leaf cells inoculated with *Pst*, we could reveal distinct cell classes, characterized by both specific and common defense transcriptional dynamics, thereby unveiling a new and original map of the battlefield between plants and pathogens.

## Material and Methods

### Plant materials and growth conditions

Plants from this study are in the Columbia background. They were grown in soil, in growth chambers at 20 °C, in short day conditions (8 h light, from 9 am to 5 pm), at 60 % hygrometry and under a light intensity of approximately 150 μmol m^-2^ s^-1^.

### Bacterial infections

Bacterial infections were performed on 6 weeks-old plants by spraying. The *Pseudomonas syringae* pv. *tomato* DC3000 WT strain was grown on LB medium supplemented with 50 µg/ml of rifampicin. Fresh cultures in mid-exponential phase were washed, resuspended in 10 mM MgCl_2_ at a final OD_600_=0.1, and supplemented with 0.04 % of Silwet L-77. For the three independent replicates, spraying took place at the end of the day (between 5:30 pm and 6 pm), and leaves were collected 16 h later (between 9:30 am and 10 am). During this period, plants remained uncovered.

### Protoplast preparation, and scRNA-seq library preparation and sequencing

For each biological replicate, about 20-25 leaves coming from 8 independent plants were cut in thin slices with a razor blade, and digested in an enzymatic solution (1.5 % Cellulase R10, Yakult, 0.4 % Macerozyme R-10, Yakult, 400 mM Mannitol, 20 mM KCl, 20 mM MES pH = 5.7, 10 mM CaCl_2_, 0.1 % BSA), under vacuum at RT for 15 min, and then in growth chamber (20 °C) for 50 min. The protoplast solutions were filtered through a 70 µm mesh and a 40 µm cell strainer, with mild centrifugation (1 min, 150 g), and resuspended each time in W5 buffer (150 mM NaCl, 125 mM CaCl_2_, 5 mM KCl, 2 mM MES pH = 5.7). After concentration measurements using a hemocytometer, protoplasts were recovered by centrifugation and resuspended in PBS buffer supplemented with 0.04 % of BSA at a final concentration of 1000 cells / µl. 8000 protoplasts were then processed using the Next GEM Single Cell 3’ library kit v3.1 (10x Genomics) following the supplier’s instructions, and the libraries were sequenced on an Illumina NextSeq500 at the POPS platform (https://ips2.u-psud.fr/en/platforms/spomics-interactomics-metabolomics-transcriptomics/pops-transcriptomic-platform.html.)

### scRNA-seq data processing

For each biological replicate, sequencing data were preprocessed with Cellranger (v4.0.0; 10x Genomics) using default parameters and the Araport11 as *Arabidopsis* reference genome. The preprocessed data were pooled by concatenation and analyzed with the Seurat R package (v3.2.2) ^31^. Genes expressed in 3 cells or less as well as cells expressing 200 genes or less were removed from the analysis. The tolerated percentages of mitochondrial and chloroplastic reads per cell were set at 5 %. Cells were normalized via the SCTransform function and PCA was performed on the 2000 most variable features using the FindVariableFeatures function where 30 dimensions were kept. We clustered the cells with the FindClusters function, using the Louvain algorithm and a resolution r = 1. The neighbors used to perform the clustering algorithm were found using the FindNeighbors function. The list of marker genes was generated using the FindAllMarkers function with the min.pct and logfc.threshold parameters set at 0.25.

The Monocle3 package (v1.0.0) ^32^ (https://cole-trapnell-lab.github.io/monocle3/) was used for analysis of transcriptional dynamics. Seurat object was transferred into a Monocle3 cell_data_set object using SeuratWrappers (v0.3.1) (https://github.com/satijalab/seurat-wrappers). The cluster_cells function was used to compute partitions needed for trajectory analysis and then clusters were reassigned to previously computed clusters found with Seurat. Cell trajectories were built using the learn_graph function. Differential gene expressions across trajectories were found using graph_test, and only genes with high confidence (q-values< 5.10-10 on the Moran’s I test) were retained to form the gene modules. Gene modules were obtained using the find_gene_modules function with a resolution r = 0.01.

For visualization, the gene expression space was reduced using Uniform Manifold Approximation and Projection (UMAP), the violin plots were generated with the VlnPlot function (Seurat). The expression profiles of individual genes, in cells and in clusters, were obtained using the plot_cell function, and the plot_gene_by_group function (Monocle3). The expression scores of gene modules, in cells and in clusters, were obtained using the plot_cell function and the aggregate_gene_expression function (Monocle3).

## Results

### scRNA-seq analysis reveals 10 relevant distinct cell populations in *Arabidopsis* leaves challenged with *Pst*

To investigate cell-type defense responses, we spray-inoculated *Arabidopsis* plants with *Pst*. Leaves were collected 16 h later and subjected to a short digestion protocol to produce protoplasts which were then processed through the microfluidic Chromium technology (10x Genomics) to generate scRNA-seq libraries. We repeated this experimental procedure three independent times and obtained three independent dataset we pooled by concatenation. The cell clustering was carried out on the pooled dataset and resulted in the identification of 18 distinct cell populations (C0 to C17) including 11206 high quality cells. The median numbers of genes and reads per cell are 2386 and 7676 respectively. The UMAP projection of the 18 clusters is shown in Figure S1. The numbers of cells in each cluster range from 1003 for the largest C0 cluster to 61 for the smallest C17 cluster (Table S1).

To understand the biological relevance of the 18 cell populations, we first analyzed the expression levels, in the different cells of the different clusters, of various cell-type marker genes found in the literature. These genes included markers of mesophyll cells, epidermal cells, guard cells, companion cells, vascular S cells and hydathode cells ^33–35^. Remarkably this method allowed us to identify in a relatively straightforward manner the cell identities of 11 clusters (Figure S2). C0, C3, C4, C8, C10, and C17 are thus a priori mesophyll cells, C9 and C1 are a priori epidermal cells, while C2, C15 and C16 correspond a priori to vascular S cells, companion cells and guard cells respectively. Even if they show high expression levels of mesophyll marker genes, the identities of C5, C6, C7, C11, C12, C13 and C14 remained at this step uncertain. No cluster could be attributed to the hydathode identity, maybe because our experimental conditions did not permit the recovery of such cell type.

Next, we created a list of upregulated marker genes for each cluster (Table S2). This list displays several hundred of marker genes for all clusters, except C14 which displays only 43 marker genes. We therefore considered C14 as a clustering artifact and removed it from subsequent investigation. Using the list of marker genes we performed Gene Ontology (GO) analysis, and revealed in the different clusters specific enrichments in biological processes related to stress responses and to cell identity (Table S2). Consistently with their mesophyll annotation, C0, C4, C8, and C10 show the strongest enrichments in processes related to photosynthesis and translational elongation. Similarly, C7, C11, and C12 show strong and specific enrichments in these processes, supporting the notion that those clusters are also mesophyll cells. Besides, C7, C8 and C12 show enrichments in processes like Et, SA and JA signaling pathways, responses to hypoxia and ROS, or response to bacterium, suggesting that these three clusters correspond to responsive mesophyll cells whereas C0, C4, C10 and C11 correspond to healthy mesophyll cells. The C8 notably shows a specific enrichment in response to unfolded protein. The epidermal C9 is characterized by fluid transport, cuticle development, and wax and cellulose biosynthetic processes, while C1 shows enrichments in processes related to responses to chitin, hypoxia, water deprivation, JA / SA / Et / ABA-signaling pathways as well as regulation of signal transduction and regulation of cell-to-cell communication, suggesting that C9 and C1 correspond to healthy and responsive epidermal cells respectively. Interestingly the companion cell C15 shows a strong enrichment in polyamine synthesis, especially in putrescine synthesis, an observation that can be related to the role of polyamines in biotic stress responses and their apparent privileged transport through phloem tissues ^36, 37^. C16, identified as the cluster of guard cells, appears to be mostly driven by ABA-mediated processes, which is congruent with the well-established role of ABA in the regulation of stomata aperture and water transport^38^. At last, C2 shows specific and strong enrichments in hydrogen sulfide and glucosinolate biosynthetic processes, confirming that C2 should encompass the vascular S cells of the phloem parenchyma in which glucosinolate biosynthesis typically occurs ^39^. In addition, our GO analysis found that four clusters were characterized only by processes related to stress responses. Thus C3, C5, C6 and C13 show strong enrichments in JA signaling, lignification, detoxification and response to water deprivation. At last, C17 includes preponderantly GO terms associated with nucleosome assembly and chromatin remodeling, suggesting that cells of this cluster could undergo division or endoreplication. To further rationalize our clustering, we looked at the cell distribution between the three biological replicates in each cluster. As shown in Figure S3, we detected drastic replicate bias for some clusters, unveiling important variations between the three independent inoculations. Nonetheless, C1, C2, C5, C8, C9, C13, C15, C16 and C17 exhibit a cell distribution between the three replicates that is in line with the distribution in the total number of cells. We therefore inferred that these 9 clusters contain the most robust biological information. Remarkably we retrieved among them all the cell types we identified previously through the marker approach and the GO analysis, with the notable exception yet of the healthy mesophyll cells. Indeed, C10 and C11 appear to be specific of replicates 1 and 2 whereas C0 and C4 are composed almost uniquely of cells coming from replicates 1 and 3 respectively (Figure S4). As leaves from the three replicates were exposed to slightly different light periods before harvest (see Material and Methods), an explanation could be that scRNA-seq is powerful enough to capture differences in the photosynthesis program between the three biological repetitions. For simplification, and since this study mostly focuses on the defense responses, we decided to consider only C10 and C11 as representative of healthy mesophyll cells.

Taken altogether our preliminary analyses came to the identification of 10 biologically relevant cell clusters that can serve to further investigate cell-type defense responses. The UMAP projection of these 10 clusters is shown in Figure 1. To further characterize the defense responses of these clusters, we focused on the most strongly and significantly upregulated genes (FCh ≥ 2^0.7^, adjusted p value < 0.05) (Table S3). In the following, marker genes will always refer to this more stringent definition, unless otherwise mentioned.

**Figure 1:**
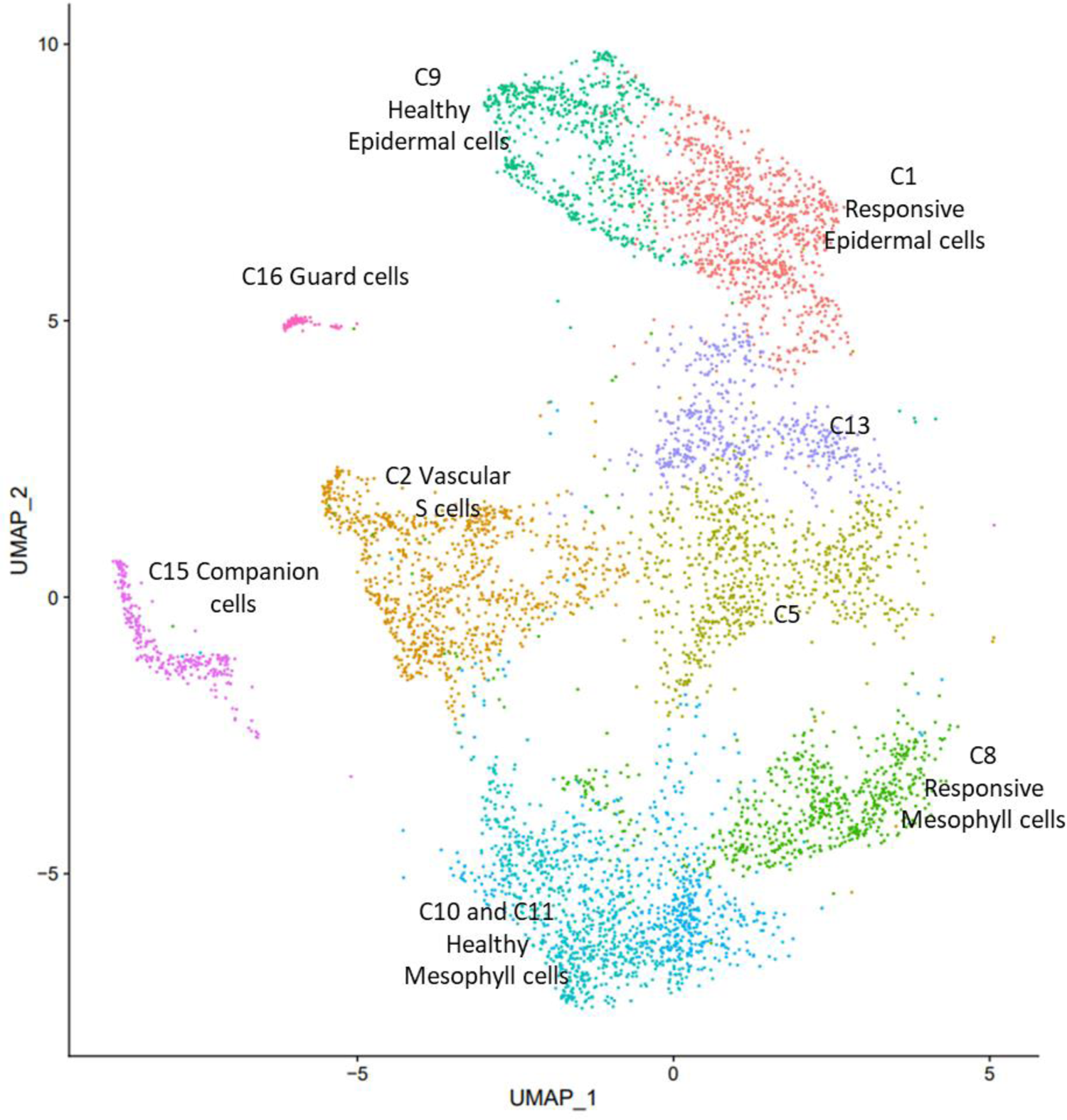
UMAP projection of the 10 relevant clusters. Annotation was performed from expression profiles of cell-type marker genes and GO analysis.

### Characteristic epidermal defense response

To characterize epidermal defense responses, we first compared C9 marker genes with a list of epidermal genes coming from an independent scRNA-seq experiment performed on non-challenged *Arabidopsis* plants of the same age ^33^. We observed a strong overlap between the two sets of genes (64/130 for C9) (Table S3), confirming again the healthy epidermal identity of the C9 cells. Nonetheless, among the genes which are specific to C9 compared to Kim et al. ^33^, we found enrichments for processes related to water transport and response to other organisms. Six plasma membrane intrinsic protein (PIP) aquaporin genes are thus part of the C9 marker genes: *PIP2E*, *PIP1A*, *PIP1D*, *PIP1B*, *PIP3A*, and *PIP2A*. This is consistent with the notion that water allocation is a critical process at the interface between the host and the pathogen ^40^. Regarding genes associated with response to other organisms, we noticed that several of them are involved in Et synthesis and signaling like the *ACC synthase 6* and the *ACC oxidase 2* genes as well as five *Et response Factors*: *ERF017*, *ERF5*, *ERF109, ERF018*, and *ERF72*. In addition, we found among the C9 markers several genes encoding proteins involved in biotic stress response. Those include *AT2G45180*, a nsLTP family-related gene whose expression was recently reported to be suppressed by *Pst* ^41^, *AT4G14450* encoding a proline/serine rich protein that interacts with the immune kinase MPK6 and whose expression is induced by PAMPs ^42^, and *RALF23*, and *PREPIP1* which code for small peptides acting as phytocytokines ^43, 44^. Interestingly, when we plotted the expression levels of all these 17 genes in the UMAP representation, we observed that they showed a progressive decrease between cells from C9 and C1 populations (Figures S5, S6), indicating that these genes are very likely repressed in response to *Pst*.

The C1 cluster that we identified as the responsive epidermal cluster also shares numerous marker genes with the set of healthy epidermal genes from Kim et al., 2021 ^33^ (48/151), (Table S3). C1 and C9 have 12 marker genes in common, 8 of them appearing to be epidermal markers according to Kim et al, 2021 ^33^, the 4 others being *AT1G25275*, a thionin-like gene, *AT1G66090* a TIR-NBS gene upregulated during PTI and ETI ^45^, *AT2G05520* an ABA-, SA-, and Et-responsive gene encoding a glycine-rich protein, and the E3 ligase *AT4G17245*. The expression profiles of these 4 genes showed that they either progressively increase between cells from C9 and C1 populations or remained at the same levels (Figure S7), suggesting either that their induction mediate early epidermal responses to the pathogen or that they are new markers of the epidermal cells. In the specific C1 marker genes (compared to Kim et al., 2021 ^33^), we retrieve many stress genes that overall depict a response to *Pst* mainly characterized by hypoxic stress as well as hormone signaling (JA, ABA, and SA) (Table S3). Remarkably, using TF2Network ^46^ to search for transcriptional regulators that may explain the expressions of the C1 marker genes, we found that the 10 most important transcriptional factors are WRKY transcription factors (Table 1). In contrast no consistent list of transcriptional regulators could be generated for C9.

**Table 1:**
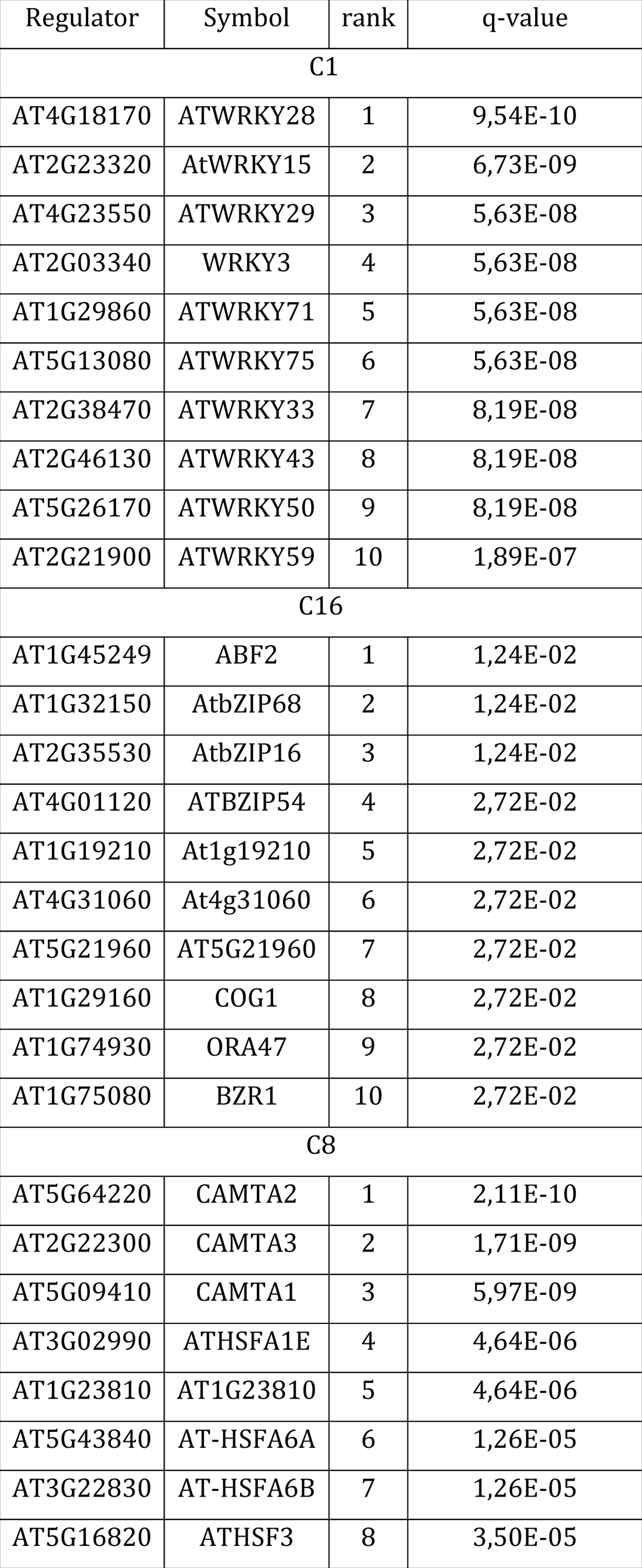
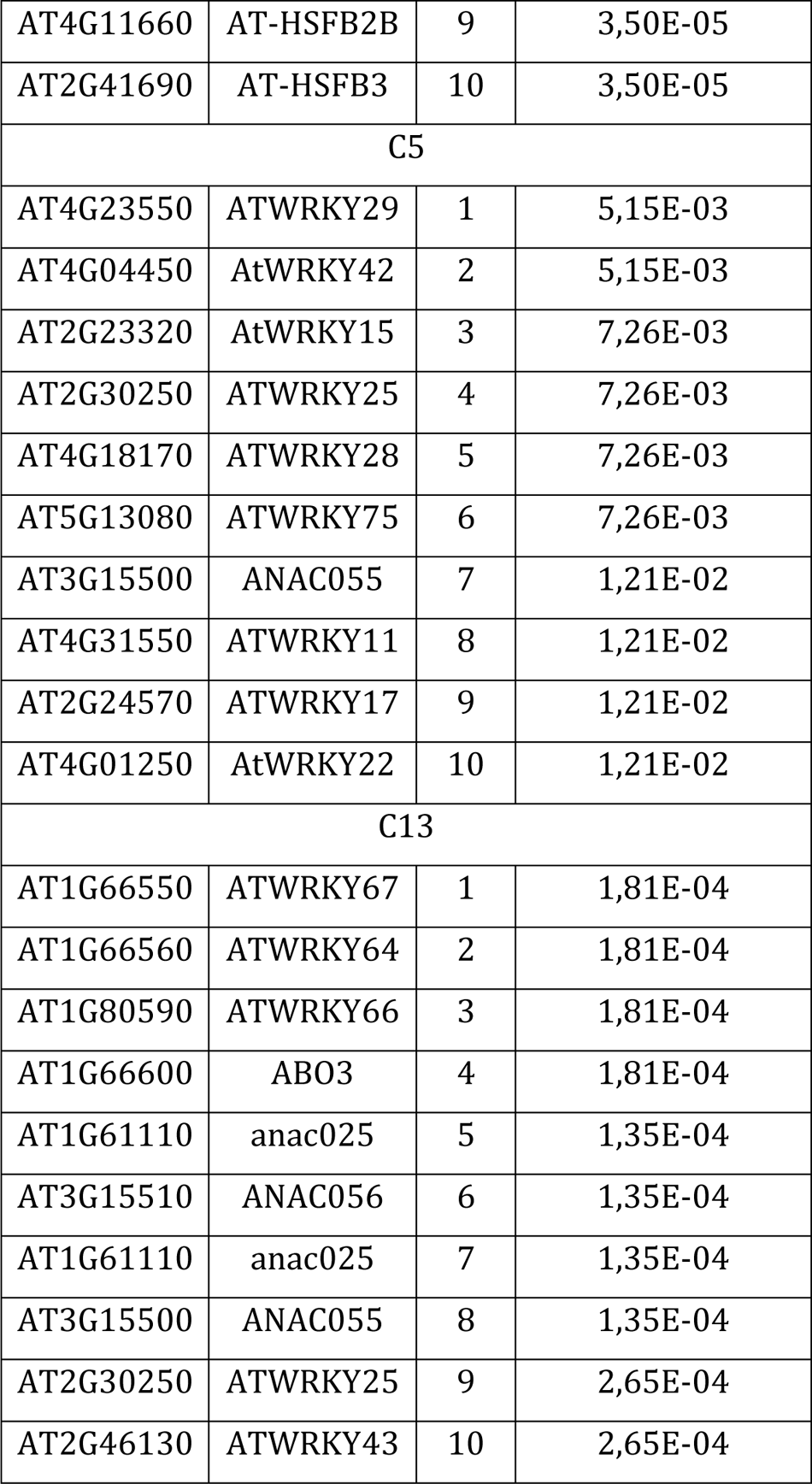
List of regulators for the upregulated genes in C1, C16, C8, C5 and C13 populations. The list was generated using TF2Network (http://bioinformatics.psb.ugent.be/webtools/TF2Network/) and the stringent list of marker genes for each cluster. The regulators are listed according to their ranks and q-values.

Taken collectively the data from C1 and C9 delineate an epidermal transcriptional reprogramming upon *Pst* infection, mainly characterized, on one hand, by the inhibition of water transport and of some Et signaling components, and on the other hand by the activation of WRKY-guided responses to hypoxia, JA, SA and ABA.

### Characteristic guard cell defense response

By comparing the 58 marker genes of C16 with the list of unchallenged guard cell marker genes from Kim et al., 2021 ^33^, we found an overlap of 12 members, notably among the highest expressed genes (Table S3), which comforted us in the attribution of guard cell identity to C16. Specific C16 marker genes compared to Kim et al., 2021 ^33^ revealed important GO enrichments in processes like transport of fluid / water (involving the aquaporin genes *PIP2A*, *PIP1B* and *PIP2B*), transport of one-carbon compounds (like CO_2_) as well as response to desiccation (Table S3) whereas no such enrichments could be detected with the list of healthy guard cell marker genes from Kim et al., 2021 ^33^. These data suggest that ABA plays an important role in the adaptation of C16 cells to *Pst*. Consistently, analysis with TF2Network proposes the ABA Responsive Elements-Binding Factors 2 and three other bZIP transcription factors as the most important regulators for C16 reprogramming (Table 1). Besides, if in the guard cell data from Kim et al., 2021 ^33^, the myrosinase *TGG1* is not detected while *TGG2* is the strongest upregulated gene, in our data, among the C16 marker genes, both *TGG1* and *TGG2* are the two genes with the highest upregulation levels (Table S3), suggesting that *TGG1* induction could be a specific response to *Pst*.

Overall C16 tends to show that guard cell responses are mostly governed by ABA signaling. This is consistent with the known role of ABA in the positive regulation of stomata closure, limiting the ability of the pathogen to colonize the plant host ^38^. In addition, guard cells could also contribute to the mustard oil bomb strategy of defense against *Pst* through the expressions of the myrosinases *TGG1* and *TGG2* which are known to be involved in the production of toxic glucosinolate-derived compounds preventing massive invasion of the pathogen ^47, 48^.

### Characteristic mesophyll defense response

Our preliminary analyses allowed identifying C0, C4, C10 and C11 as encompassing healthy mesophyll cells. In line with this, there is a strong overlap between the marker genes of these four clusters and the mesophyll marker genes of unchallenged plants from the study of Kim et al., 2021 ^33^ (187/240) (Table S3). Moreover the 53 remaining specific marker genes of C0, C4, C10 and C11 are predominantly annotated as involved in photosynthetic process, albeit we retrieved among them *PIP2A*, *PIP1A*, *PIP1B* and *PIP2E*. Again, this observation indicates that suppression of these aquaporin genes might represent an important response to *Pst* as it was already observed in epidermal cells (Figure S5).

In contrast to C0, C4, C10 and C11, C8 seems to correspond to the responsive mesophyll cluster. 30 of the 101 C8 marker genes are mesophyll marker genes. As for the other 71 marker genes, they are mostly stress-related and globally uncover a basal defense response to *Pst* that is somehow reminiscent of the epidermal C1, i.e., characterized by responses to hypoxic and oxidative stress as well as hormone signaling (JA, SA, Et and ABA) (Table S3). Yet responsive epidermal (C1) and C8 marker genes have no members in common, hinting at some decisive differences between the two kinds of responses. For instance, Et signaling seems prevalent in responsive mesophyll cells compared to responsive epidermal cells with notably the upregulation of *MKK9* that is an important actor for Et synthesis and for transduction of Et effects (Table S3) ^49, 50^. Similarly, response to oxidative stress is a GO category that appears characteristic of mesophyll cells (Table S3). Besides, C8 displays specific enrichment in processes related to protein folding, protein oligomerization and response to heat stress with several heat shock proteins, especially small heat shock proteins, among the most strongly expressed marker genes (*HSP17.6A*, *HSP17.6B*, *HSP17.6C*, *HSP17.6II*, *HSP17.4*, *HSP17.4B*, *HSP17.8*, *HSP70B*, *HSP70-4*, *HSP81-2*, and *HSP101*) as well as the heat shock factor *HSFA2* (Table S3). Additionally search with TF2Network revealed that the most important transcriptional regulators for C8 are CAMTA1/2/3, HSFA1E and HSFA6A/B (Table 1).

Altogether it seems that mesophyll responses to *Pst* resemble those of epidermal cells, with, for instance, the downregulation of several aquaporin genes (Figure S5) and the enrichments in GO functions related to stress responses that greatly overlap between the two cell types (Table S3). However, a deeper look at the marker genes and their transcriptional regulators reveals important differences in the molecular mechanisms that underpin their respective responses. Notably, some of the Et signaling components which appear to be repressed in responsive epidermal cells are induced in responsive mesophyll cells (Figure S6), response to ROS stress seems also predominant in responsive mesophyll cells, and finally, whereas characteristic epidermal reprogramming is WRKY-mediated, characteristic mesophyll reprogramming involves predominantly CAMTAs and HSFs.

### Characteristic vascular defense responses

Leaf vasculatures are complex tissues encompassing several cell types ^33^. In our data, we identified only two types of vascular cells: the companion cells (C15) and the vascular S cells (C2), likely because our protocol for protoplast production does not allow us to recover other cell types. However, another possibility we cannot totally rule out is that our two vasculature populations are actually a composite of several other cell subtypes. Among the 127 marker genes of C15, 63 are common with the marker genes of companion cells from healthy leaves ^33^, while the 64 others depict a stress response mostly driven by hypoxia and ABA signaling (Table S3). In those latter, the most strongly expressed genes are two metallothionein (MT3 and MT2B), a thioredoxin (TRX3), two cyclophilins (ROC1 and ROC5), and a dehydrin (HIRD11), suggesting that companion cells could also respond to *Pst* through the implementation of protective measures against ROS (Table S3).

Interestingly the vascular S cells were not found in the study of Kim et al., 2021 ^33^, raising the hypothesis that C2 cluster could be a hallmark of induced responses to *Pst*. C2 marker genes show strong enrichments in metabolism of aliphatic glucosinolates with notably the cytochrome P450 *CYP83A1* or the glucosinolate transporter *GTR1* (Table S3). This indicates that vascular S cells could mostly contribute to defense responses to *Pst* through the mustard-oil bomb strategy, likely in collaboration with the guard cells that exhibit high expression levels for myrosinases.

### C5 and C13 are related stress specific cell populations

In our clustering we identified two cell populations to which we could not assign any cell identity but that are characterized by strong stress responses. C5 shows important enrichments in the lignification process as well as in response to biotic stress like bacterium (Table S3). For instance, we found in the C5 marker genes, *CASPL4D2* and *CASPL1D1* corroborating the recent literature showing that these two genes encode two essential enzymes for the synthesis of lignin in response to *Pst* ^26^. The C5 marker genes also include several genes known to be induced upon pathogenic bacterial challenge like *AT3G22600*, *AT4G22470*, or *AT4G37990* (Table S3).

The C13 population is mainly characterized by phytoalexin synthesis, detoxification, and SA-mediated defense responses, especially SAR (Table S3). For instance, *PAD3* and *CYP71A13* are strongly expressed in C13, indicating that camalexin production is likely specific to this cluster (Figure S8). In addition, several Gluthatione-S transferases, metal transporters or secretory carriers, endochitinases or NDR1-like genes are upregulated in C13. Interestingly we found 4 *WRKY* transcription factors among the C13 marker genes (*WRKY8*, *WRKY29*, *WRKY55* and *WRKY75*), suggesting that these regulators play critical roles for the reprogramming of C13.

We also noticed that, in line with their contiguity in the UMAP projection, C5 and C13 have 12 marker genes in common, including *CASPL4D2* and *CASPL1D1* (Table S3). This suggests that the two cell populations are actually close in terms of transcriptional reprogramming. Consistently similar WRKY and ANAC transcription factors appear to control the expression of the marker genes in C5 and C13 (Table 1). Taken altogether our data indicate that C5 and C13 might form a continuum of response to *Pst*, mostly characterized by immune processes like the production of phytoalexins and lignin that can act as barriers against the propagation of the pathogen ^26, 27^. They also highlight the fact that defense responses can lead to profound modifications, both up and down, of various transcript levels that can eventually alter cell identity in a substantial manner.

### Inequalities in pathogen sensing between cell populations

As we identified characteristic transcriptional reprogramming in different cell types, we were prompted to see whether these differences could be correlated with differences in the ability to sense the pathogens between the cell types. To address this question, we focused on the expression profiles of the *NLR* gene family as well as on those of the Leucine-rich Repeat (LRR) and LysM-containing gene classes of the *PRR* family ^51, 52^. As shown in Figures 2, 3, S8 and S9, our data unveil important differences in the expression levels of the immune receptors between the different cell populations. First, we observed that overall, both the number of expressed receptors and their expression levels increase in the responsive clusters C1, C5, C8, and C13 compared to the non-responsive clusters C9, C10, and C11. Importantly we retrieve among the immune receptors that are the most strongly expressed, genes that are well known to be involved in defense responses to *Pst*, like the *PRRs FLS2*, *EFR*, *BAK1*, and *LYK1* or the *NLRs ZAR1*, *SNC1*, and *NRG1.1*. In parallel we observed that the expression levels of the immune receptors remain globally moderate in the C2 vascular S cells, the C15 companion cells and the C16 guard cells, suggesting that these cell types might, to some significant degree, respond to *Pst* through other receptor machineries. However, it must also be noted that for these cell types we could not separate healthy and responsive populations, hence it is possible that the percentage of cells expressing immune receptors as well as their expression levels remain underestimated in these clusters compared to the responsive C1, C5, C8 and C13. At last, it is worth noting that if the *PRRs* are the most numerous to be strongly expressed in the C1, C5 and C13 populations, whereas the *NLRs* are the most numerous to be strongly expressed in the C8 population. This actually could indicate that epidermal cells are more proficient in PTI whereas mesophyll cells are more proficient in ETI.

**Figure 2:**
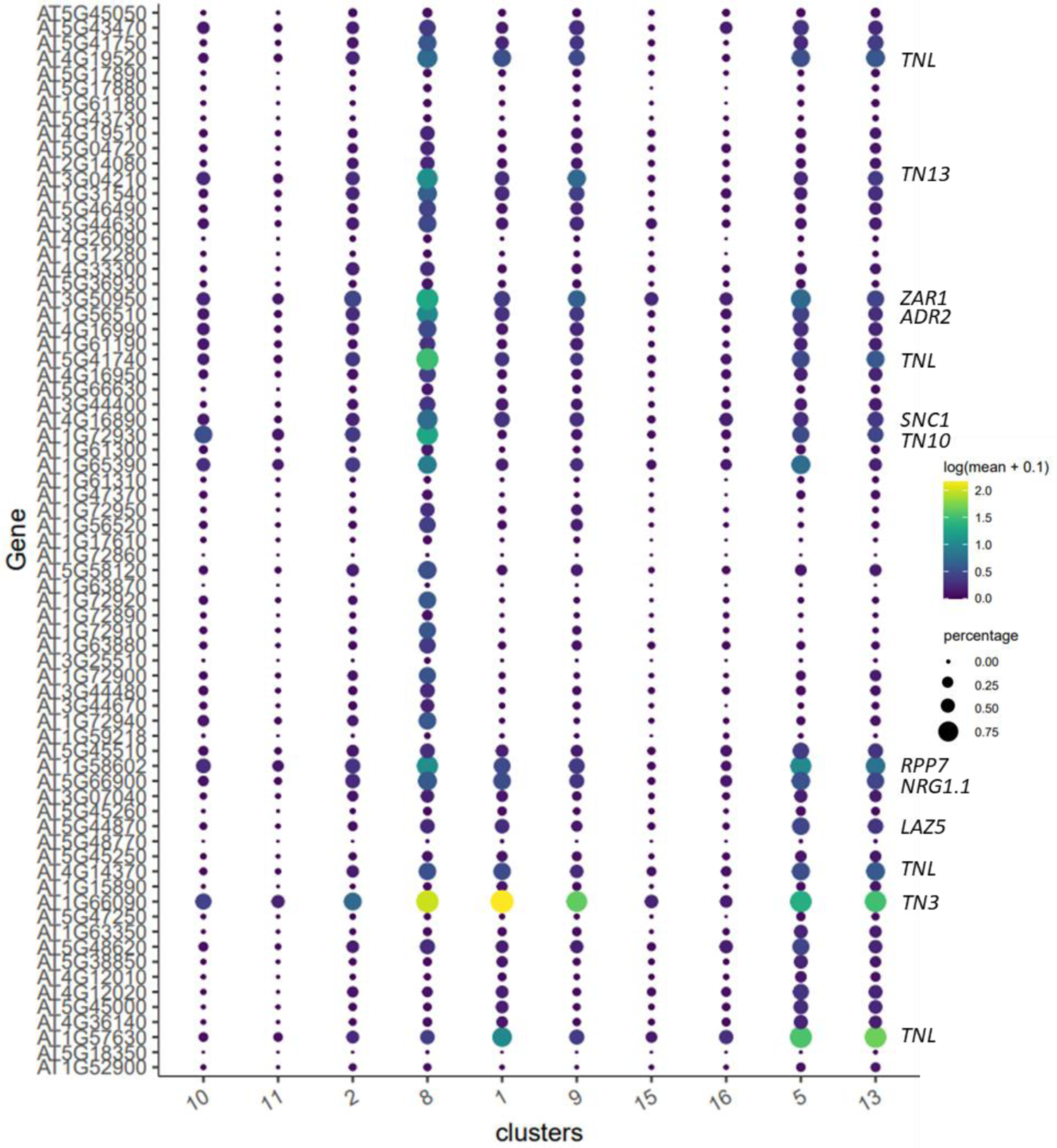
Cluster-specific expression profiles of *NLR* genes. 71 genes from the 207 list (Figure S9) were selected and are shown here. Some gene symbols are given on the right.

**Figure 3:**
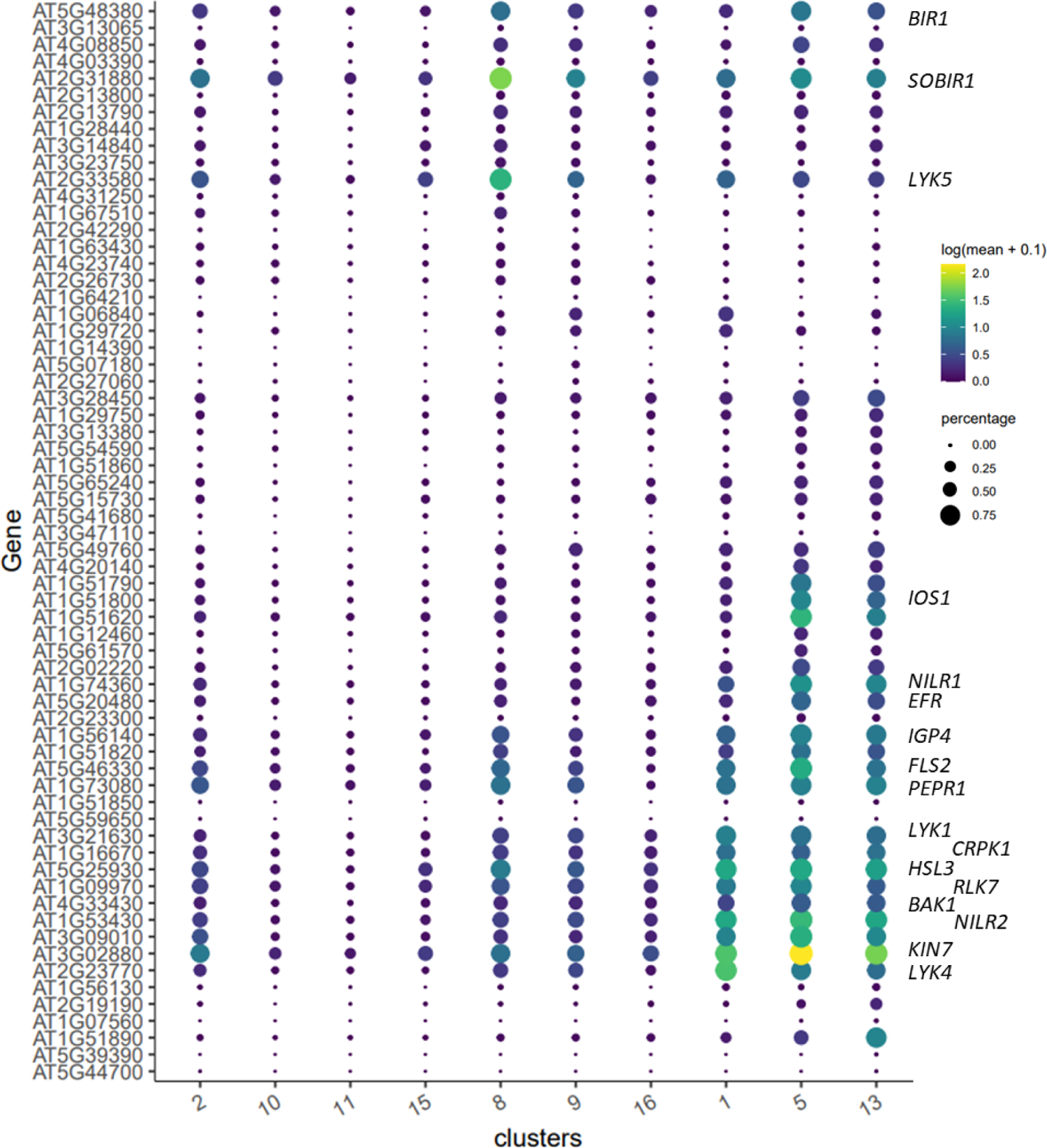
Cluster-specific expression profiles of *PRR* genes. 71 genes from the 236 list (Figure S10) were selected and are shown here. Some gene symbols are given on the right.

### Convergent and divergent evolution of cell types in response to *Pseudomonas*

Till now, our analyses were based on the list of marker genes associated with each of our clusters. If this approach is powerful enough to shed new light on the characteristic defense responses of various cell types, it might however at the same time miss genes that behave similarly in several clusters and can for instance support a general stress response. Moreover, single-cell clustering is not designed to fully apprehend the transcriptional dynamics between cells and how those might mediate different cell evolutions upon *Pst* challenge. To tackle these difficulties, we processed our clustering data with Monocle3 that allows the creation of cell trajectories and the generation of modules that group together genes showing identical expression dynamics along the trajectory within a single cluster or between several clusters. As shown in Figure 4, Monocle3 built three cell trajectories from our clustering data. The first one in the C16 guard cell populations is linear and links two distinct cell fates. The second one is in the C15 companion cells with a branching point and three distinct cell fates. The last one links the C2 vascular S cells, the C1 and C9 epidermal cells, the C10, C11 and C8 mesophyll cells as well as the C5 and C13 cells with ten branching points and ten distinct cell fates. Interestingly it appears that C5 and C13 correspond to a single cell fate (cell fate 10) towards which all vascular S cells, epidermal cells and mesophyll cells could converge. It also appears that these three cell types can diverge along a second trajectory path, at the branching point 6 for epidermal cells, 10 for vascular S cells and 8 for mesophyll cells. Further analysis of the modules generated by Monocle3 allowed identifying genes important to explain the different cell evolutions (Table S4; Figures 4 and S11). For instance, modules 3, 4, 12, 17, 19, 21, 28, 29, 33, 48, 53 and 56 are important to mediate the completion of cell fate 10 (Figures 4 and S11). Without surprise, modules 12 and 48 show strong enrichments in detoxification and lignification, respectively (Table S4), confirming our previous analyses and the importance of these processes in the C5 and C13 populations. If the other modules are globally characterized by defense responses to bacterium, some of them also allow to unveil new functions associated with the C5 and C13 populations, like sphingolipid metabolism for module 19, vacuolar transport and protein targeting to the endoplasmic reticulum for modules 21 and 28, senescence for module 33 and chorismate metabolism for module 56 (Table S4). As chorismate is the precursor of SA, this last observation suggests that cell fate 10 might be important for SA synthesis. To go further we looked at the expression profiles of several genes involved in SA synthesis and signaling. If the results for *SARD1*, *CBP60g* and *PR5* are not really conclusive, the expression patterns of *ICS1*, *PBS3*, *PR1* and *PR2* are indeed compatible with the notion that C13 is a preponderant site for SA synthesis and signaling (Figure S12).

**Figure 4:**
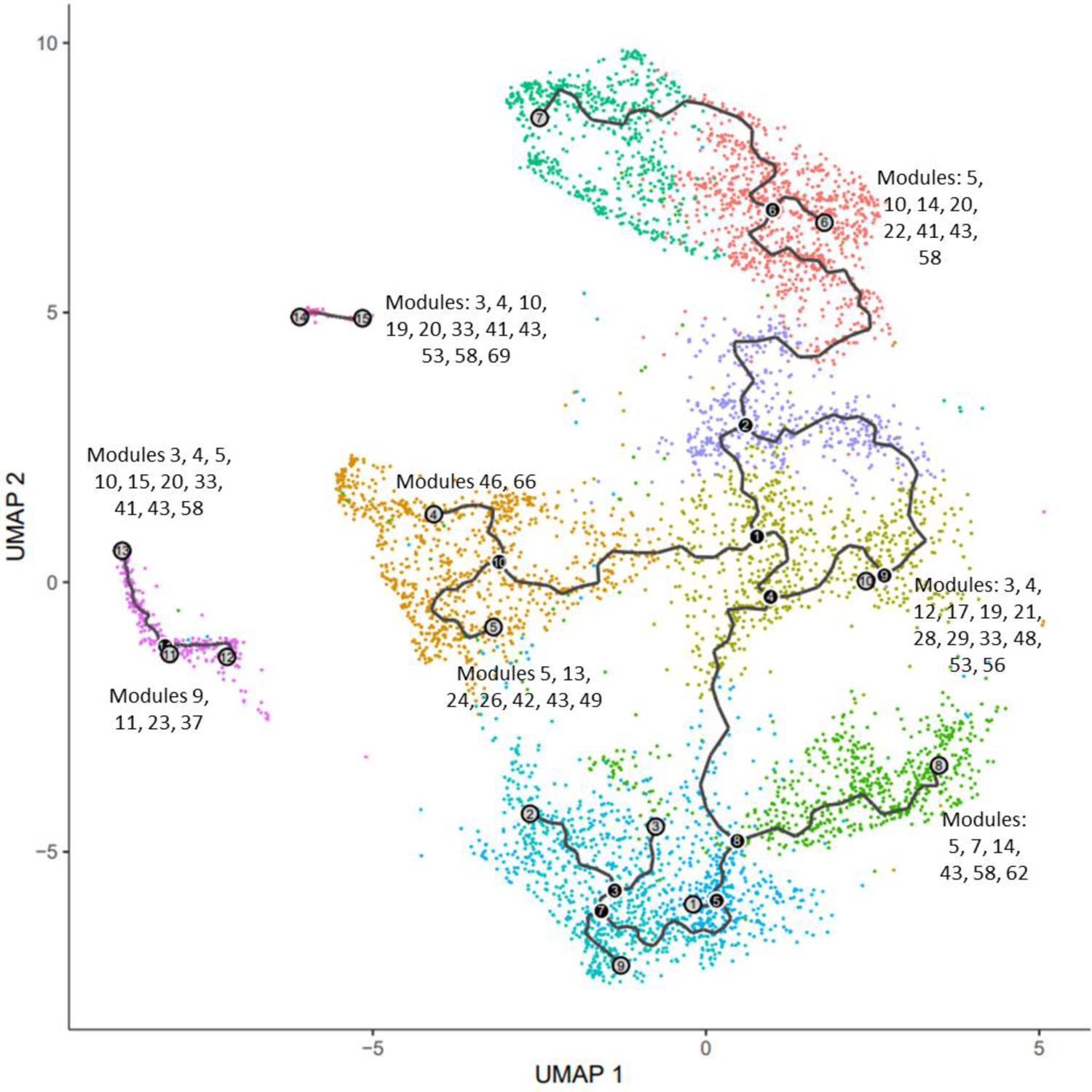
Cell trajectories. Trajectories were built using Monocle3. Black circles indicate branching points. Gray circles indicate cell fates. Texts indicate gene modules showing high expression scores around different cell fates (see Figure S11).

Concerning the epidermal and mesophyll cells, we noticed that their trajectories towards cell fate 6 and 8 respectively are mediated by several common modules (Figures 4 and S11). Those are modules 5, 14, 43 and 58, especially enriched in genes related to responses to bacteria, response to hypoxia and SA signaling (Table S4). In addition, epidermal evolution is characterized by modules 10, 20, 22 and 41 showing enrichments in phospholipid transport, response to bacteria and SA signaling (Table S4), whereas mesophyll evolution is characterized by modules 7 and 62 showing enrichments in response to hypoxia, SA signaling, response to unfolded proteins and response to ROS (Table S4).

Several gene modules involved in the epidermal and mesophyll evolutions also show strong expression scores in the guard cells located around cell fate 15 (Figures 4 and S11), suggesting that these guard cells might correspond to the ones that are the most responsive to *Pst*. The modules concerned are modules 3, 4, 10, 19, 20, 33, 41, 43, 53 and 58 (Figures 4 and S11). Besides we found that module 69 is specific to cell fate 15 (Figures 4 and S11). Although this module does not show any peculiar GO enrichment, it includes several genes involved in ABA signaling like the receptor *PYL2* and the anion channel *SLAC1* (Table S4), reinforcing our interpretation that guard cells mostly respond to *Pst* in an ABA-dependent manner, very likely resulting in the closure of the stomata.

Regarding the vasculature cells, it is more complicated to identify to which biological processes the different cell fates correspond, maybe because, as we already mentioned, these cell populations could encompass more cell types than we think. For companion cells, we observed that evolution towards cell fate 13 is supported by modules 3, 4, 5, 10, 20, 33, 41, 43 and 48 that are also associated with responsive mesophyll and epidermal cells (Figures 4 and S11), suggesting that cell fate 13 corresponds to the responsive companion cells. Interestingly module 15 that shows important enrichment in calcium transport, has specific increased expression score towards cell fate 13 (Figures 4 and S11, Table S4), possibly hinting at a specific role for companion cells in calcium signaling and transport during defense responses to *Pst*. In addition, it is possible to associate cell fate 11 with modules 9, 11, 23 and 37 characterized by respiration processes as well as by polyamine synthesis (Figures 4 and S11, Table S4). Whether this fate corresponds to another kind of companion cell responses to *Pst* is not clear. If the enrichment in genes involved in polyamine synthesis suggests that cell fate 11 could encompass responsive cells, the strong overlap between genes from module 23 and gene markers of healthy companion cells ^33^ (Table S4) actually indicates the opposite. For vascular S cells, we pinpointed two modules with very localized expression patterns: modules 46 and 66 showing enrichments in processes related to phloem or xylem histogenesis (Figures 4 and S11, Table S4). If we consider these cells as corresponding to cell fate 4, we may assume that the responses of vascular S cells to *Pst* are associated with cell fate 5 and are driven by the modules 5 and 43, enriched in processes related to response to hypoxia, the modules 13, 26 and 49, enriched in processes related to ribosome processing and translation, and the modules 24 and 42 enriched in processes related to sulfur amino acid metabolism and glucosinolate synthesis (Figures 4 and S11, Table S4).

In our data we could also identify modules that have high expression scores only in healthy populations, indicating that the genes which compose them are actually downregulated in response to *Pst*. For instance, module 6 is specific to healthy mesophyll cells (Figure S11). As its genes show enrichments in processes related to photosystem assembly or chlorophyll metabolism (Table S4), we inferred that mesophyll photosynthesis is actively inhibited in response to *Pst*. Likewise, modules 16, 18 and 25 are specifically expressed in healthy epidermal cells, indicating that some processes related to cell wall composition and cell pigmentation are downregulated during responses to *Pst* in those cells (Figure S11, Table S4). Remarkably, module 45 is also specific to healthy epidermal cells (Figure S11). Since it shows enrichment in processes related to water transport (Table S4), this confirms our previous observation that aquaporin functions are generally repressed in response to *Pst*.

Taken collectively, our analysis with Monocle3 allowed us to further refine our understanding of responses to *Pst*. First, they brought a description of the cell-type characteristic defense responses not only consistent with, but also complementary to the one we proposed based on the list of marker genes, raising several new hypotheses regarding the specialization of the different cell types in response to *Pst*. Then they uncovered the existence of common transcriptional defense reprogramming shared by several cell types which can lead to similar cell evolutions and identical cell fates. In this regard, the trajectory built by Monocle3 and depicting a divergence of the epidermal cells, vascular S cells, and mesophyll cells between, on one hand, a common immune population and, on the other hand, cell-type specific responses, is of special interest.

### Cell divergence does not reflect differentiation between immunity and susceptibility

To explain the divergence of epidermal, vascular S and mesophyll cells in response to *Pst*, we wondered whether it might reflect the differentiation between immune and susceptible cells. It is considered that *Pst* mediates plant susceptibility mainly through the activation of the JA and ABA signaling that antagonize the immune effects of the SA signaling ^10, 18, 19^. Yet when we characterized the defense responses specific to a given cell type, we often observed concomitant enrichments in processes related to both JA, ABA and SA. This is notably true for the epidermal and mesophyll cells as well as for the C5 and C13 cells (Tables S2 and S3). Besides we noticed that the module 1 from the Monocle3 analysis that shows enrichments in processes principally related to JA and ABA signaling have high expression scores in all the responsive clusters (Figure S11, Table S4). In line with this, the expression profiles of the ABA synthesis genes and some ABA signaling actors globally indicate that ABA-related processes are ubiquitous in our clustering (Figure 5A). Finally, we looked at the transcript levels of 28 genes that are presented as markers of susceptibility ^53^. As shown in Figure 5B, no clear pattern could be deduced from our data, several of the genes being expressed in numerous cells and at high levels in various populations, regardless of the identities or the nature (healthy/responsive) of these latter. Altogether our data suggest that susceptible processes operate simultaneously in all the cells responsive to *Pst* and that they cannot account for the divergence of the different cell types.

**Figure 5:**
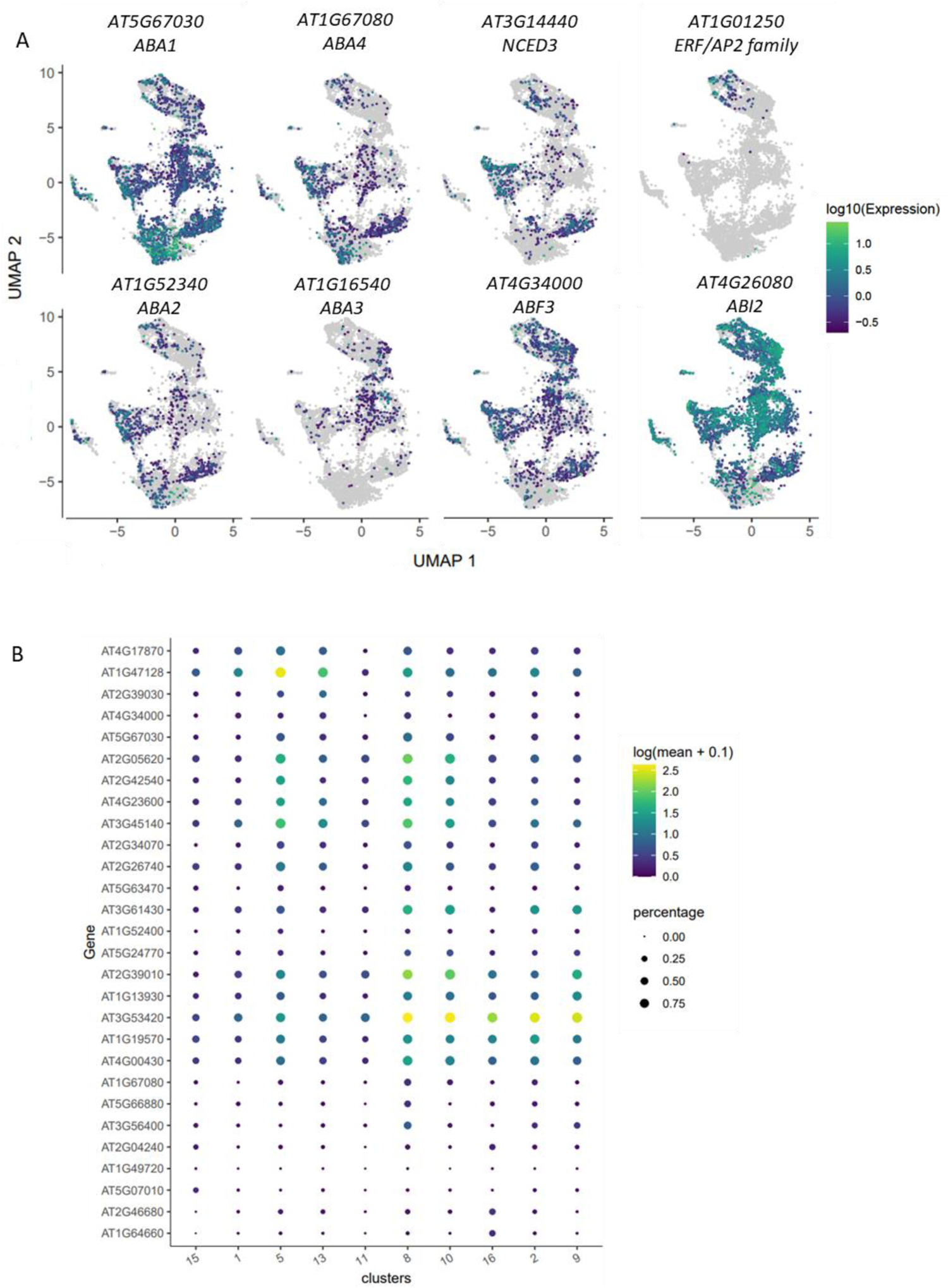
Expression profiles of susceptible genes. (A) Cell specific expression profiles of genes involved in ABA synthesis and signaling. (B) Cluster-specific expression profiles of ABA- and JA-related susceptible genes ^53^.

## Discussion

In this study, using scRNA-seq technology and combining several in silico analyses, we could decipher the plant defense response to a pathogen at an unprecedented single cell resolution. Importantly our data are not only consistent with the current knowledge on the topic, but they also raise several new interesting hypotheses regarding the *Arabidopsis* / *Pseudomonas* interactions, and provide a wealth of information for further molecular investigations. An attempt to integrate all this is presented in Figure 6. In the following, we will discuss briefly how our data bring original elements to three fundamental questions.

**Figure 6:**
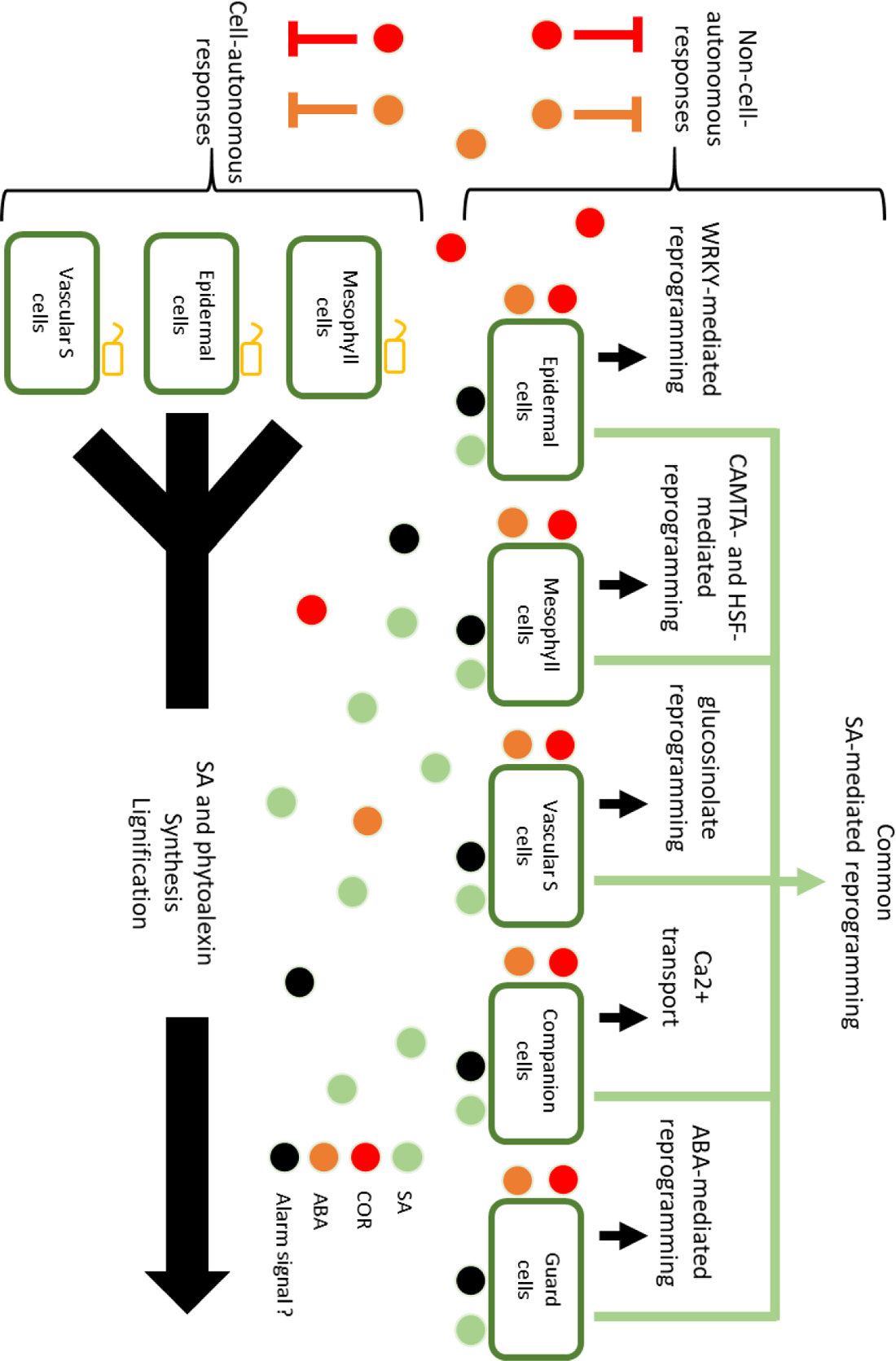
Model of transcriptional defense reprogramming at the cell-type level in response to *Pst*. In a first step, pathogen detection triggers cell-autonomous responses mediated by immune totipotency and resulting in the synthesis of signal alarms like SA, as well as in the synthesis of phytoalexins and lignin acting as barriers against the pathogen. In a second step, alarm signals trigger non-cell-autonomous responses. Features of the cell-type specific non-autonomous responses are indicated by black arrows. The common component of non-cell-autonomous responses, mostly mediated by SA signaling is indicated by a green arrow. Concomitantly coronatine produced by *Pst* and ABA antagonize autonomous and non-autonomous responses, limiting cell evolutions towards immune populations.

*Are all cell types immune-totipotent?* Based on the list of marker genes in the Seurat clustering as well as on the gene modules of Monocle3, we could identify and characterize cell-type defense responses, showing that all cell types don’t respond identically to the pathogen and that they can specialize in peculiar defense processes. Remarkably we could associate the distinct transcriptional reprogramming with specific sets of regulators, indicating that transcription factors known to be involved in the response to *Pst* (e.g. WRKYs, CAMTAs, and ANACs) may act prominently in selective cell populations. Similarly, we found that the expression profiles of key immune receptors can vary considerably between different cell clusters, suggesting that all cell types are not equal in their ability to sense the pathogen, and that some might be more proficient than others in specific kinds of immunity. In parallel, we observed that the different cell types can also undergo common transcriptional reprogramming. The construction of cell trajectories even supports the possibility that, in response to *Pst*, epidermal, mesophyll and some vascular cells can converge towards a unique cell fate mainly characterized by resistance functions like lignification, detoxification or phytoalexin synthesis.

From this, emerges a scenario in which the defense responses of the different cell types (at least for mesophyll, vascular S and epidermal cells) could be divided into two components. The first one would be shared, resistance-centered, and could therefore substantiate the notion of immune-totipotency. The second one would be cell-type specific, allowing the implementation and coordination of several distinct protective measures. If so, the factors that determine the respective influence of these two response components still need to be elucidated.

*How do susceptibility and immunity take place at the single cell level?* In a recent scRNA-seq study, it was proposed that, upon *Pst* infection, cells would evolve from early immune responses to late susceptible responses ^53^. Our own data do not really retrieve this pattern, showing rather a concurrent activation of immune and susceptible processes in all the cells. A simple explanation for this discrepancy could be differences in the experimental set up between the two studies. Indeed, we performed our analyses earlier than in Zhu et al., 2022 ^53^ (16 hpi vs 24 hpi), it is therefore possible that our data lack the strong and late susceptible responses. An alternative could be differences in the degree of plant resistance or pathogen virulence between the two experiments. Our interpretation is that immune and susceptible processes occur synchronously in all cells, and that it is this competition that drives cell evolution towards specific responsive populations. In this regard, the outcome of a pathogen infection might be more reliably predicted by the relative sizes or proportions of these responsive populations rather than by the expression features of some susceptible or immune genes. According to this, the absence in Zhu et al., 2022 ^53^ of clusters equivalent to C5 and C13 could indicate that the plants of this study are more susceptible than ours. In the future it would be greatly interesting to test the hypothesis mentioned above, with scRNA-seq analyses of leaves inoculated with different strains of *Pst* that are more or less virulent.

*What are the cell-autonomous and non-cell-autonomous responses to Pst?* To explain the divergence of the mesophyll, epidermal and vascular S cells into two separate populations, an exciting possibility could be that it reflects the difference between cell-autonomous and non-cell-autonomous responses. In this case, we would have on one hand the C13 and C5 populations, composed of cells coming from different types, not only committed to stop the pathogens, but also essential for the synthesis of proximal or distal alarm signals, like SA. This would correspond to the autonomous responses of cells directly in contact with the pathogens, i.e. at the closest of the battlefronts, and characterized by the immune-totipotent component of the defense responses. On the other hand, we would have cell-type characteristic responses, triggered by intercellular communication, and allowing plant cells to mount prophylactic measures in anticipation of upcoming pathogen attacks. This would correspond to the non-cell-autonomous responses, mediated by both common and distinct components of the defense transcriptional reprogramming. If such a model is true, the nature of the intercellular signals as well as the efficiency of the cell-type responses to impede infection remain to be further clarified.

## Supporting information

FiguresS1-S10S12

## Acknowledgements

This study was supported by INRAE funding. The POPS platform benefits from the support of Saclay Plant Sciences-SPS (ANR-17-EUR-0007). The authors thank the IPS2 bioinformatic platform (Marion Verdenaud and Frédéric Desprez, https://ips2.u-psud.fr/en/presentation/services-d-appui-a-la-recherche/bioinformatic-tray.html) for facilitating installations of the Seurat and Monocle3 packages. The authors also thank Marie Boudsocq (Researcher, CNRS) for technical assistance with protoplast preparation.

## Author contributions

ED and JL designed the study. ED, SP, LST and JL performed the experiments. ED, BB and JL processed the data and discussed the interpretations. JL wrote the manuscript with inputs from ED, BB, JC^3^ and JC^1,2^.

## Supplemental Figure titles and legends

**Figure S1:** UMAP projection of the 18 clusters obtained by PCA.

**Figure S2:** Cluster-specific expression profiles of cell-type marker genes. Size of the circles represent the percentage of cells expressing the gene, while the color represents the transcript levels. The marker genes were selected from the literature (Kim et al., 2021 ^33^; Routaboul et al., 2022 ^34^; Tenorio Berrío et al., 2022 ^35^). GC, CC, EC, HC, MC and SC stand for guard cells, companion cells, epidermal cells, hydathode cells, mesophyll cells and vascular S cells respectively.

**Figure S3:** Cell repartition of the three biological replicates in the total population and in each cluster. In the total 11206 cells of the analysis, 2861 (25,5%) come from replicate 1, 5397 (48%) from replicate 2 and 2948 (26,5%) from replicate 3. The clusters showing a repartition consistent with the repartition in the total number of cells are highlighted in red.

**Figure S4:** UMAP projection and cell composition between replicates of the healthy mesophyll clusters C0, C4, C10 and C11.

**Figure S5:** Cell-specific expression profiles of 6 aquaporin genes in the UMAP projection.

**Figure S6:** Cell-specific expression profiles of C9 marker genes in the UMAP projection. Genes were related to Et signaling and responses to biotic stress.

**Figure S7:** Cell-specific expression profiles of 4 common C9 and C1 marker genes in the UMAP projection.

**Figure S8:** Cluster-specific expression profiles of genes involved in camalexin synthesis.

**Figure S9:** Cluster-specific expression profiles of *NLR* genes. 207 *NLR* genes (Meyers et al., 2003 ^51^) are showed.

**Figure S10:** Cluster-specific expression profiles of *PRR* genes. 236 genes from the *LRR I-XIII* and *LysM* families (Shiu & Bleecker, 2001 ^52^) are shown.

**Figure S11:** Expression scores of the gene modules. (A) Cluster-specific expression scores of the 71 gene modules. (B) Cell-specific expression scores of the 71 gene modules (1 per sheet).

**Figure S12:** Cell-specific expression profiles of genes involved in SA synthesis and signaling.

## Supplemental table titles and legends

**Table S1:** Number of cells and marker genes associated to each cluster.

**Table S2:** List of marker genes for each cluster. The list was generated with Seurat. Avg_logFC (log 2) represents the upregulation of the gene in this cluster compared to the mean of all the other clusters. Pct.1 and pct.2 represent the percentage of cells expressing the gene in this cluster and in all the other clusters, respectively. The Gene Ontology (GO) analysis of biological processes was performed with Panther on the basis of all marker genes for each cluster.

**Table S3:** Stringent list of marker genes for each cluster. Each tab corresponds to a cell population and includes the list of marker genes with their upregulation level and their annotations from TAIR 10. The lines in pink correspond to cell-type marker genes found in Kim et al., 2021 ^33^. The enrichments in GO biological processes indicated below were generated using only the specific responsive genes (not in pink).

**Table S4:** Composition of the 71 gene modules generated by Monocle3. When necessary, GO enrichments or gene descriptions are given on the right of each module. For module 23 the genes in pink correspond to the genes in common with the markers of the healthy companion cells (Kim et al., 2021 ^33^).

