## Supplementary material for "Cell specialization and coordination in *Arabidopsis* leaves upon pathogenic attack revealed by scRNA-seq": FiguresS1-S10S12

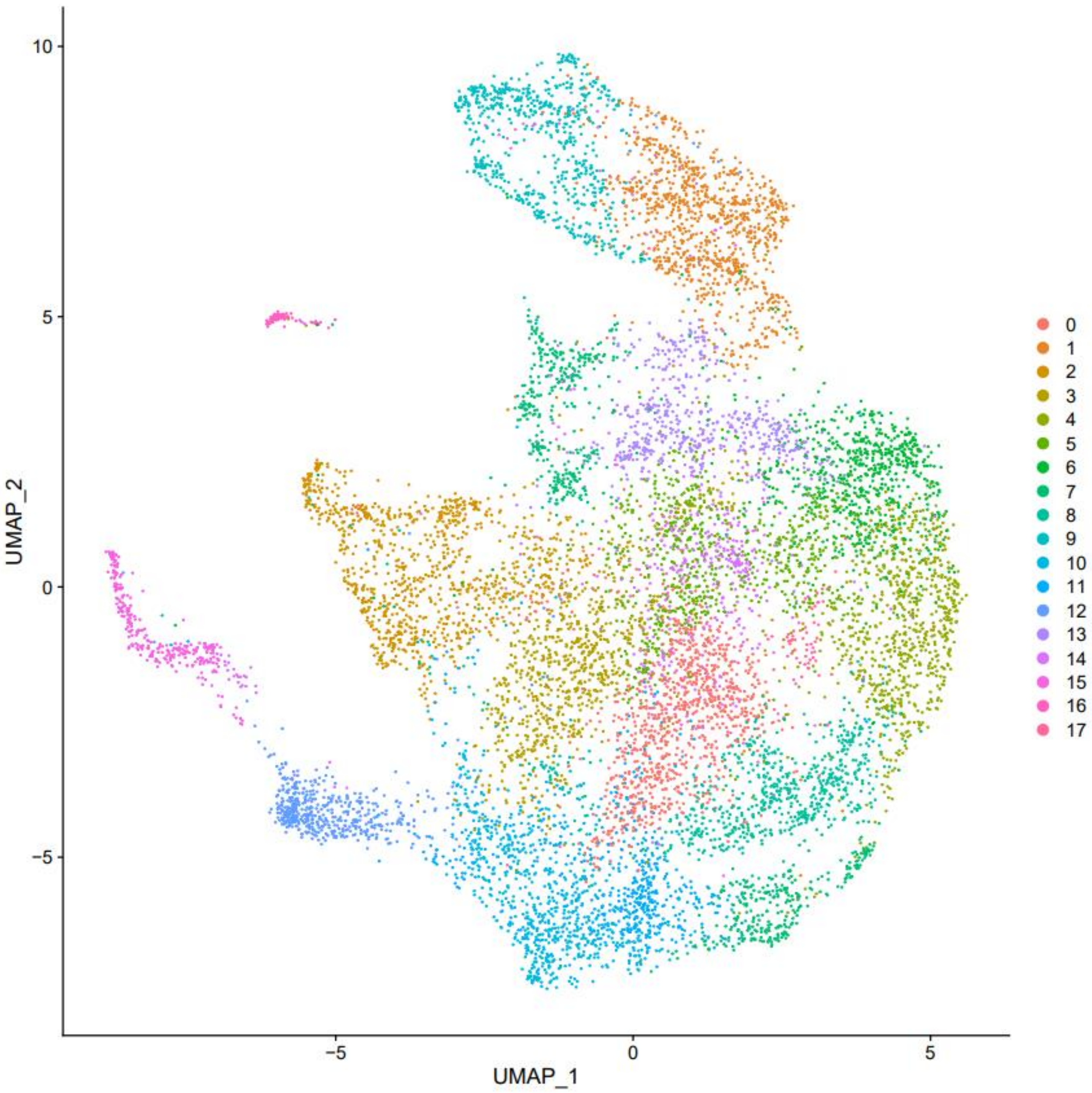


**Figure S1: UMAP projection of the 18 clusters obtained by PCA.**

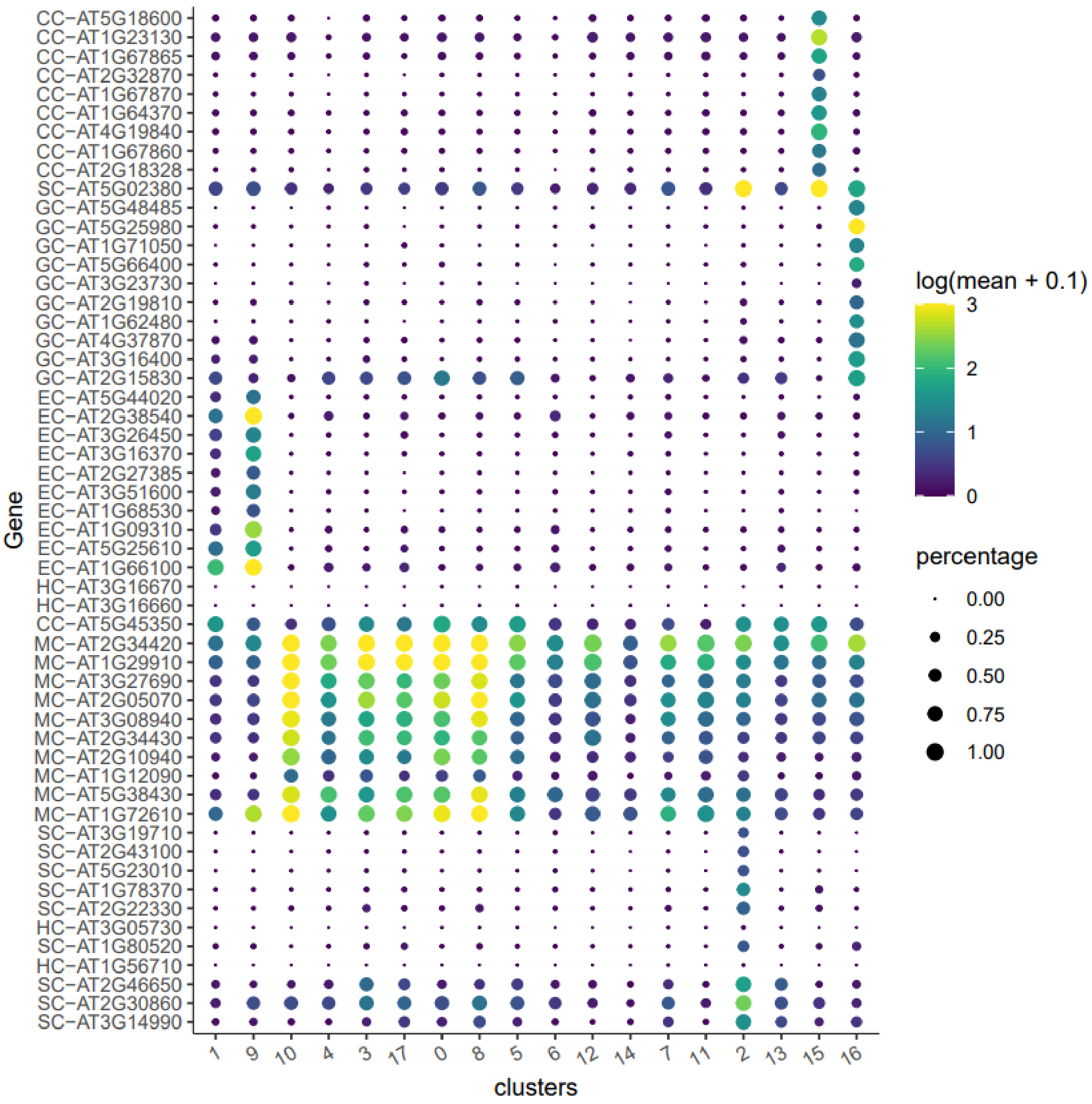


**Figure S2: Cluster-specific expression profiles of cell-type marker genes.** Size of the circles represent the percentage of cells expressing the gene, while the color represents the transcript levels. The marker genes were selected from the litterature (Kim et al., 2021; Routaboul et al., 2022; Tenorio Berrío et al., 2022). GC, CC, EC, HC, MC and SC stand for guard cells, companion cells, epidermal cells, hydathode cells, mesophyl cells and vascular S cells respectively.

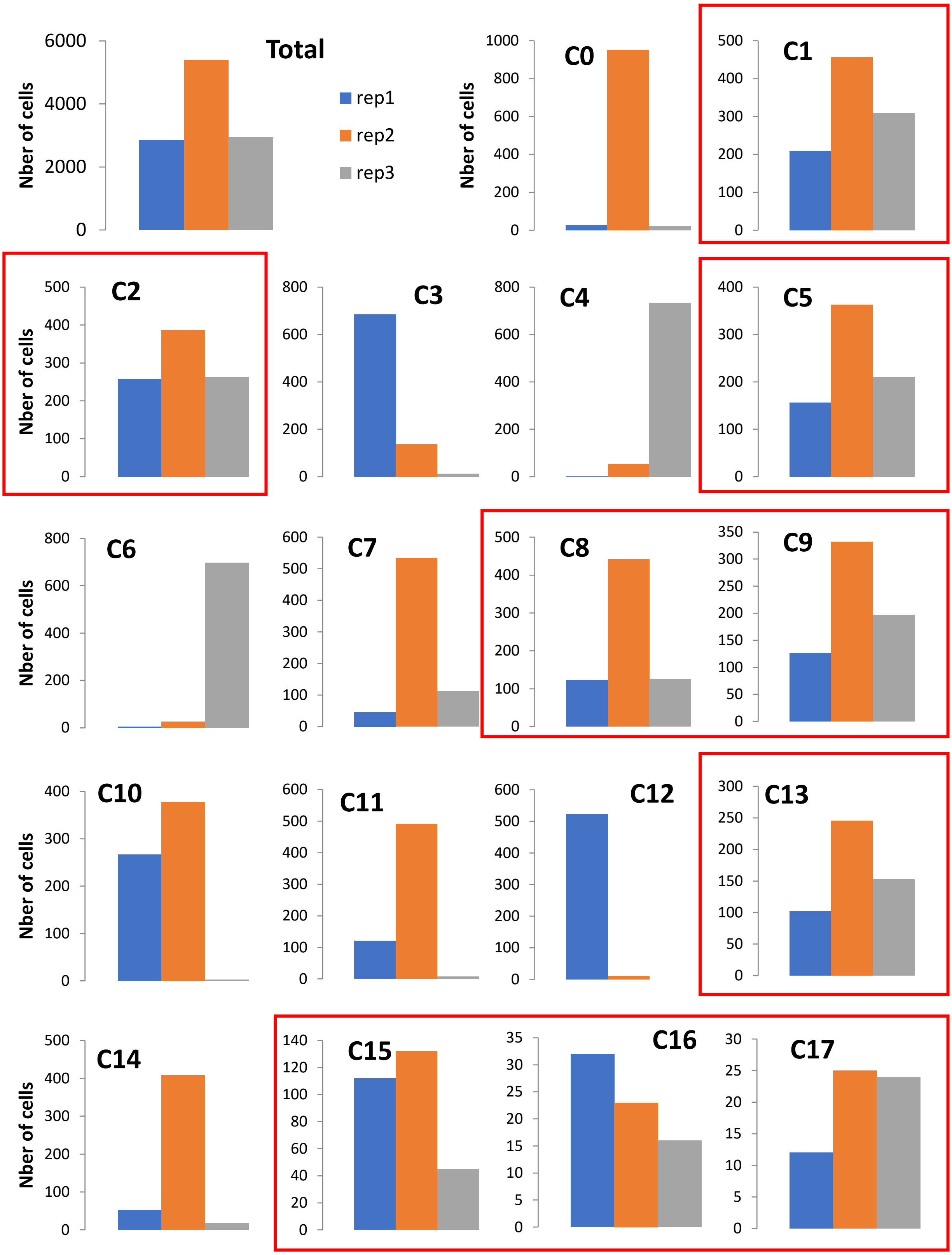


**Figure S3: Cell repartition of the three biological replicates in the total population and in each cluster**. In the total 11206 cells of the analysis, 2861 (25,5%) come from replicate 1, 5397 (48%) from replicate 2 and 2948 (26,5%) from replicate 3. The clusters showing a repartition consistent with the repartition in the total number of cells are highlighted in red.

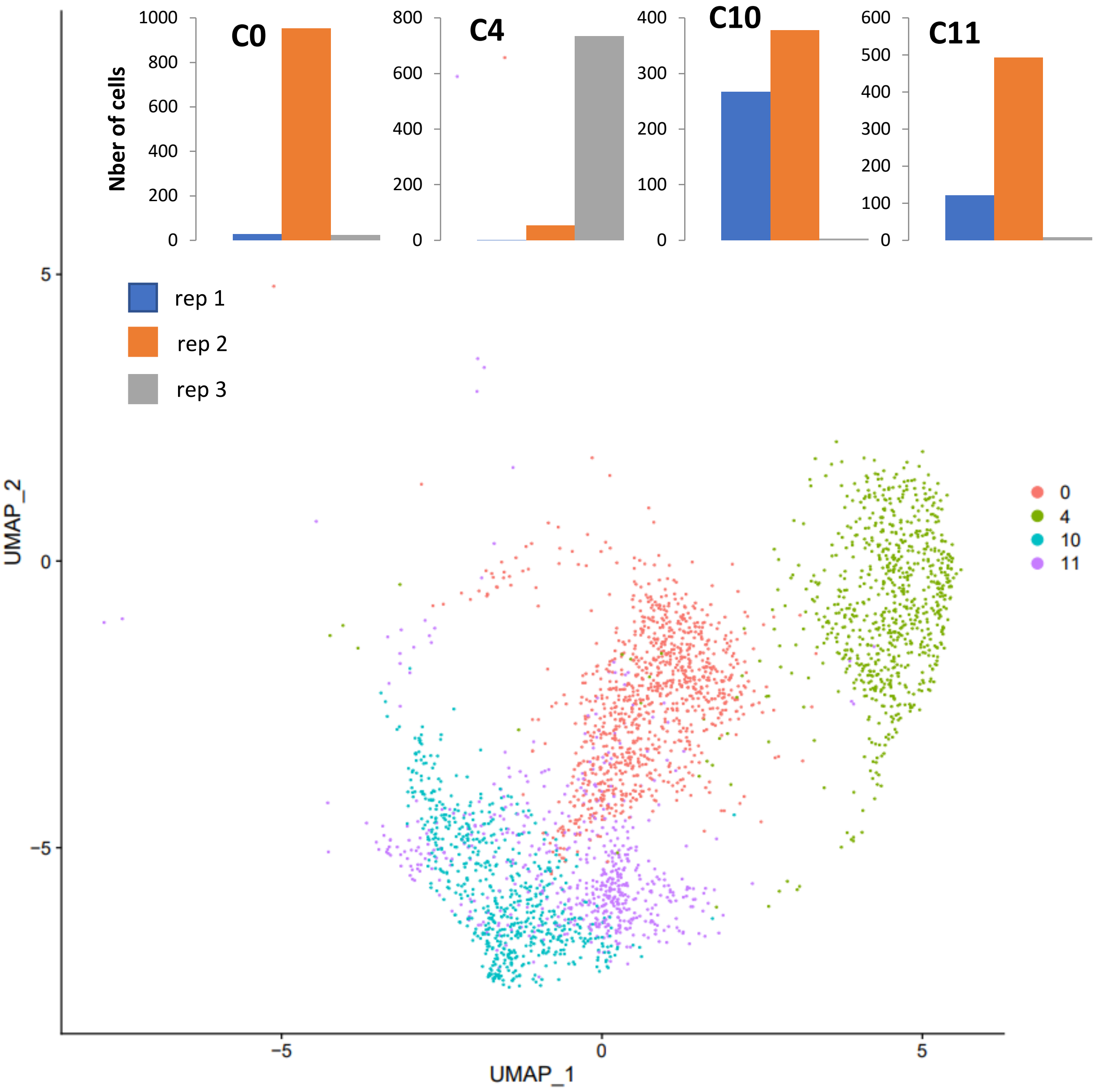


**Figure S4: UMAP projection and cell composition between replicates of the healthy mesophyl clusters C0, C4, C10 and C11.**

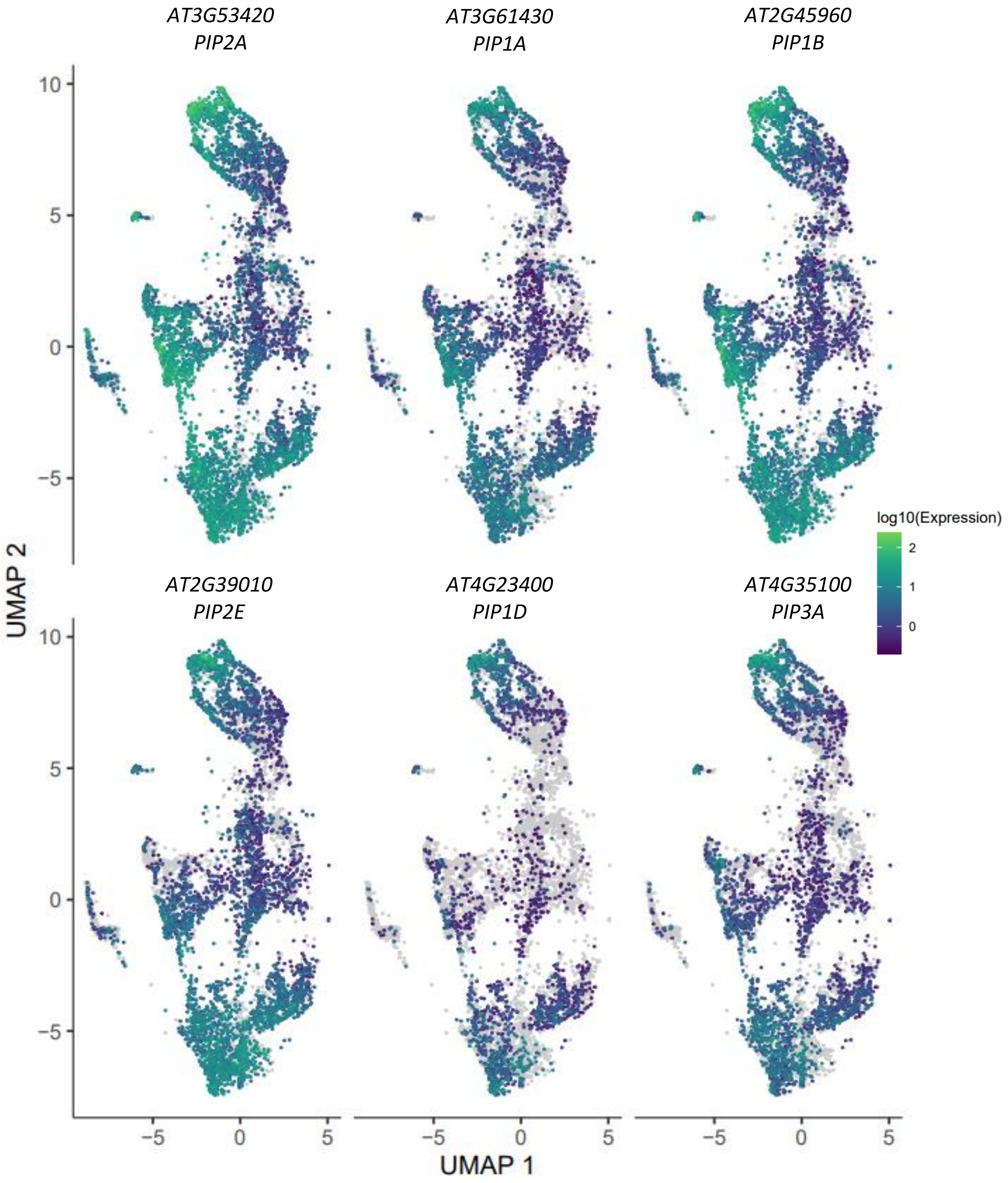


**Figure S5: Cell-specific expression profiles of 6 aquaporins in the UMAP projection.**

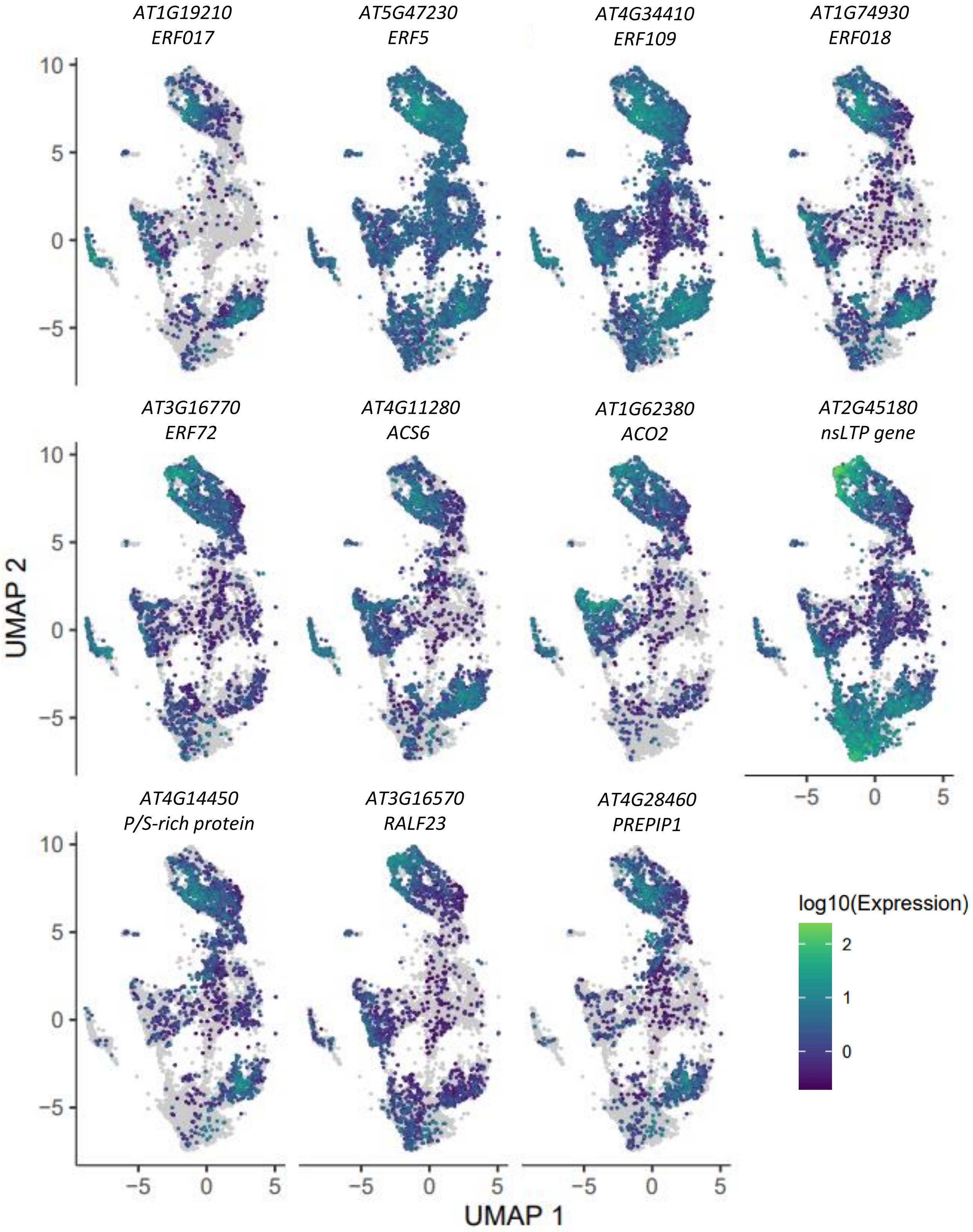


**Figure S6: Cell-specific expression profiles of C9 marker genes in the UMAP projection.** Genes were related to Et signaling and responses to biotic stress.

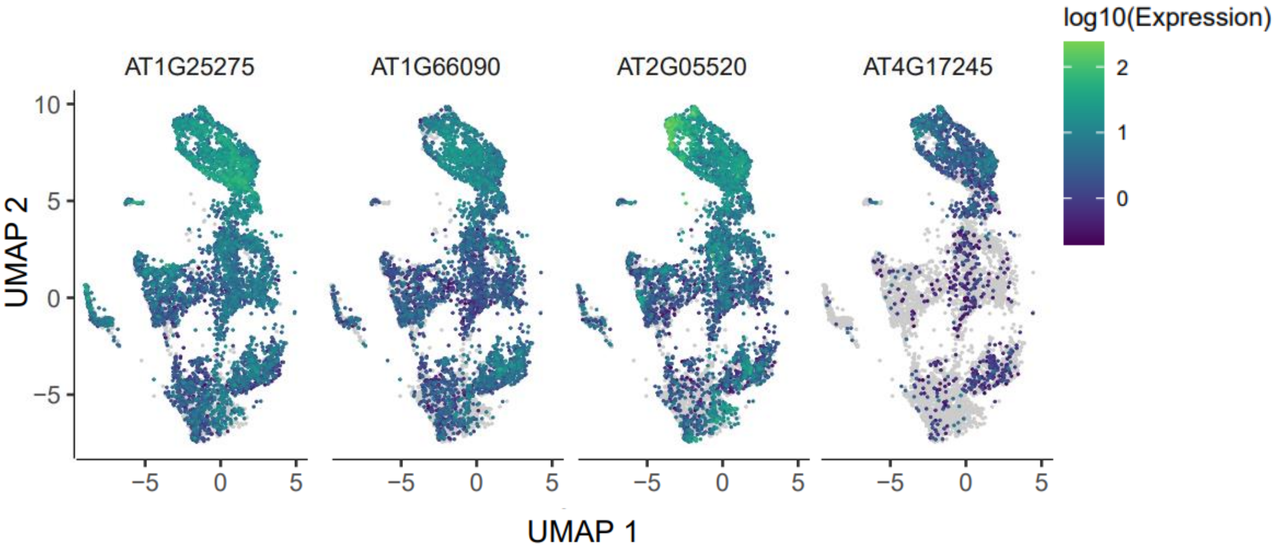


**Figure S7: Cell-specific expression profiles of common C9 and C1 marker genes in the UMAP projection.**

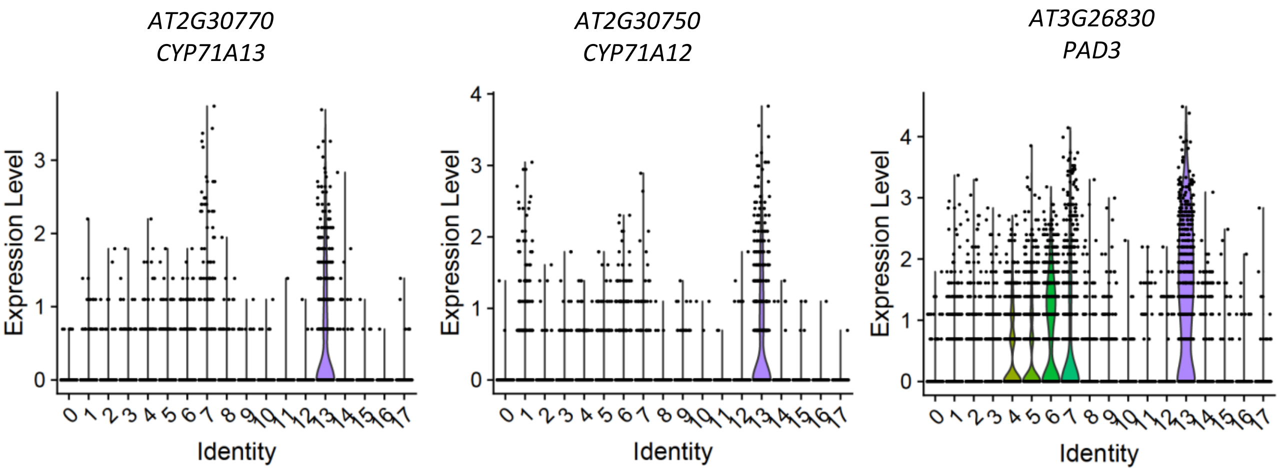


**Figure S8: Cluster-specific expression profiles of genes involved in camalexin synthesis.**

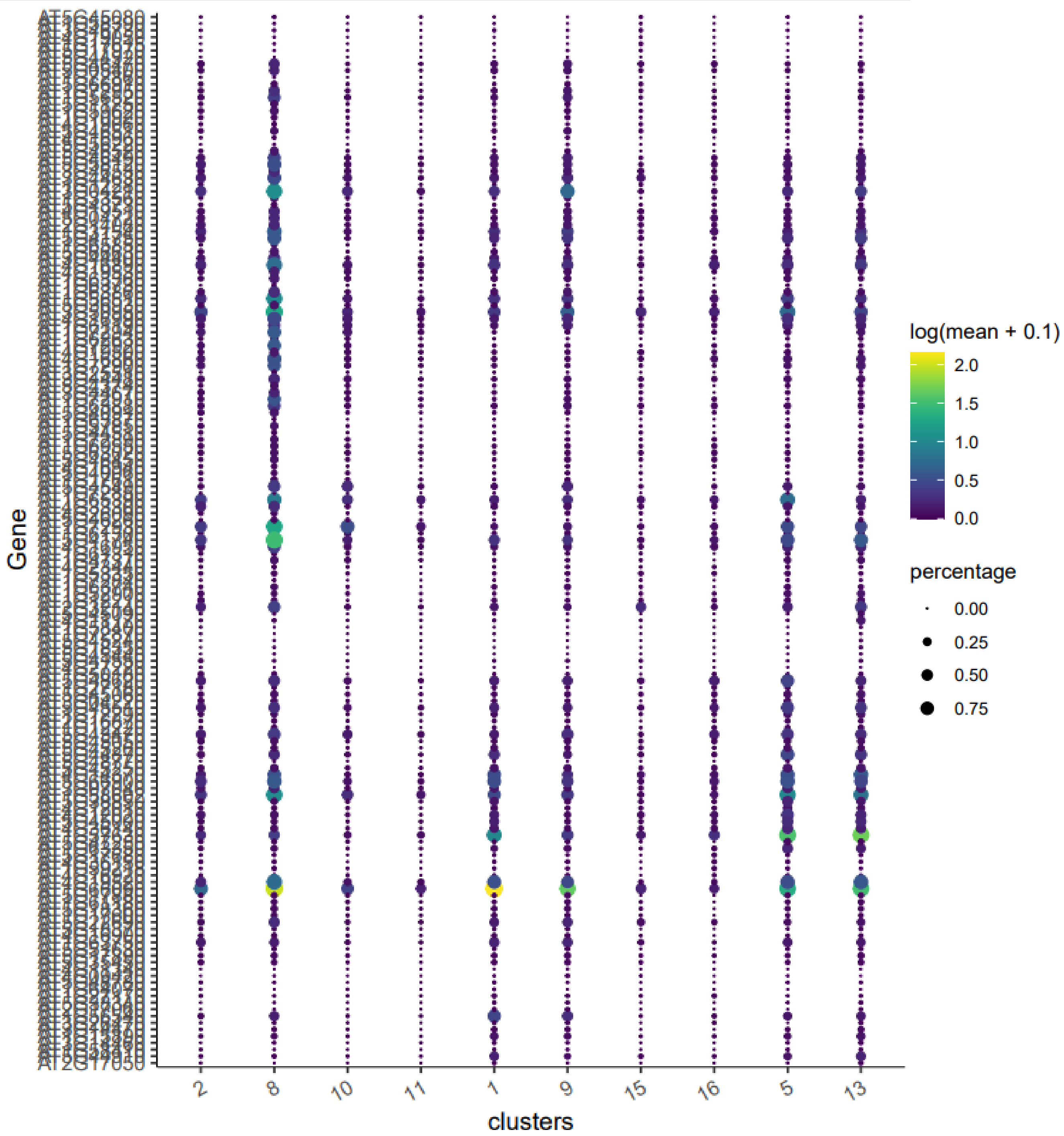


**Figure S9: Cluster-specific expression profiles of *NLR* genes.** 207 *NLR* genes (Meyers et al., 2003) are showed.

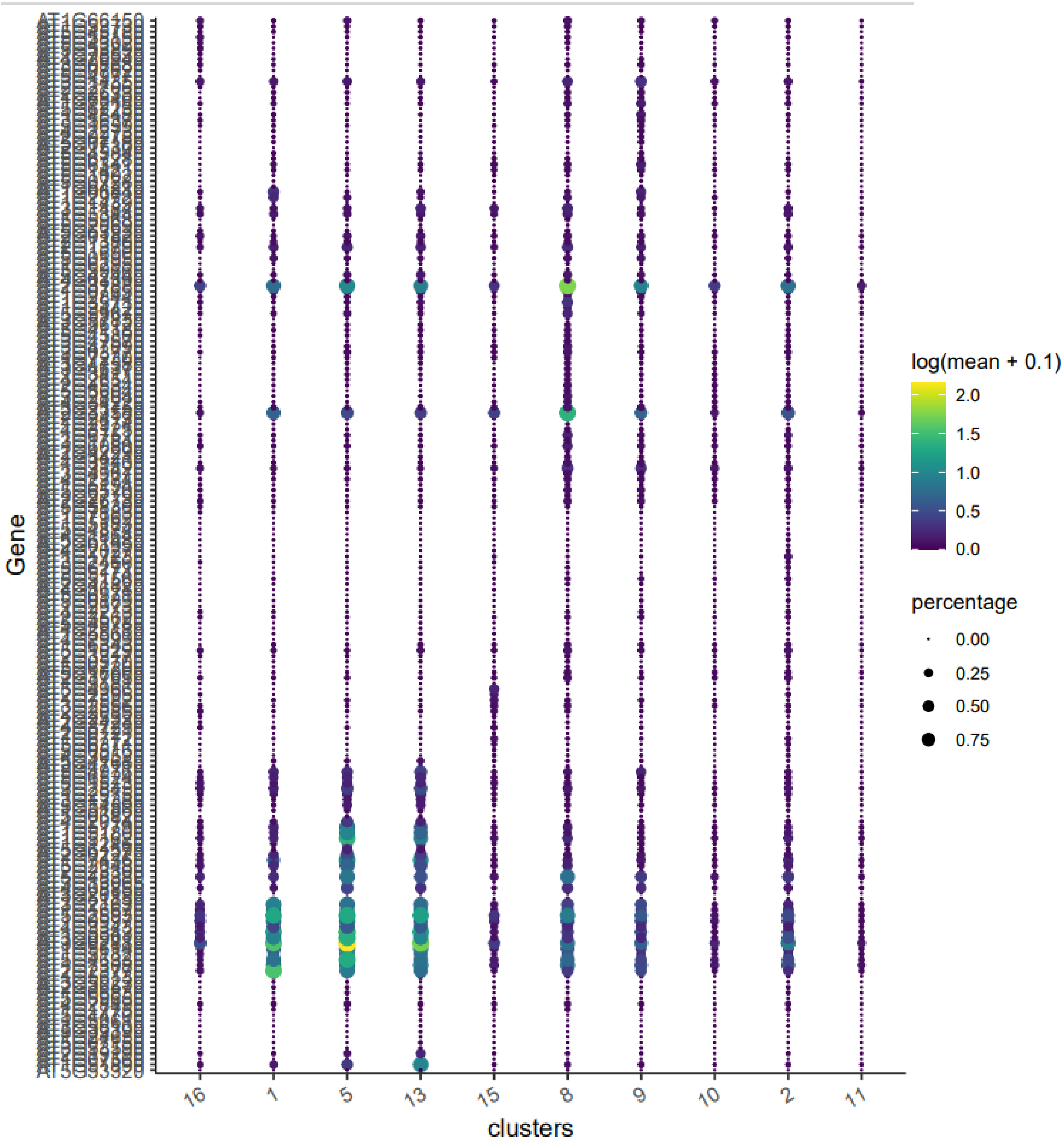


**Figure S10: Cluster-specific expression profiles of *PRR* genes.** 236 genes from the LRR I-XIII and LysM families (Shiu & Bleecker, 2001) are shown**.**

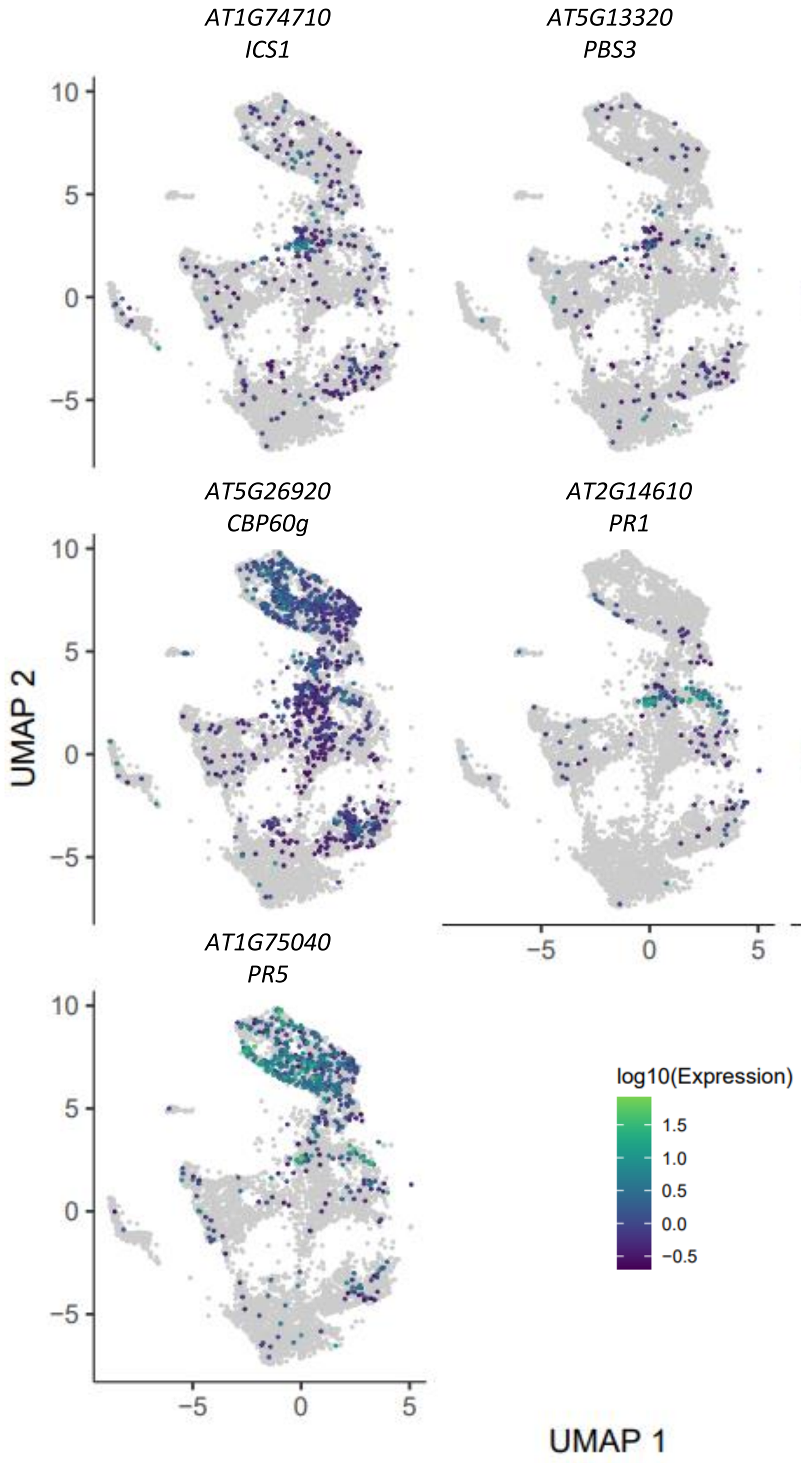


**Figure S12: Cell-specific expression profiles of genes involved in SA synthesis and signaling.**
