## Supplementary material for "Cell specialization and coordination in *Arabidopsis* leaves upon pathogenic attack revealed by scRNA-seq": FigureS11

**Figure S11: Expression scores of the gene modules.** (A) Cluster-specific expression scores of the 71 gene modules. (B) Cell-specific expression scores of the 71 gene modules (1 per sheet).

A

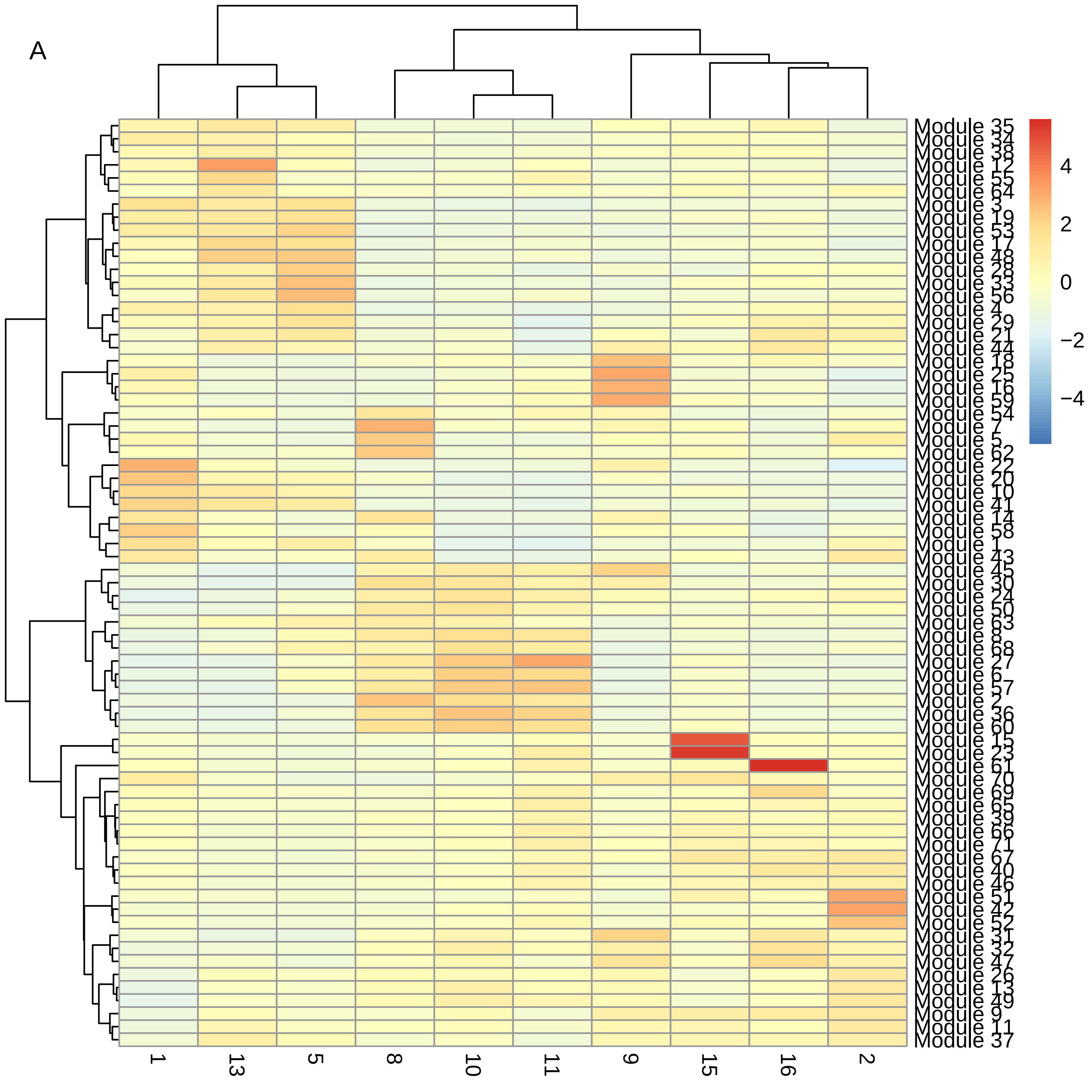

Module 35
Module 34
Module 38
Module 12
Module 55
Module 64
Module 3
Module 19
Module 53
Module 17
Module 48
Module 28
Module 33
Module 56
Module 4
Module 29
Module 21
Module 44
Module 18
Module 25
Module 16
Module 59
Module 54
Module 7
Module 5
Module 62
Module 22
Module 20
Module 10
Module 41
Module 14
Module 58
Module 1
Module 43
Module 45
Module 30
Module 24
Module 50
Module 63
Module 8
Module 68
Module 27
Module 6
Module 57
Module 2
Module 36
Module 60
Module 15
Module 23
Module 61
Module 70
Module 69
Module 65
Module 39
Module 66
Module 71
Module 67
Module 40
Module 46
Module 51
Module 42
Module 52
Module 31
Module 32
Module 47
Module 26
Module 13
Module 49
Module 9
Module 11
Module 37
1
13
5
8
10
11
9
15
16
2
4
2
0
−2
−4

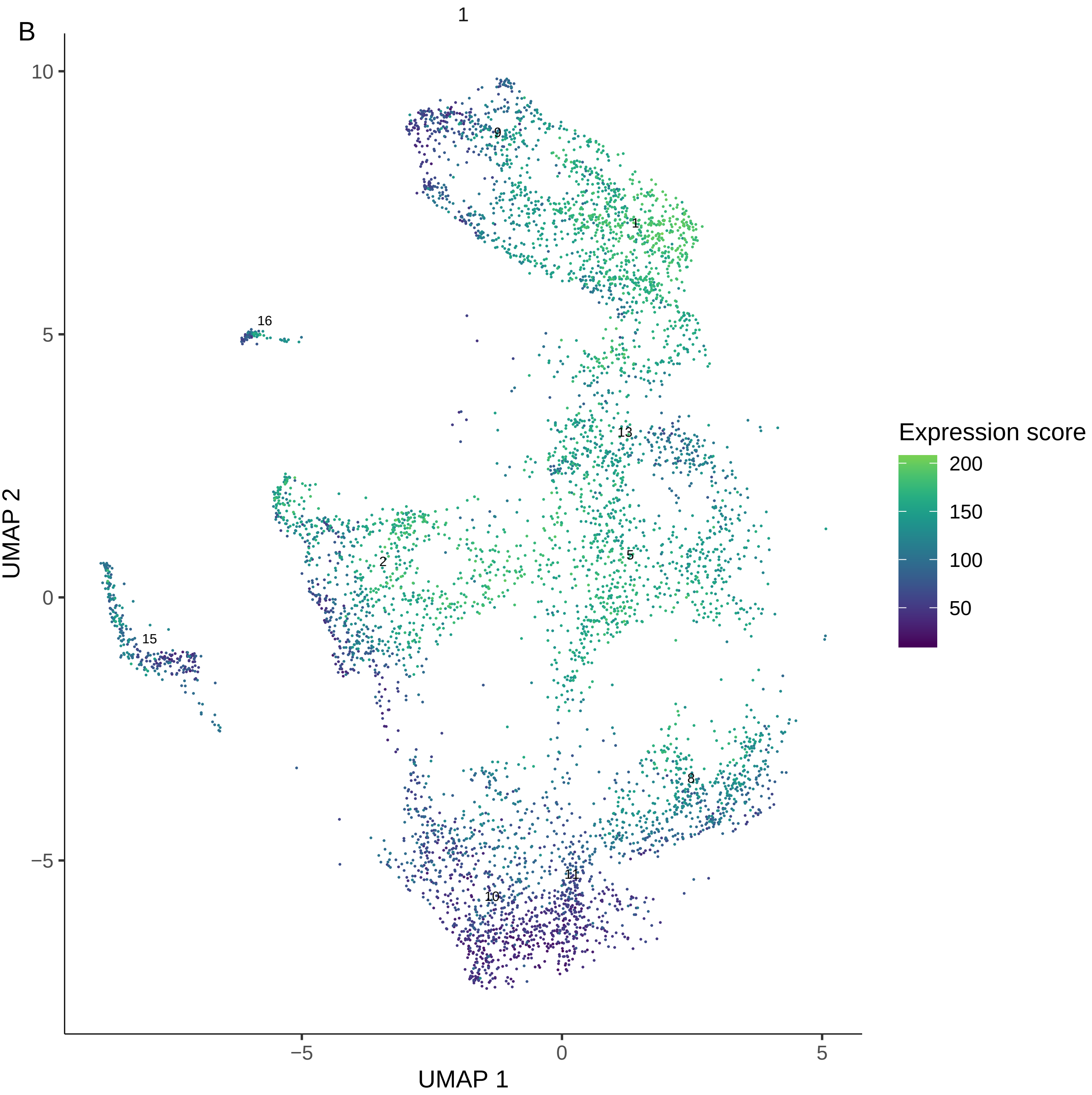

B
1
10
5
0
−5
−5
0
5
UMAP 1
UMAP 2
9
1
16
13
2
5
15
8
11
10
Expression score
200
150
100
50

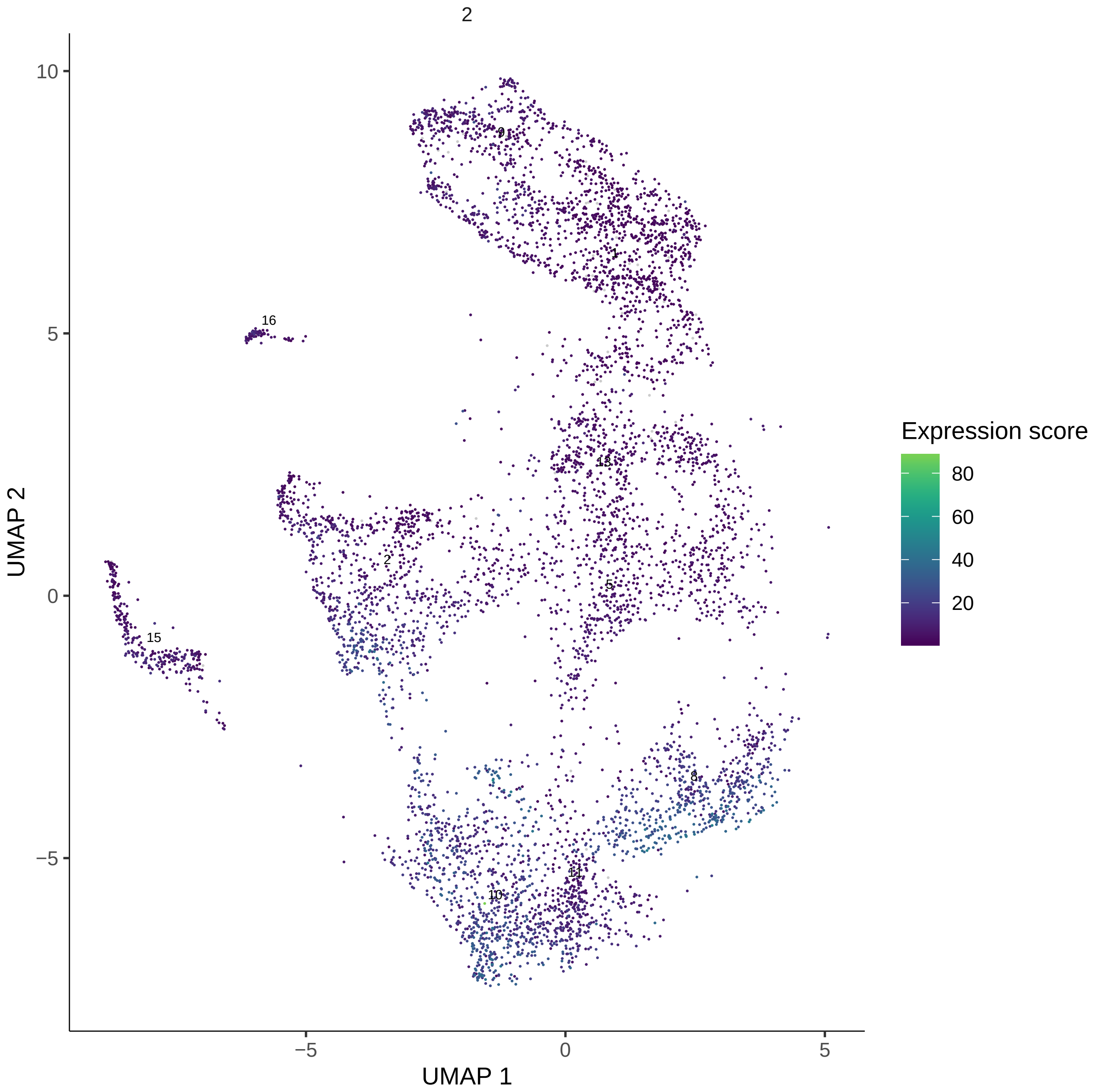
2
UMAP 1
UMAP 2
Expression score
80
60
40
20
10
5
0
−5
−5
0
5
9
1
16
13
2
5
15
8
11
10

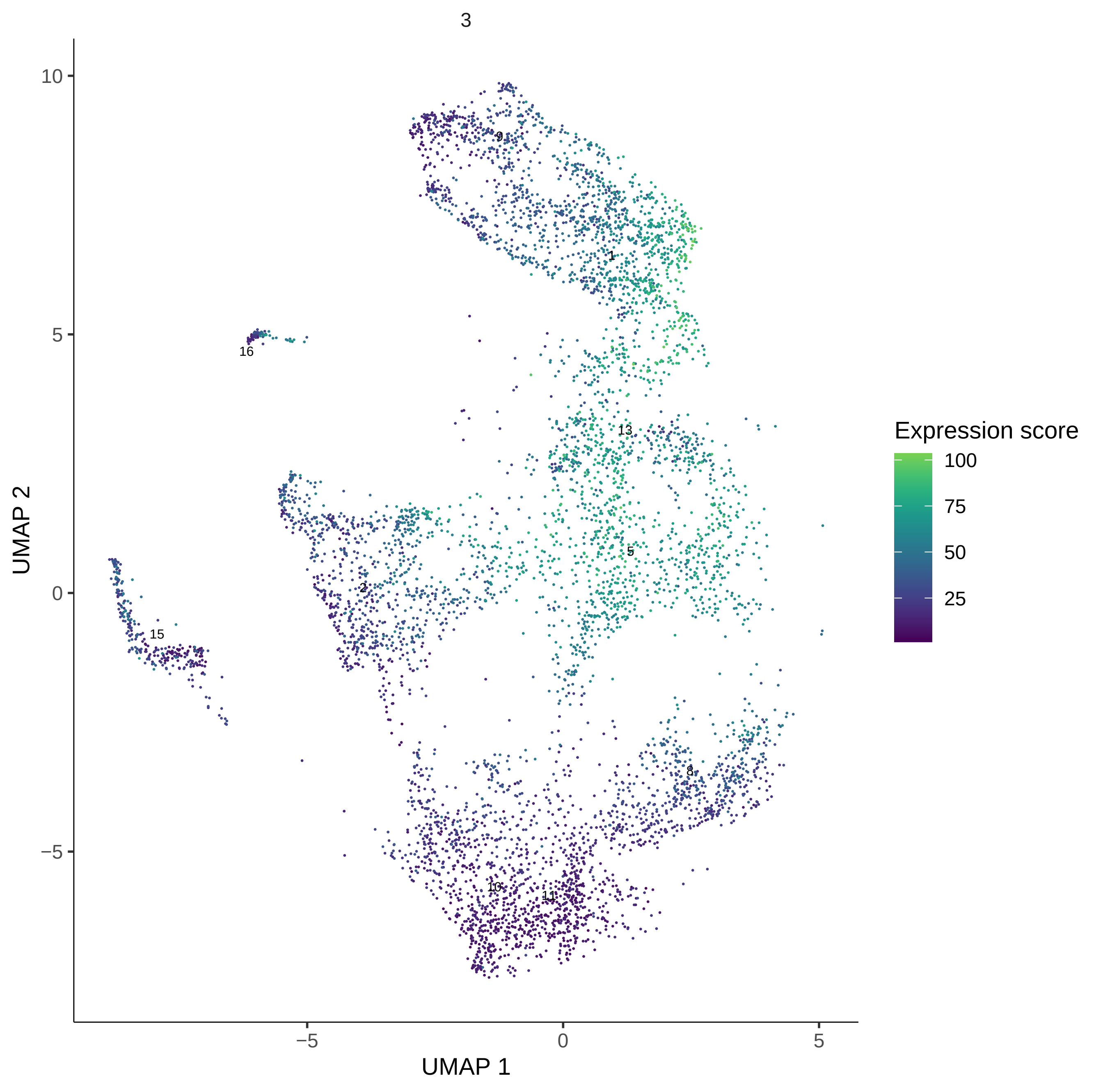
3
UMAP 1
UMAP 2
−5
0
5
10
Expression score
100
75
50
25
9
1
16
13
5
2
15
8
10
11

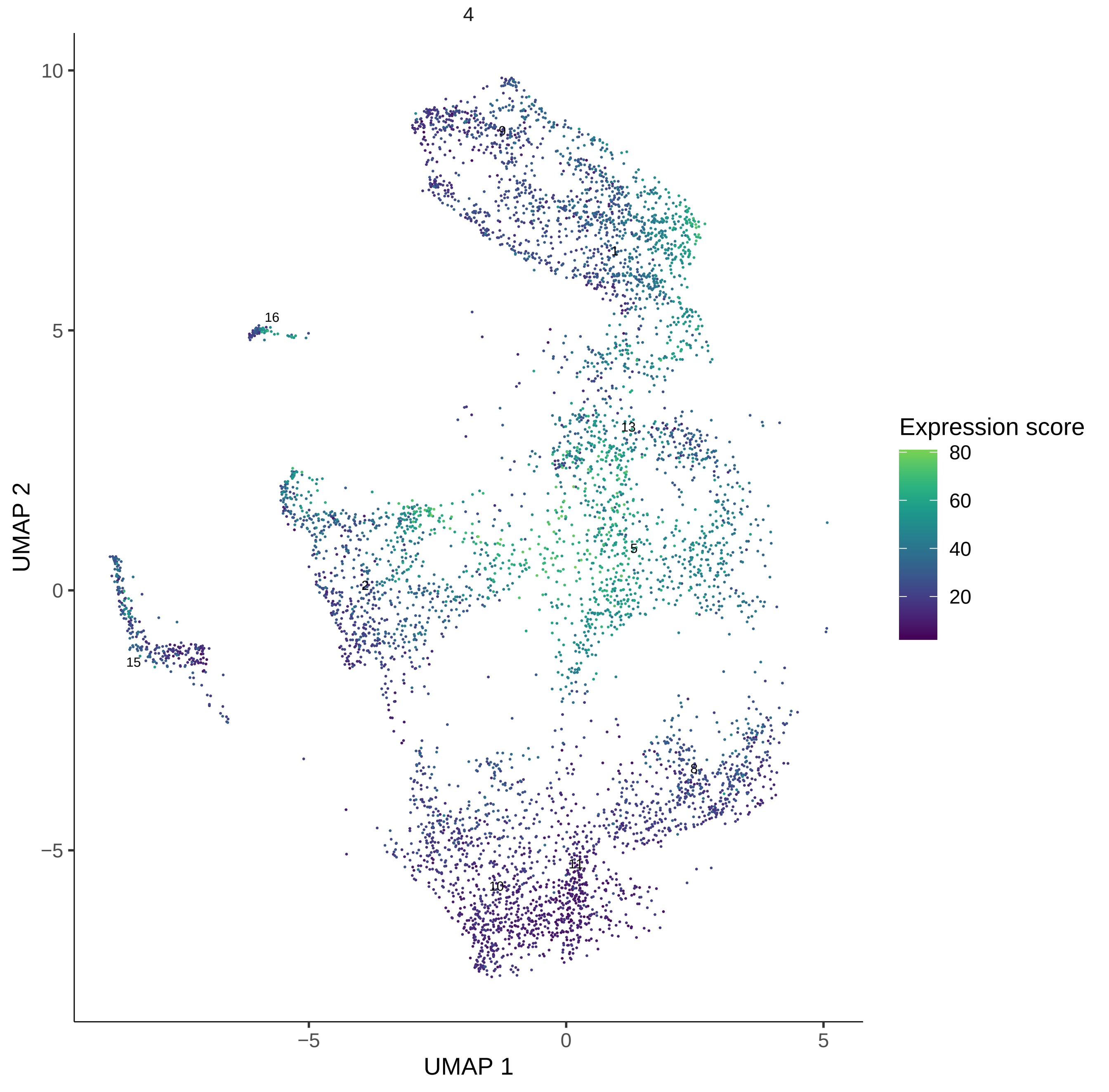
4
10
5
0
−5
−5
0
5
UMAP 1
UMAP 2
9
1
16
13
5
2
15
8
11
10
Expression score
80
60
40
20

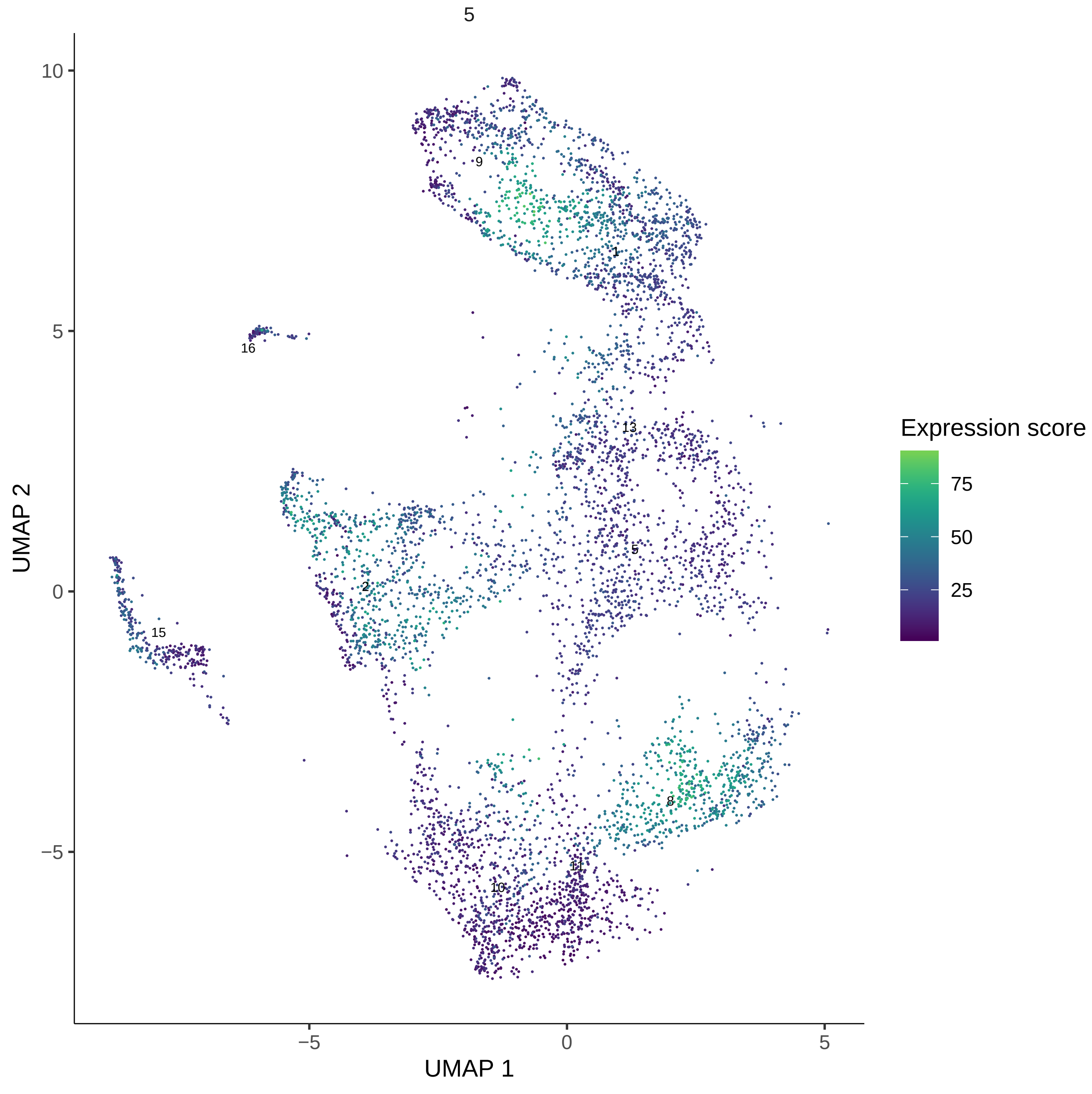
5
10
5
0
−5
UMAP 2
−5
0
5
UMAP 1
9
1
16
13
5
2
15
8
11
10
Expression score
75
50
25

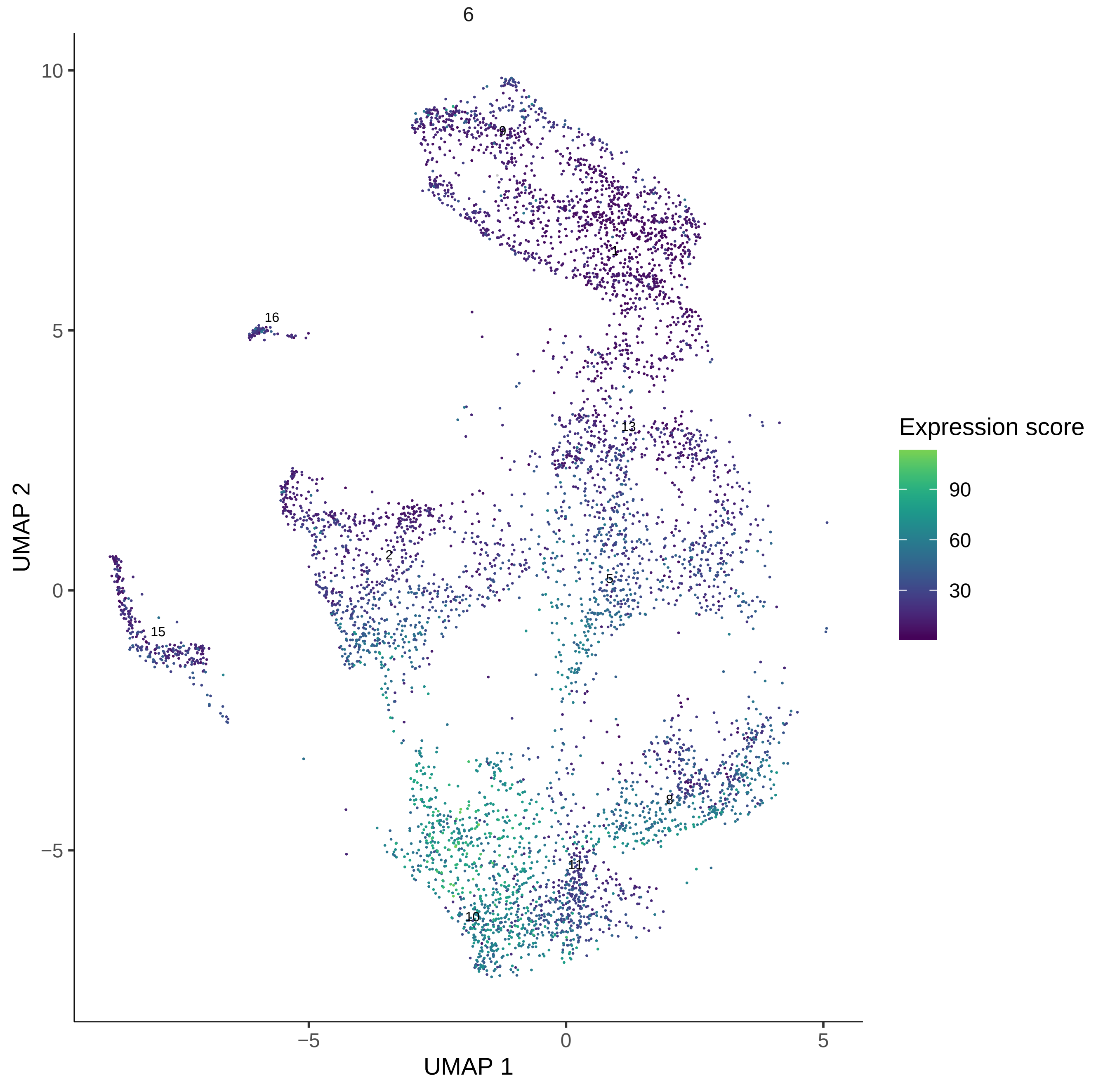
6
10
5
0
−5
−5
0
5
UMAP 1
UMAP 2
16
9
1
13
2
5
15
8
11
10
Expression score
90
60
30

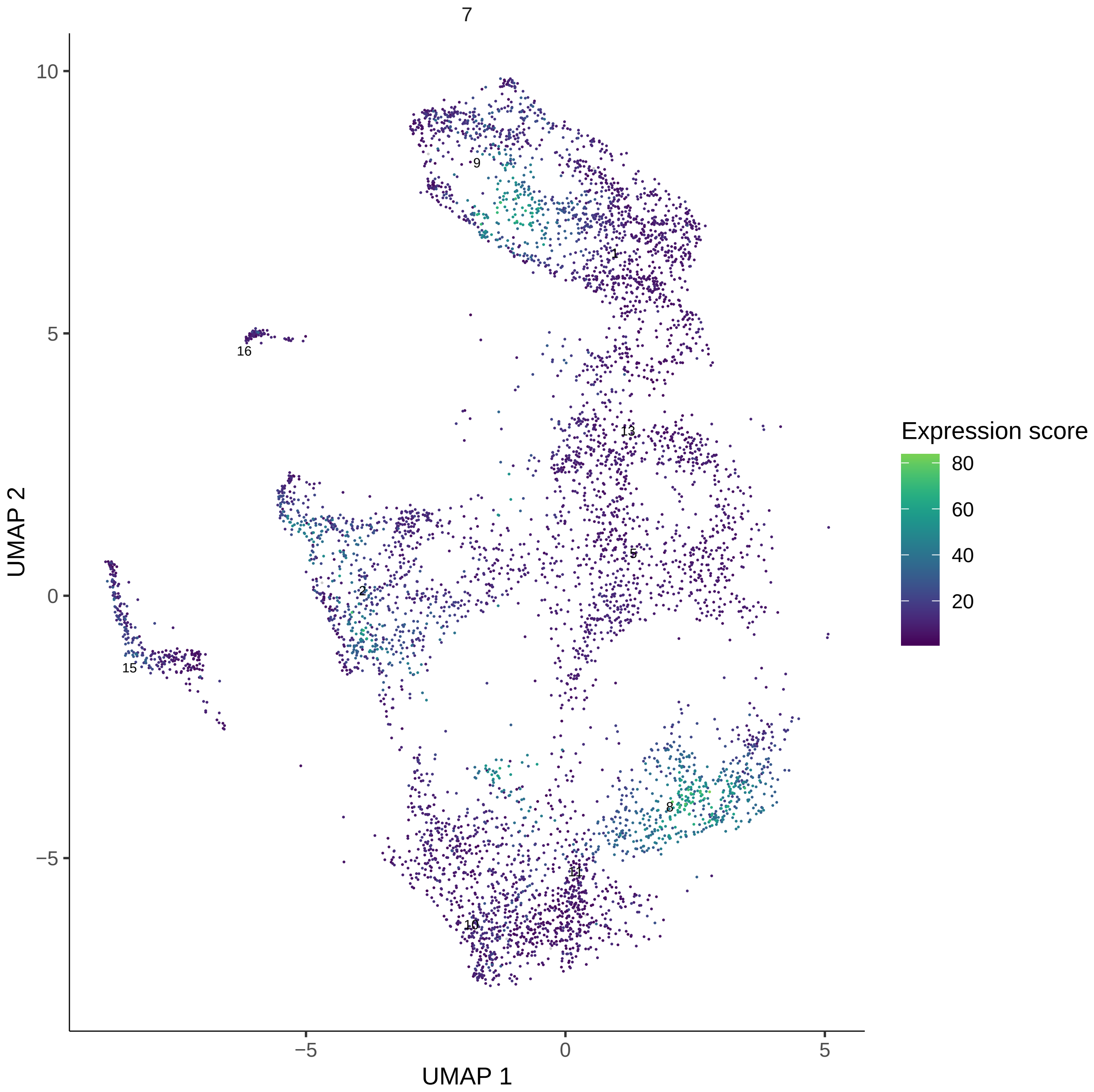
7
UMAP 1
UMAP 2
−5
0
5
10
Expression score
80
60
40
20
9
1
16
13
5
2
15
8
11
10

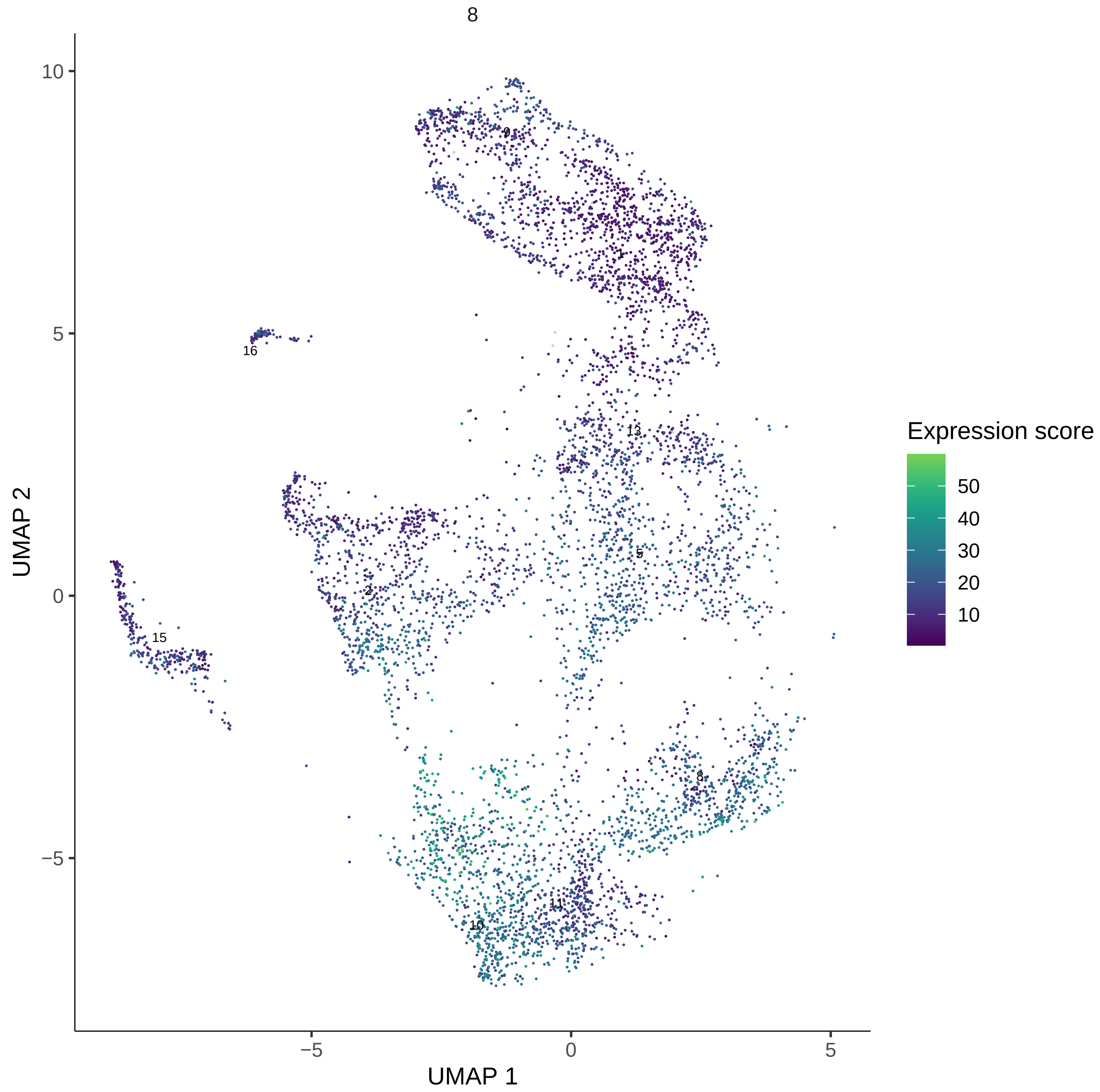

8
UMAP 2
UMAP 1
10
5
0
−5
−5
0
5
9
1
16
13
5
2
15
8
11
10
Expression score
50
40
30
20
10

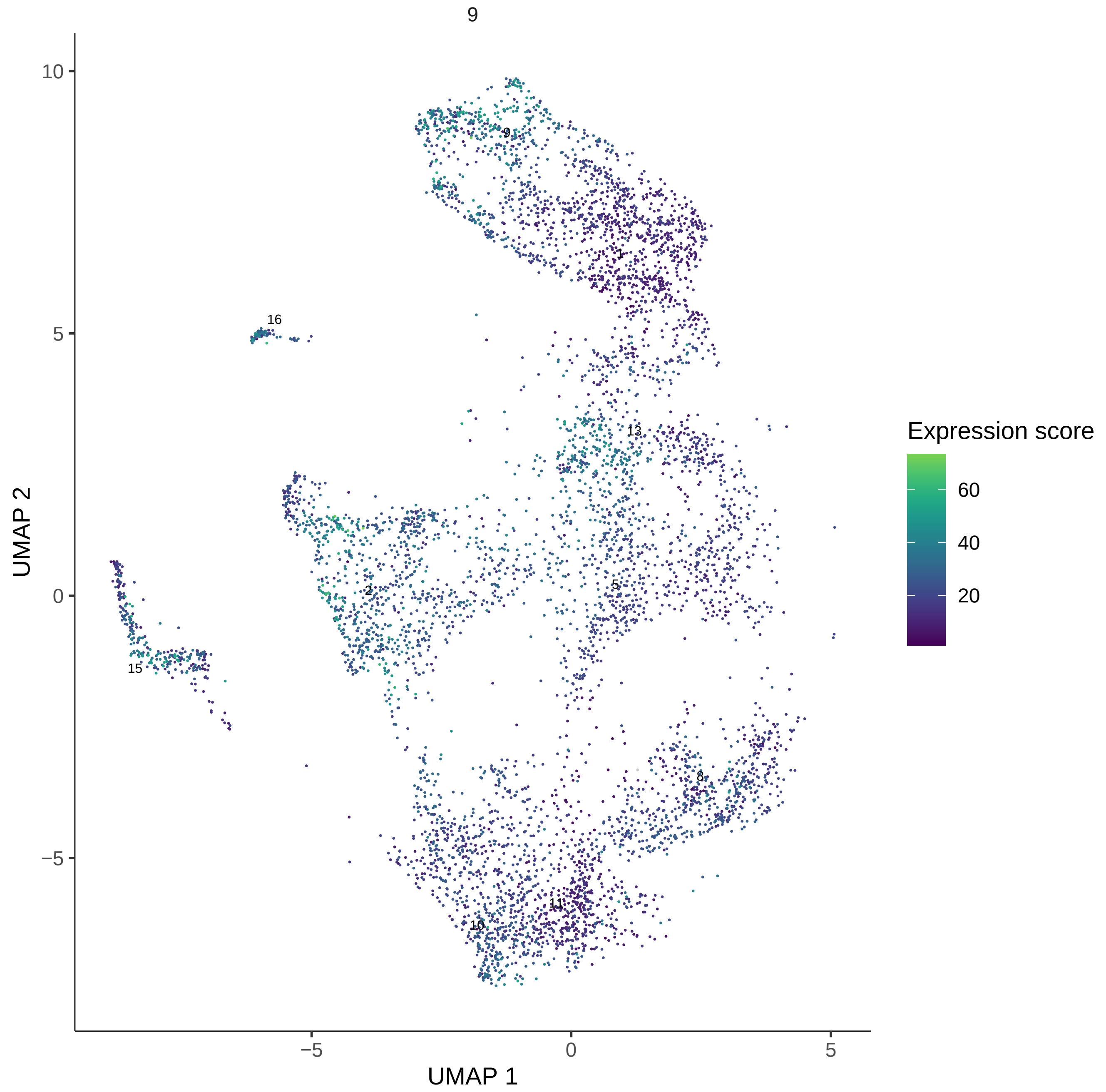

9
UMAP 1
UMAP 2
Expression score
60
40
20
10
5
0
−5
16
15
2
13
1
8
11
10

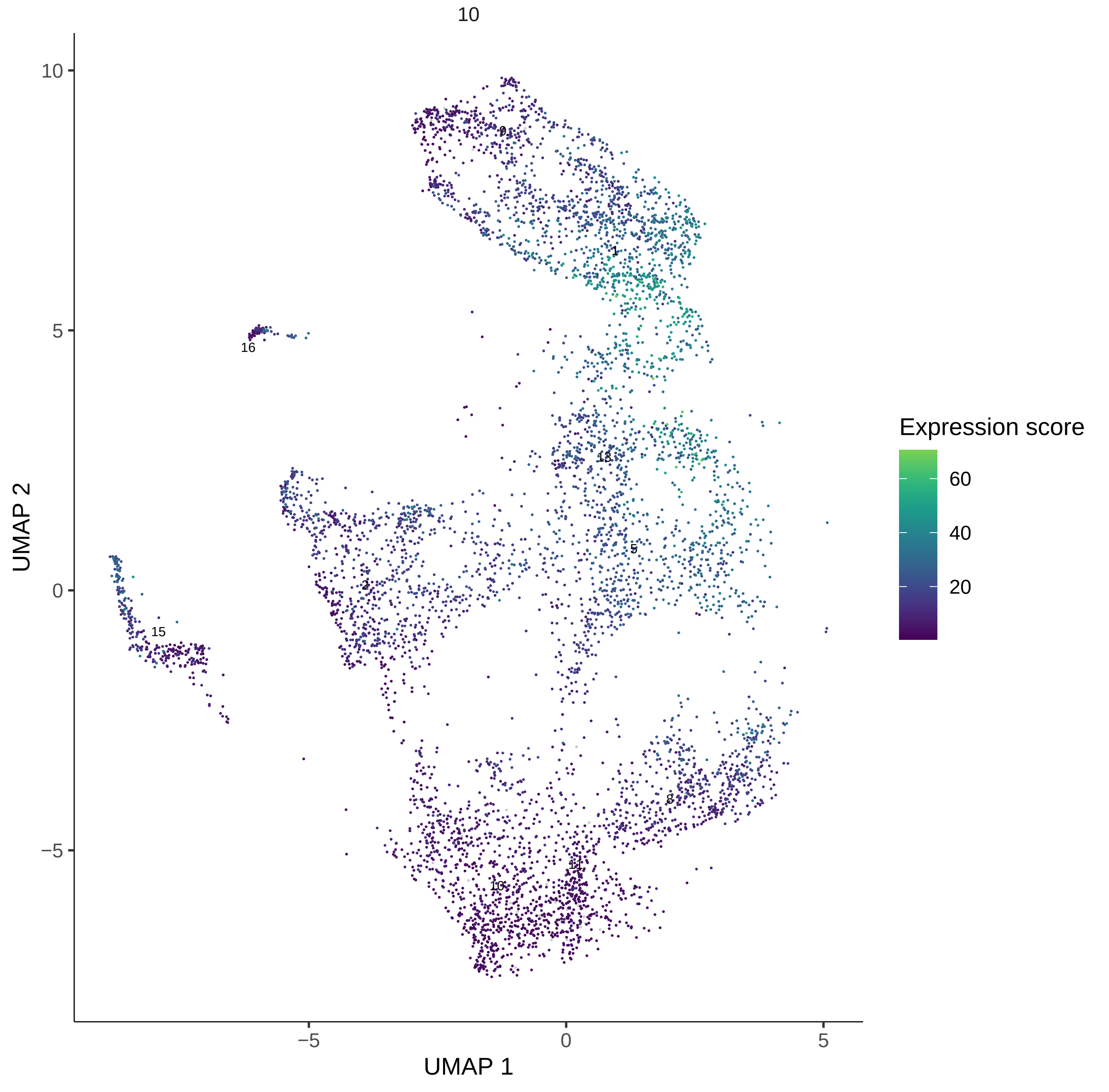
10
UMAP 1
UMAP 2
Expression score
60
40
20
10
5
0
−5
9
1
16
13
5
2
15
8
11
10

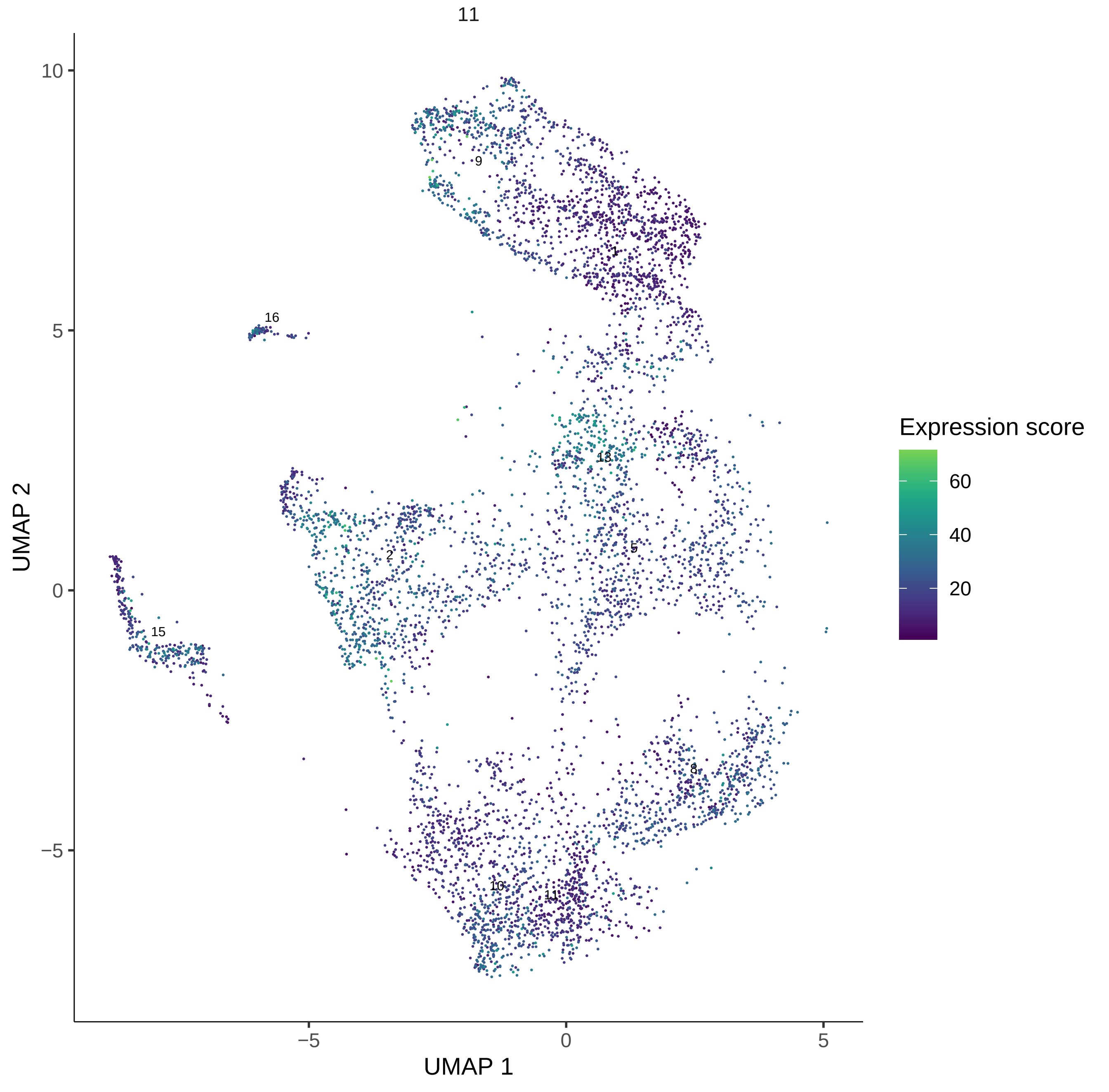

11
10
5
0
−5
−5
0
5
UMAP 1
UMAP 2
9
1
16
13
2
5
15
8
10
11
Expression score
60
40
20

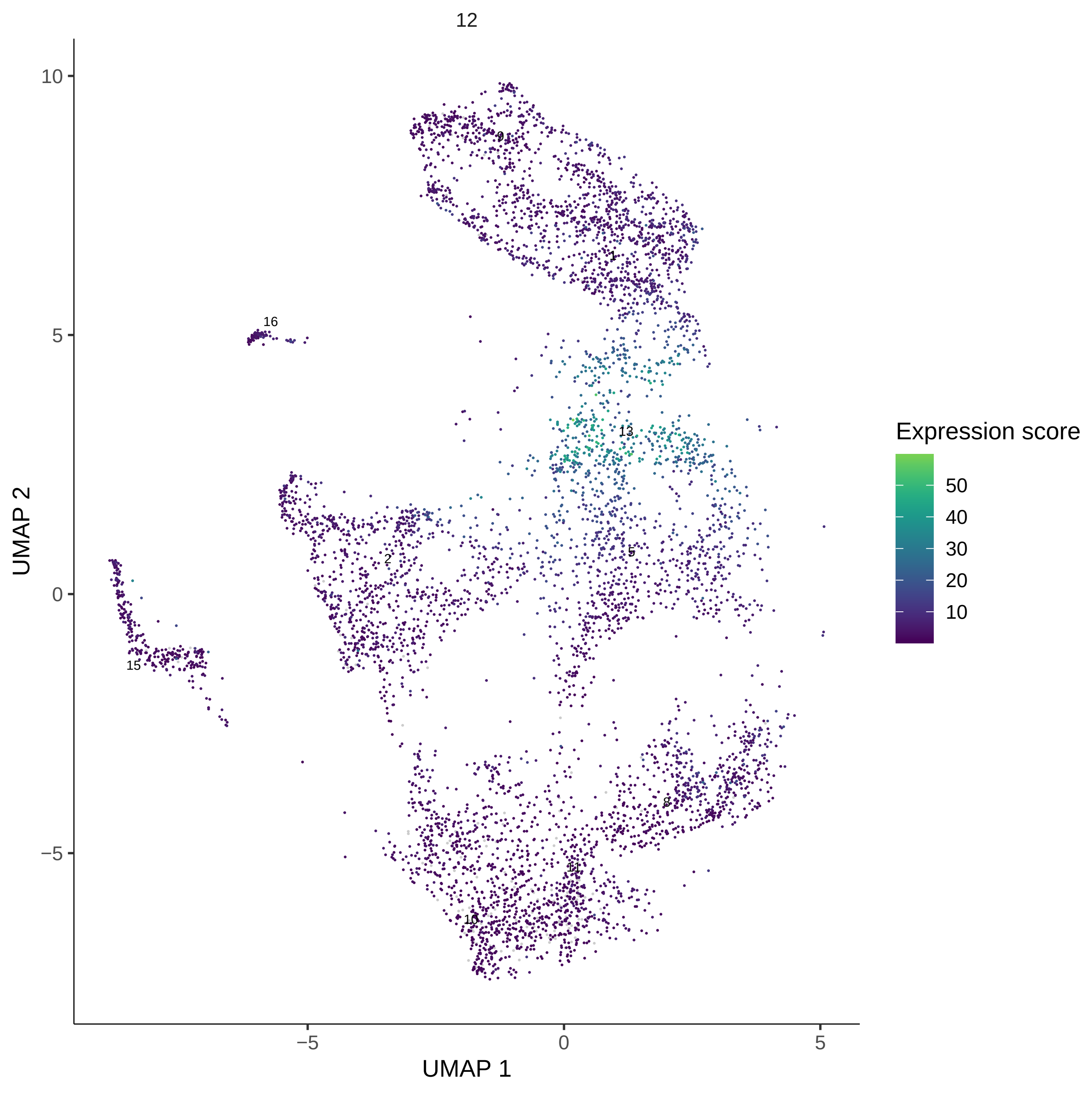
12
UMAP 2
UMAP 1
10
5
0
−5
−5
0
5
9
1
16
13
5
2
15
8
11
10
Expression score
50
40
30
20
10

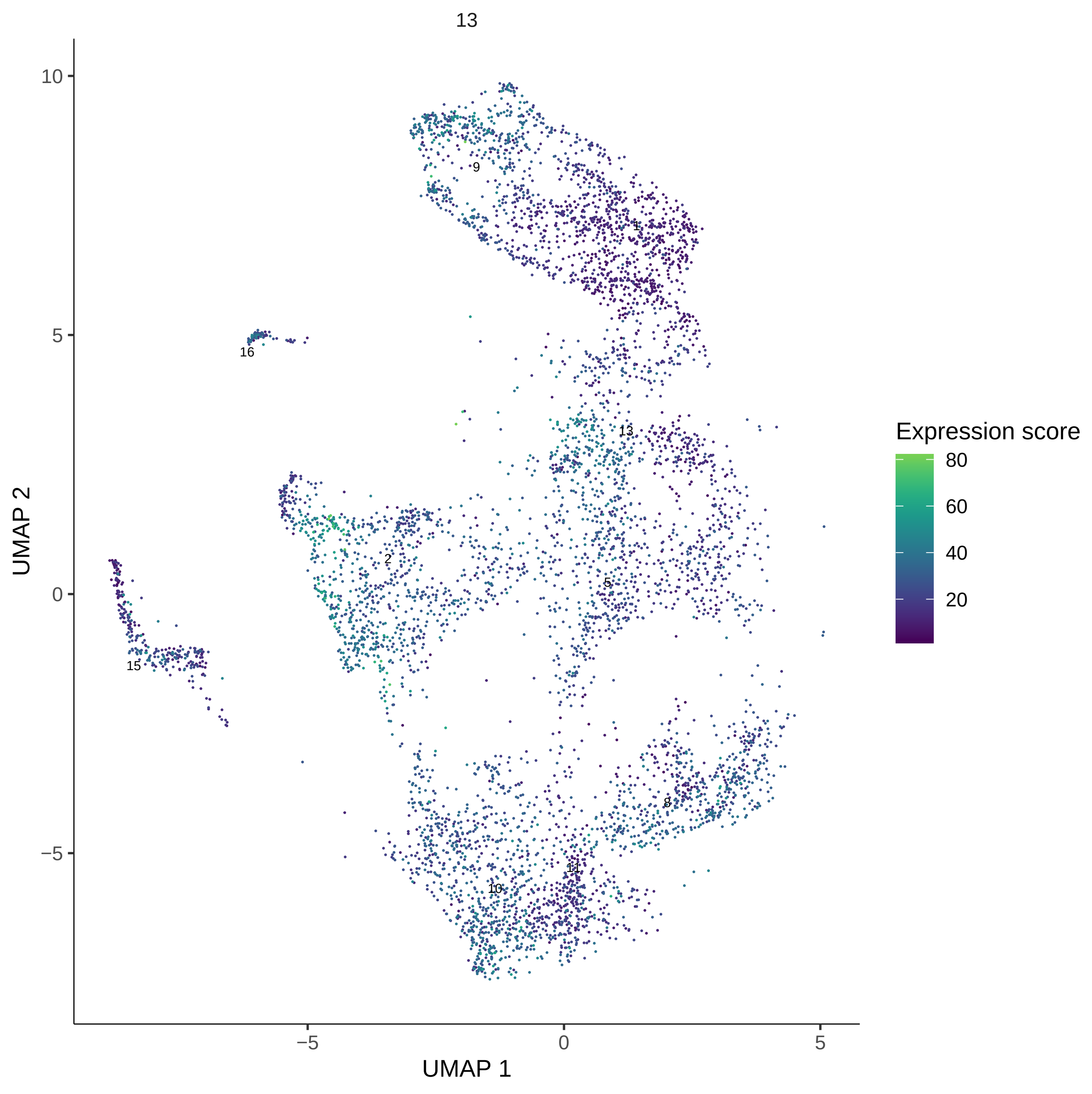
13
UMAP 2
UMAP 1
10
5
0
−5
−5
0
5
9
1
16
13
2
5
15
8
11
10
Expression score
80
60
40
20

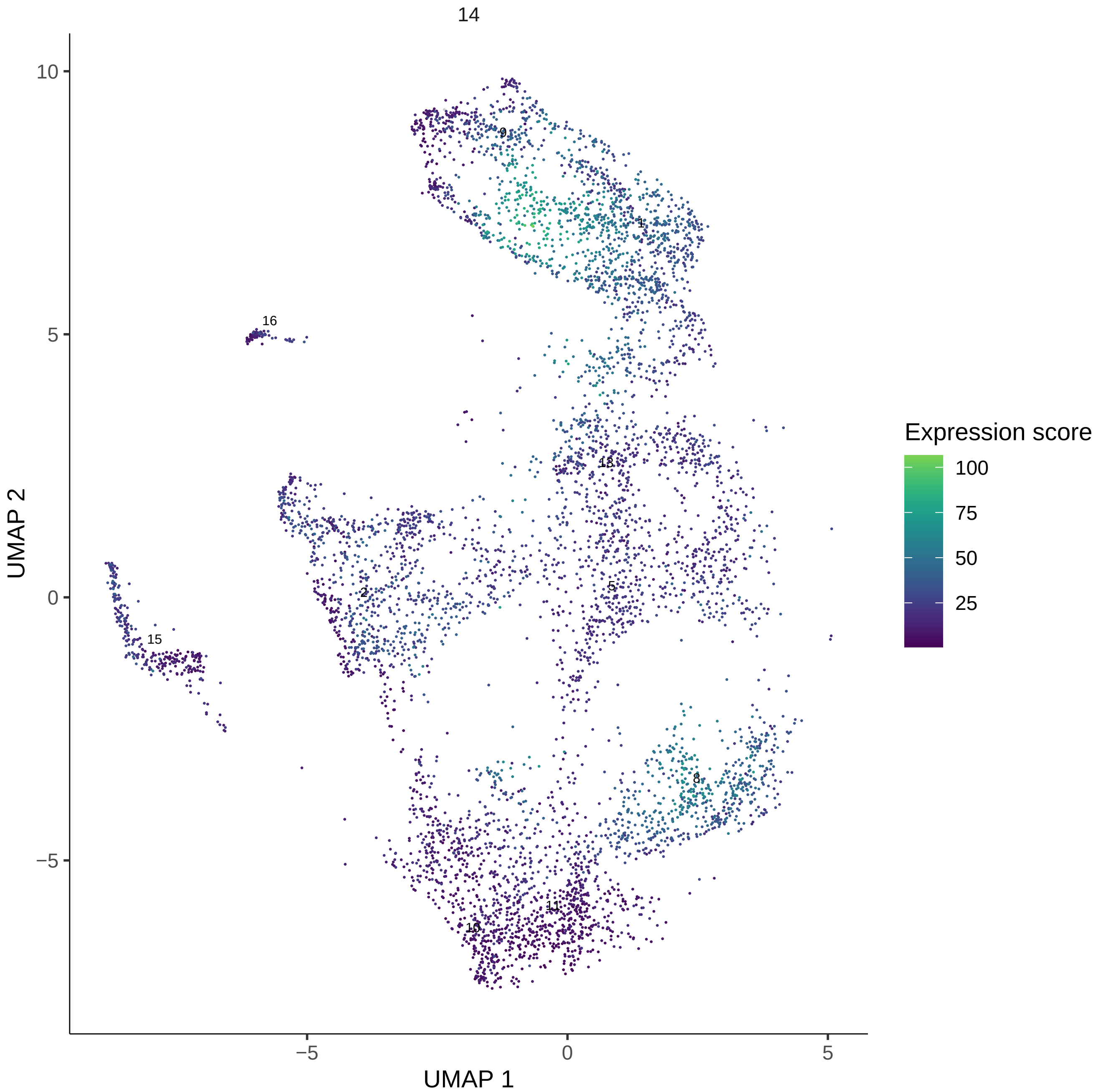
14
UMAP 1
UMAP 2
Expression score
100
75
50
25
10
5
0
−5
9
1
16
13
2
5
15
8
11
10

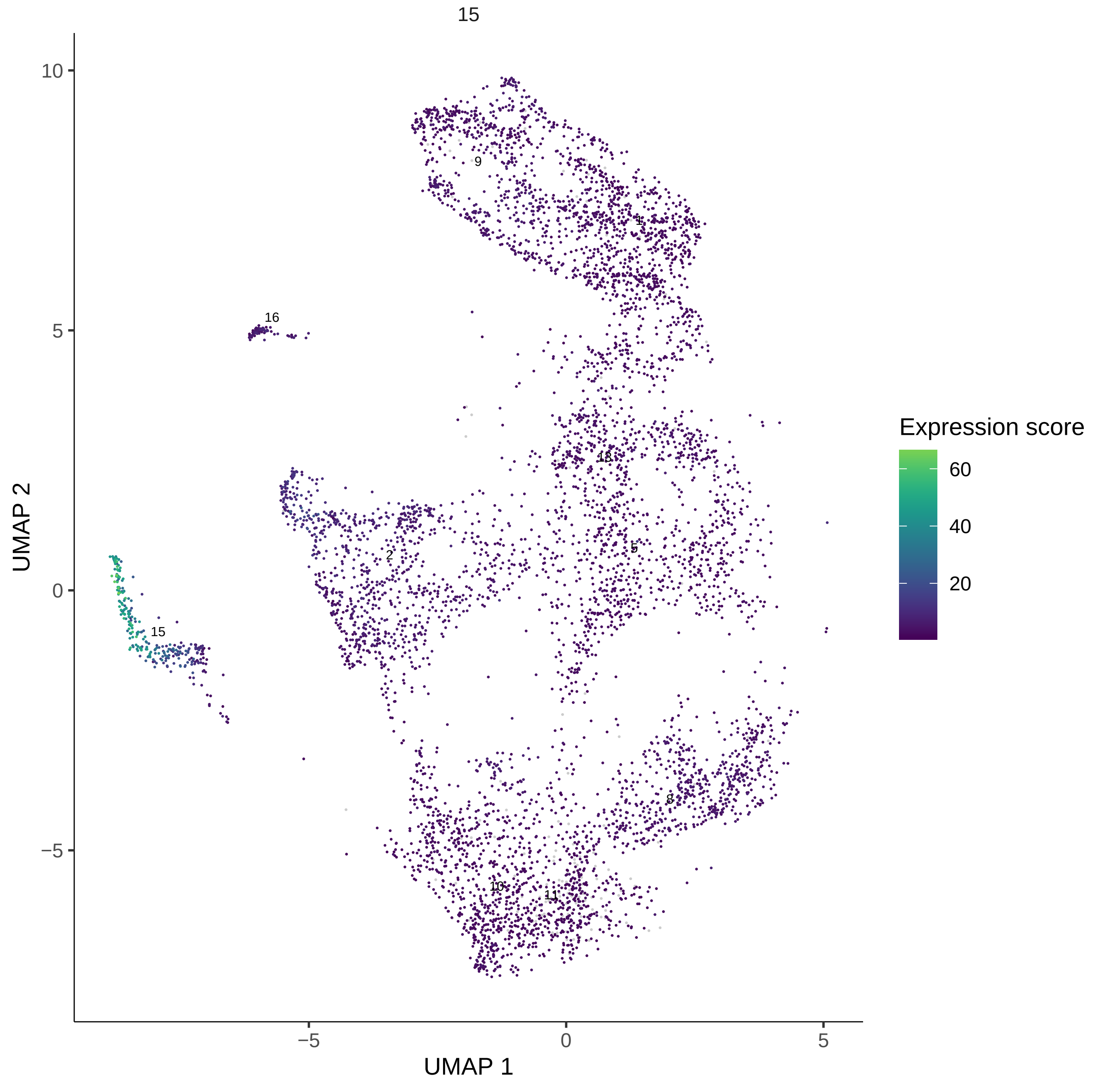

15
Expression score
60
40
20
UMAP 2
UMAP 1
10
5
0
−5
−5
0
5
9
1
16
13
2
5
15
8
10
11

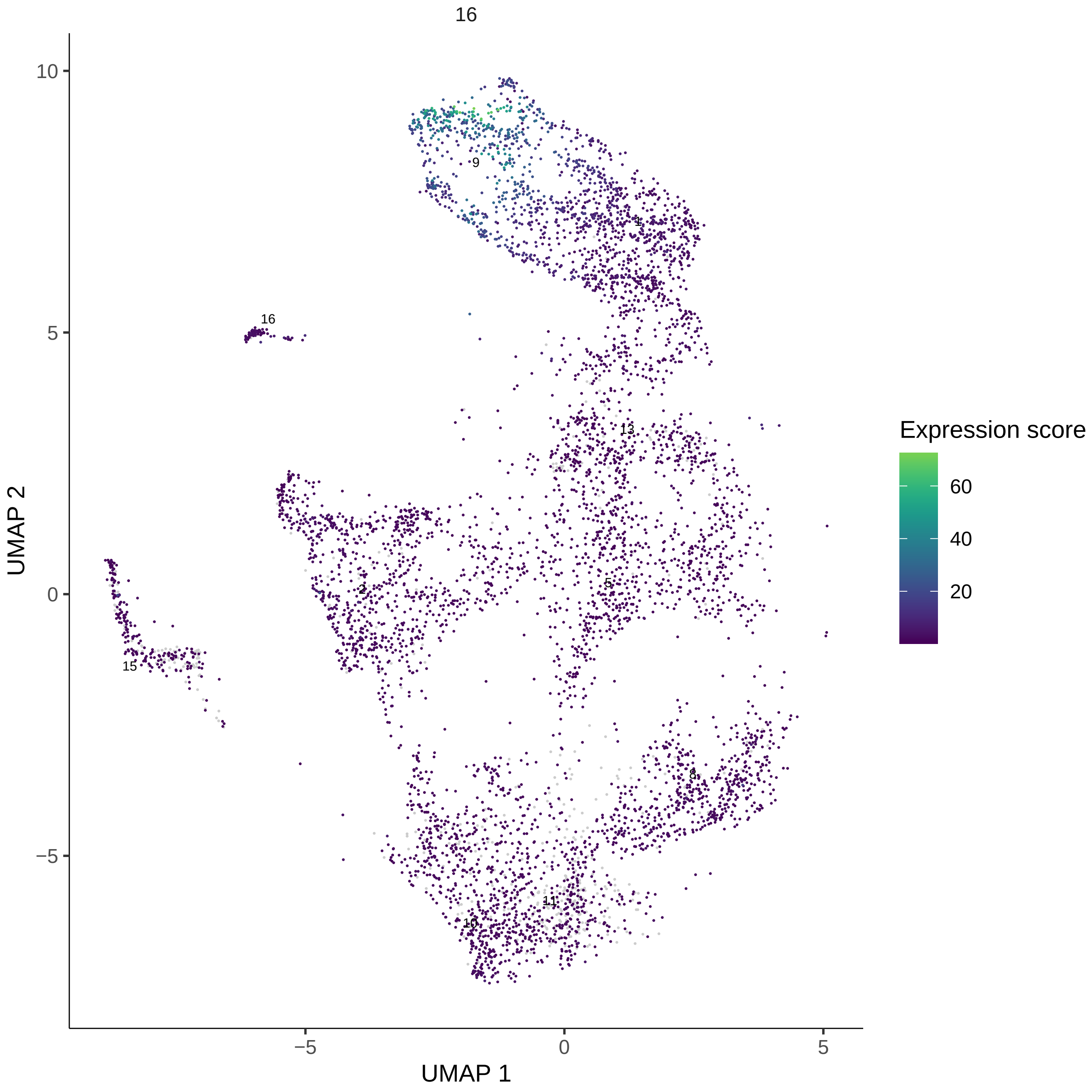

16
UMAP 2
UMAP 1
10
5
0
−5
−5
0
5
9
1
16
13
2
5
15
8
11
10
Expression score
60
40
20

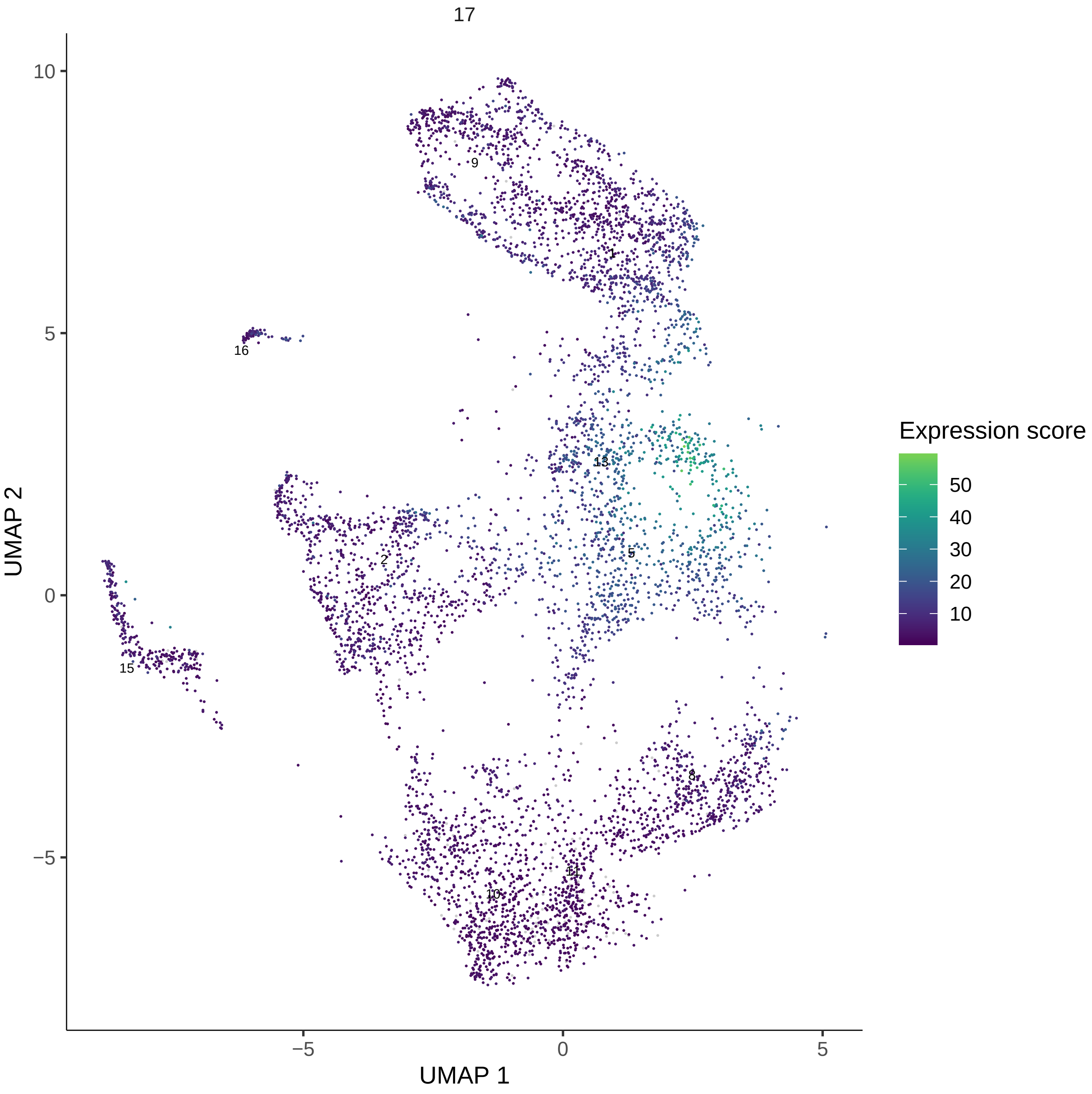
17
Expression score
50
40
30
20
10
UMAP 2
UMAP 1
10
5
0
−5
−5
0
5
9
1
16
13
2
5
15
8
11
10

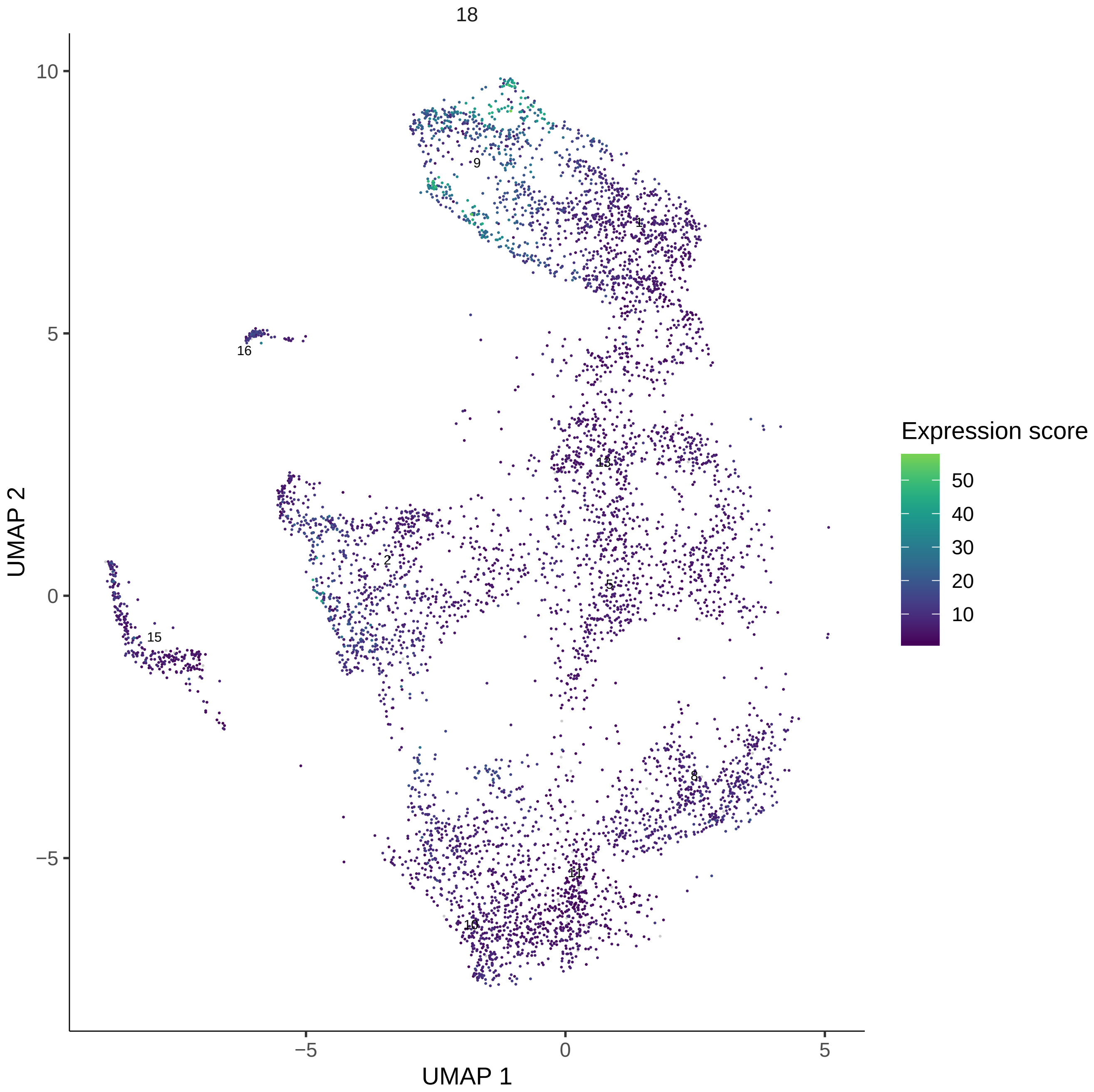
18
UMAP 2
UMAP 1
10
5
0
−5
−5
0
5
9
1
16
13
2
5
15
8
11
10
Expression score
50
40
30
20
10

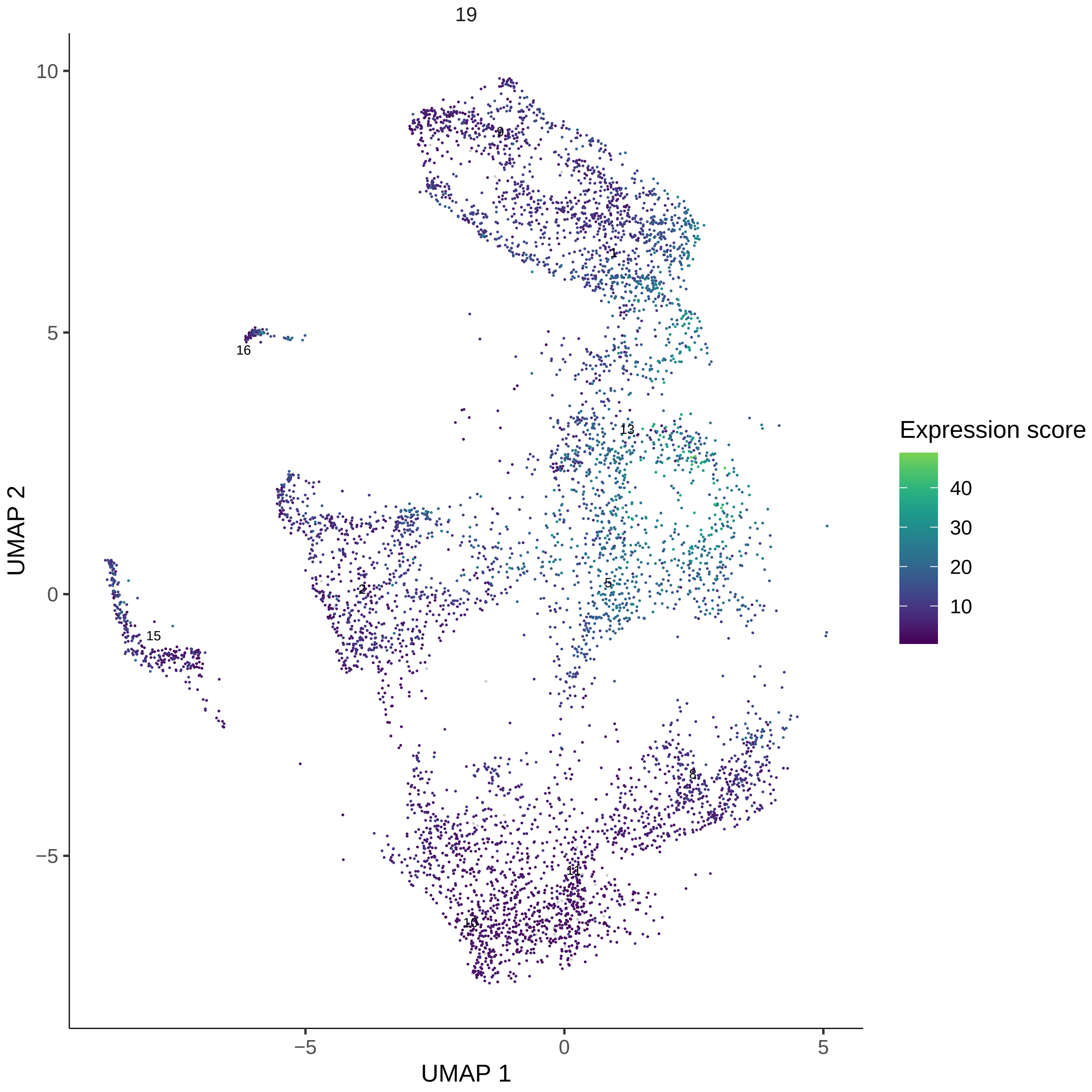
19
10
5
0
−5
−5
0
5
UMAP 1
UMAP 2
9
1
16
13
5
2
15
8
11
10
Expression score
40
30
20
10

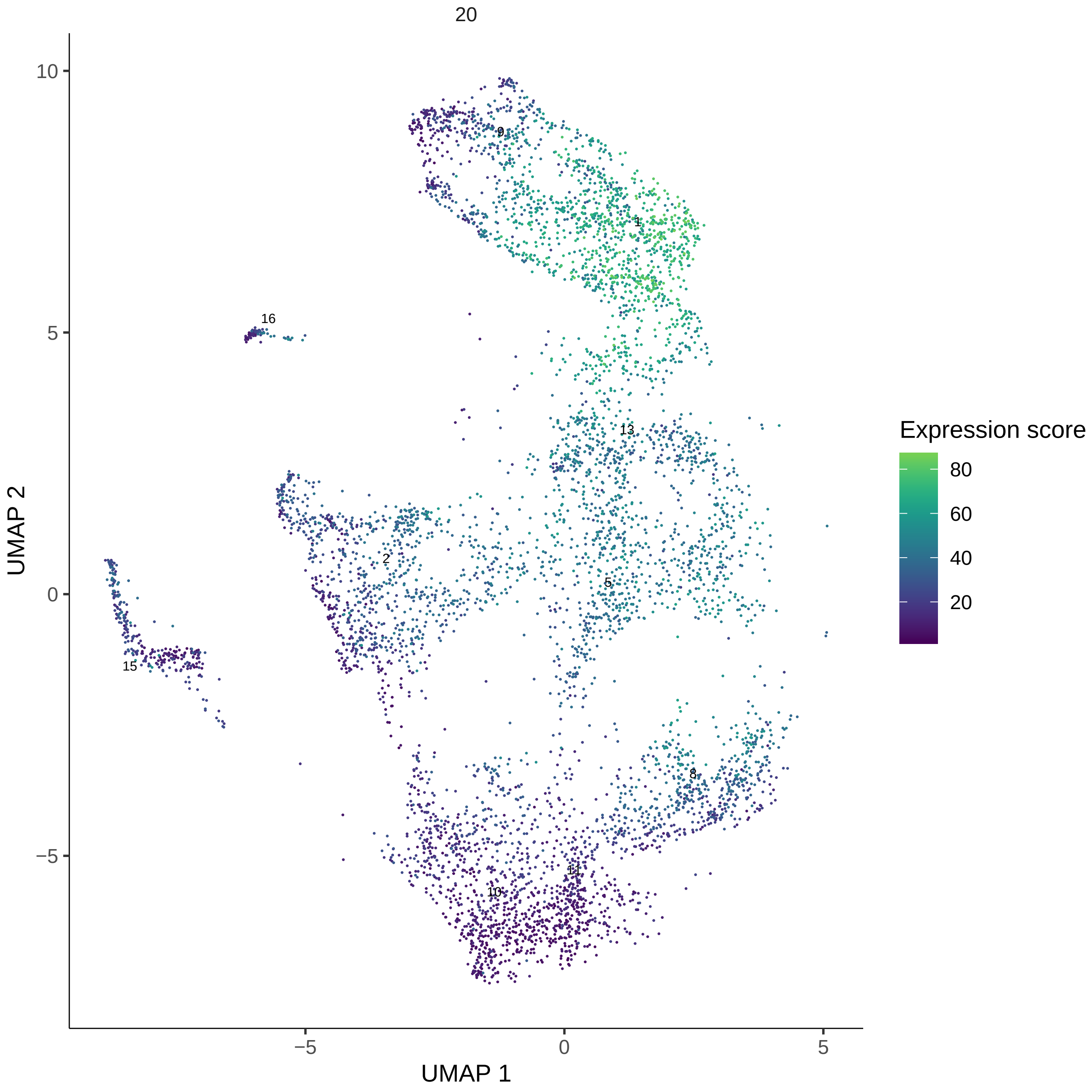
20
UMAP 1
UMAP 2
Expression score
80
60
40
20
10
5
0
−5
−5
0
5
9
1
16
13
2
5
15
8
11
10

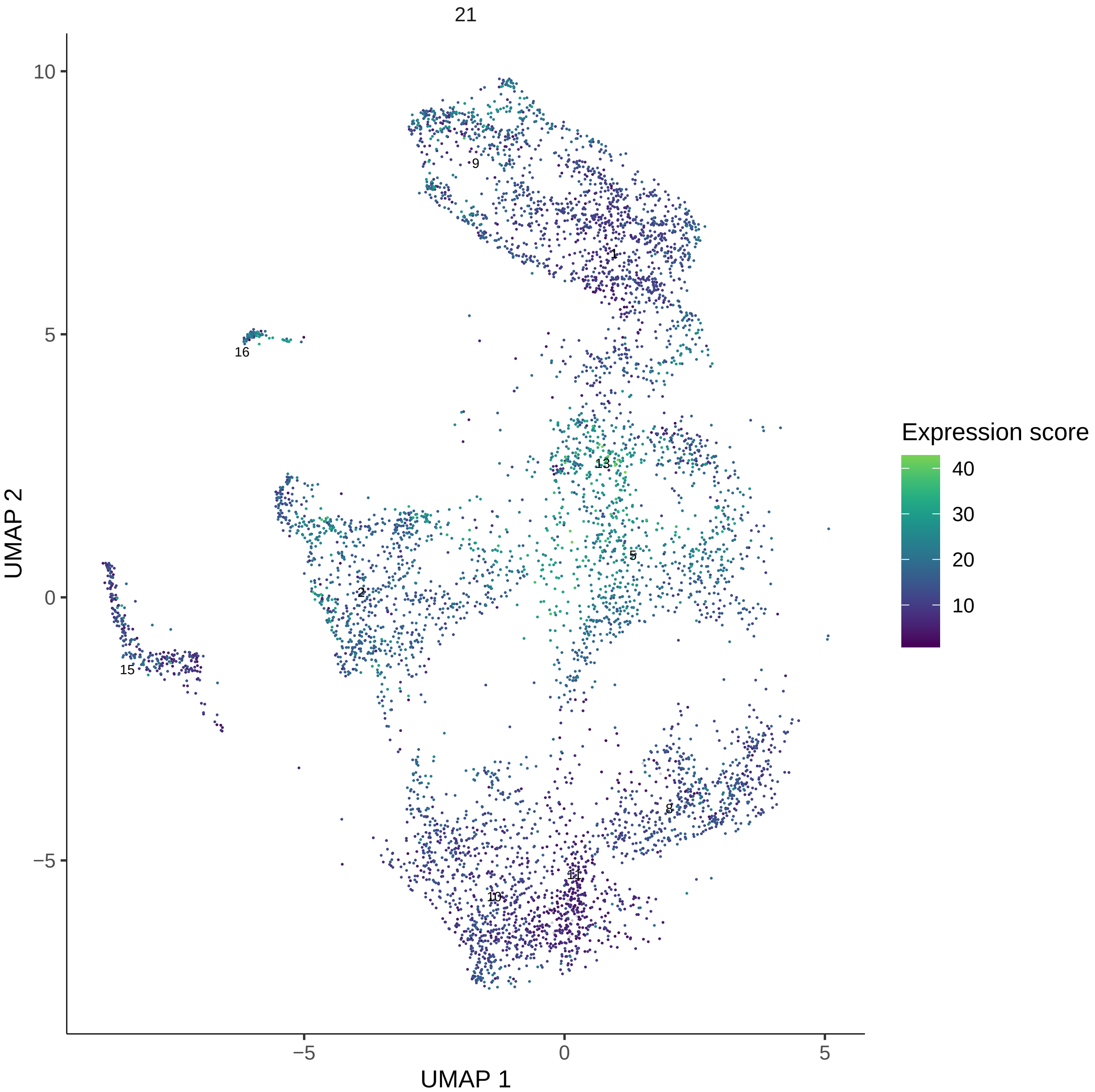
21
UMAP 1
UMAP 2
Expression score
40
30
20
10
9
1
16
13
5
2
15
8
11
10
−5
0
5
10

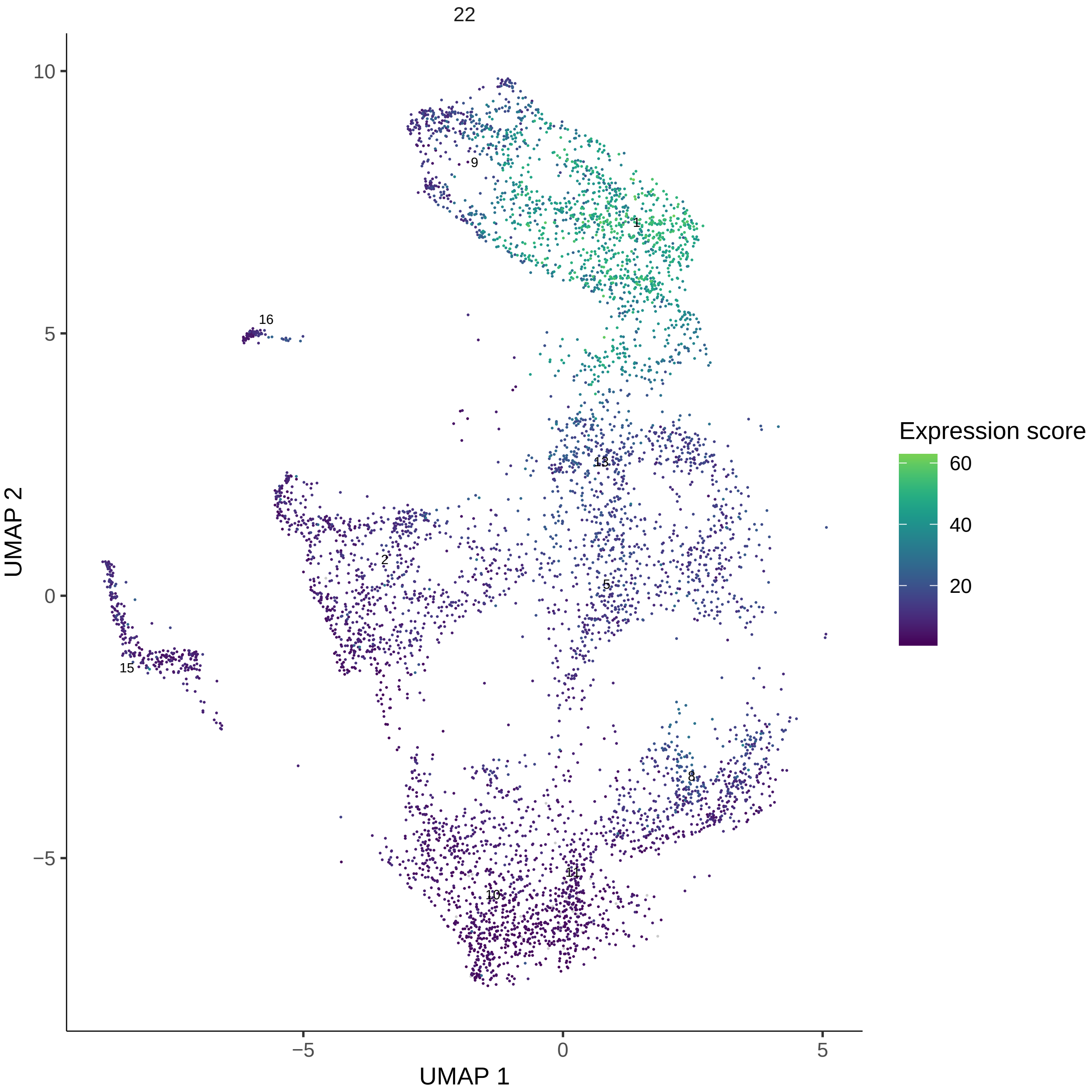

22
UMAP 1
UMAP 2
Expression score
60
40
20
10
5
0
−5
9
1
16
13
2
5
15
8
11
10

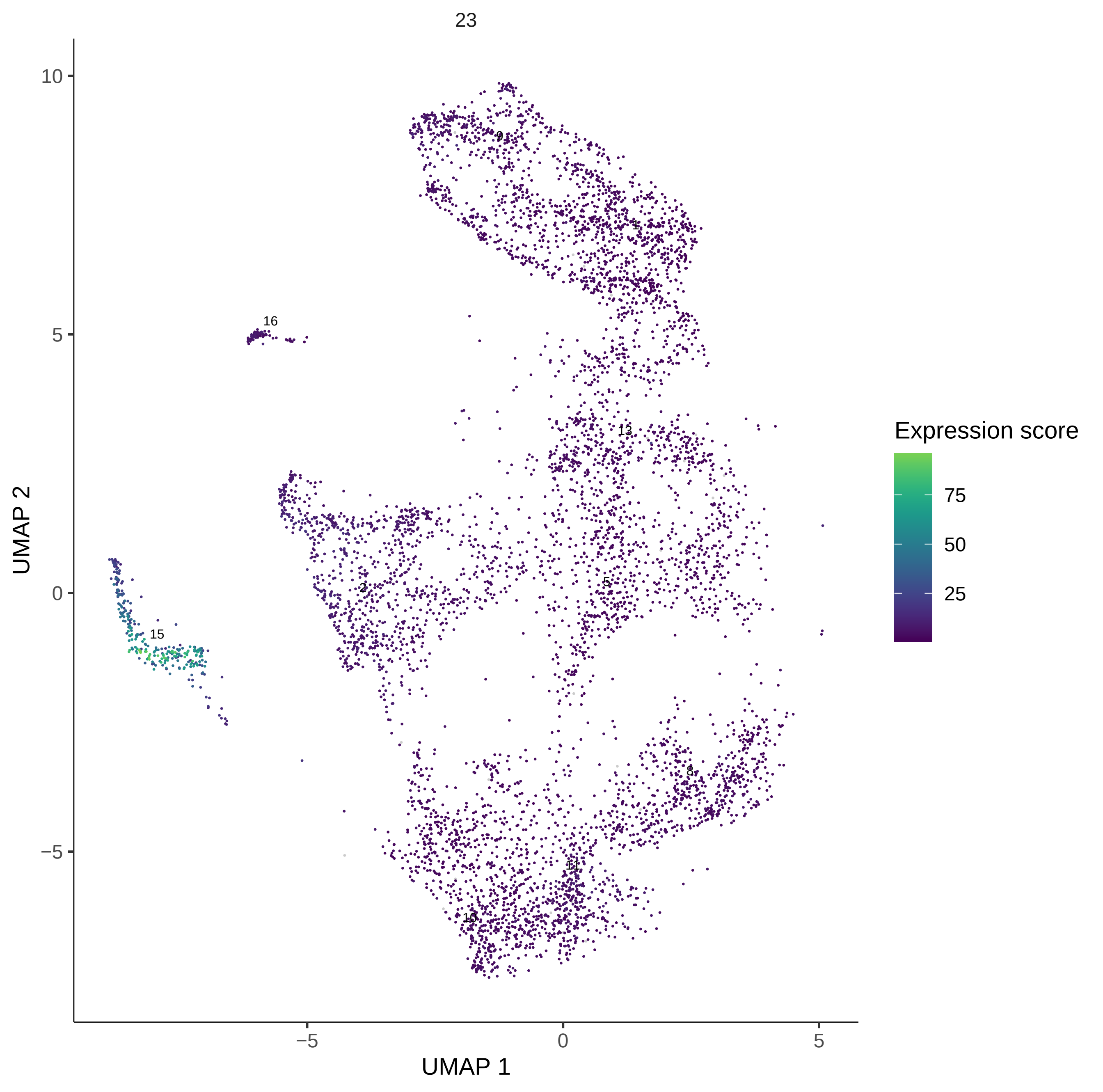

23
UMAP 1
UMAP 2
Expression score
75
50
25
10
5
0
−5
9
1
16
13
2
5
15
8
11
10

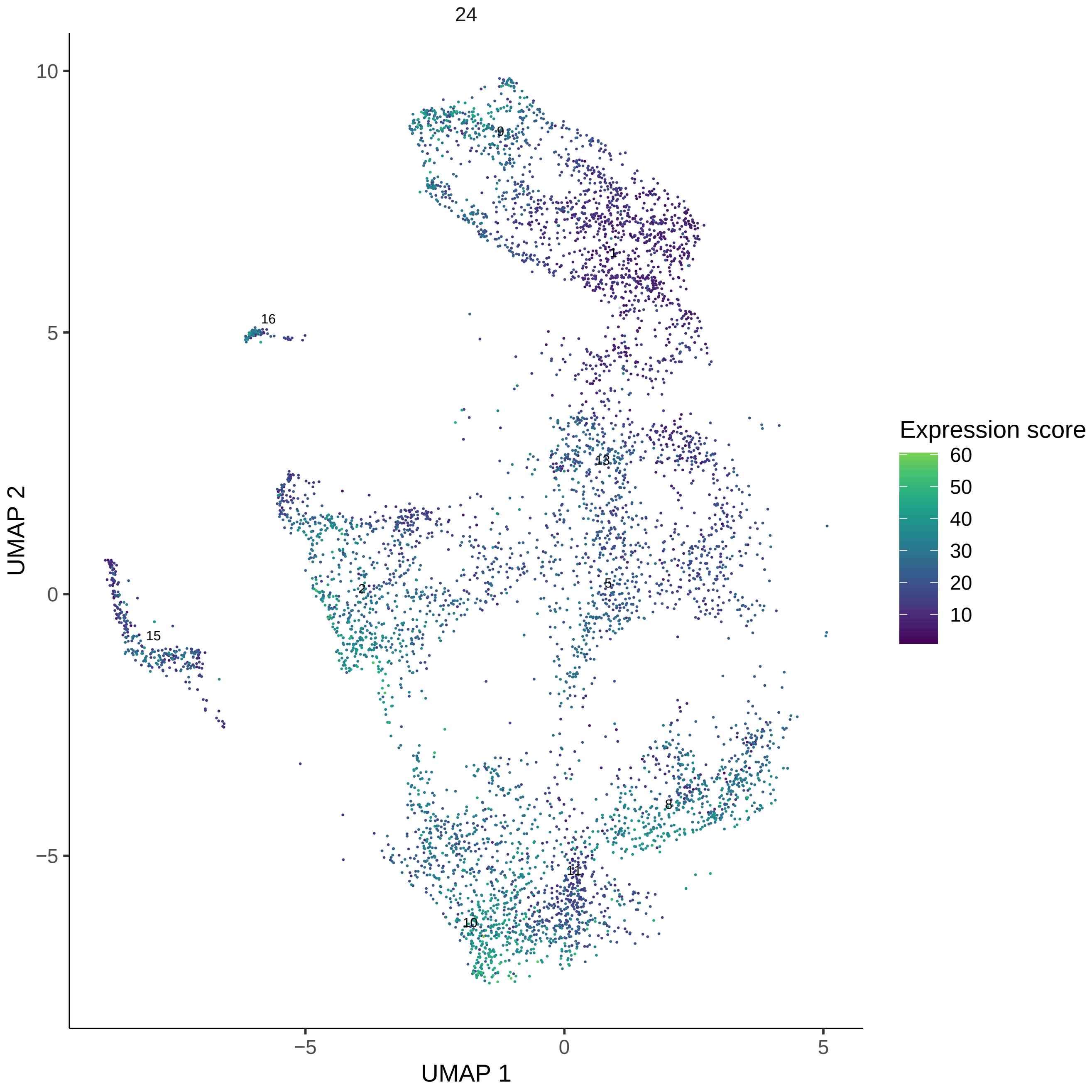

24
Expression score
60
50
40
30
20
10
UMAP 2
UMAP 1
10
5
0
−5
9
1
16
13
2
5
15
8
11
10

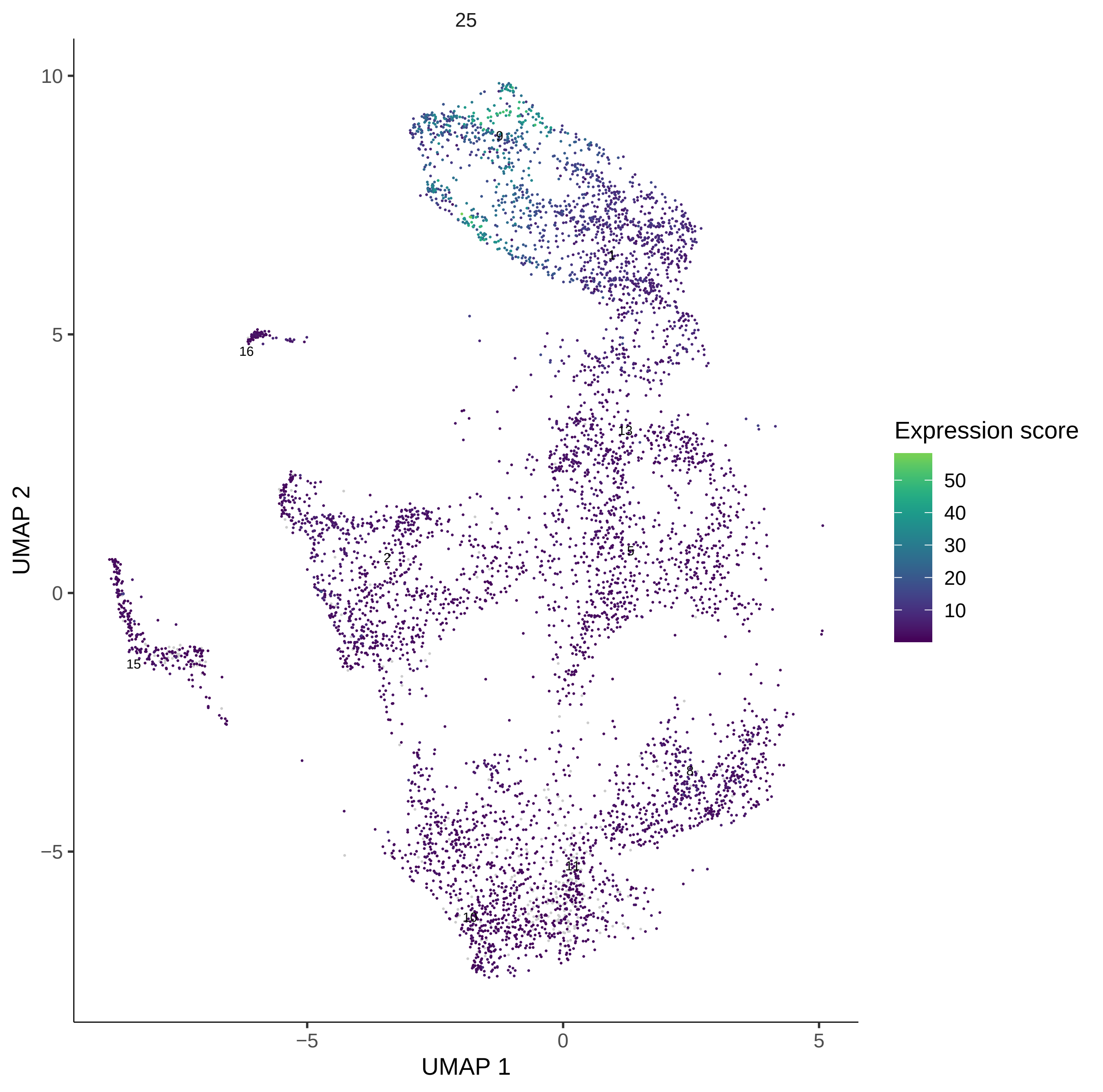

25
UMAP 2
UMAP 1
10
5
0
−5
−5
0
5
9
1
16
13
2
5
15
8
11
10
Expression score
50
40
30
20
10

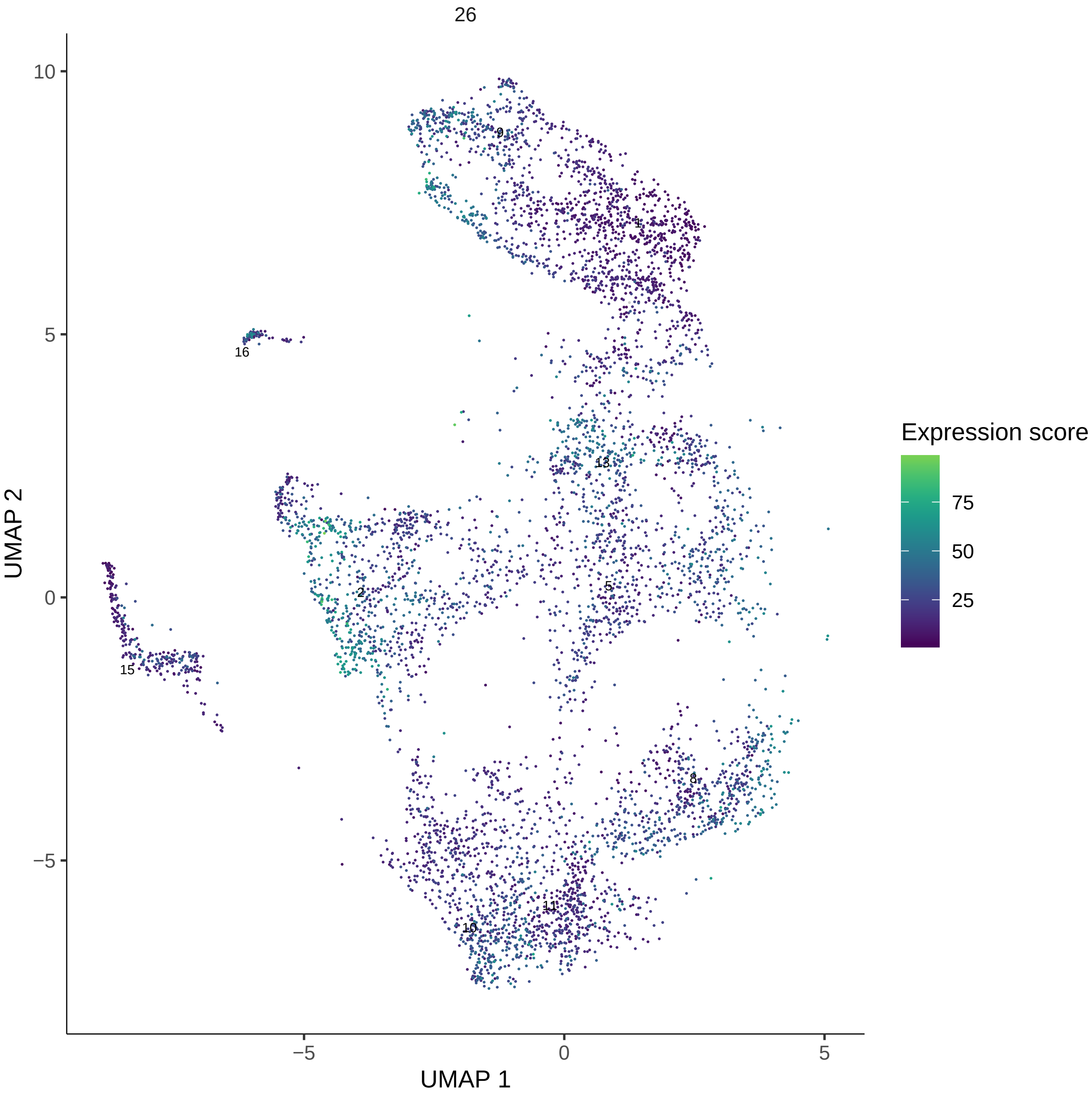
26
UMAP 1
UMAP 2
Expression score
75
50
25
10
5
0
−5
9
1
16
13
2
5
15
8
11
10

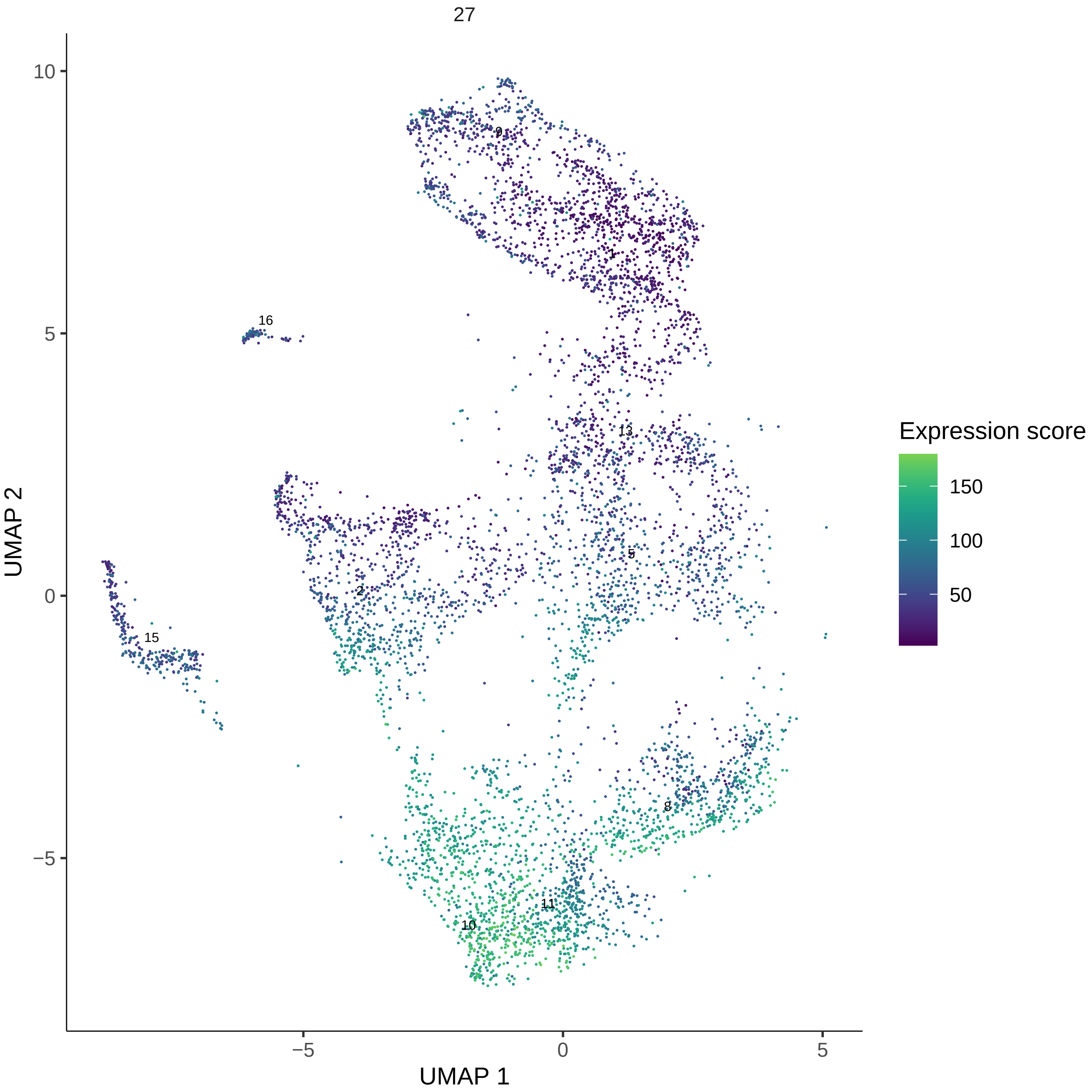
27
Expression score
150
100
50
UMAP 2
UMAP 1
10
5
0
−5
−5
0
5
9
1
16
13
5
2
15
8
11
10

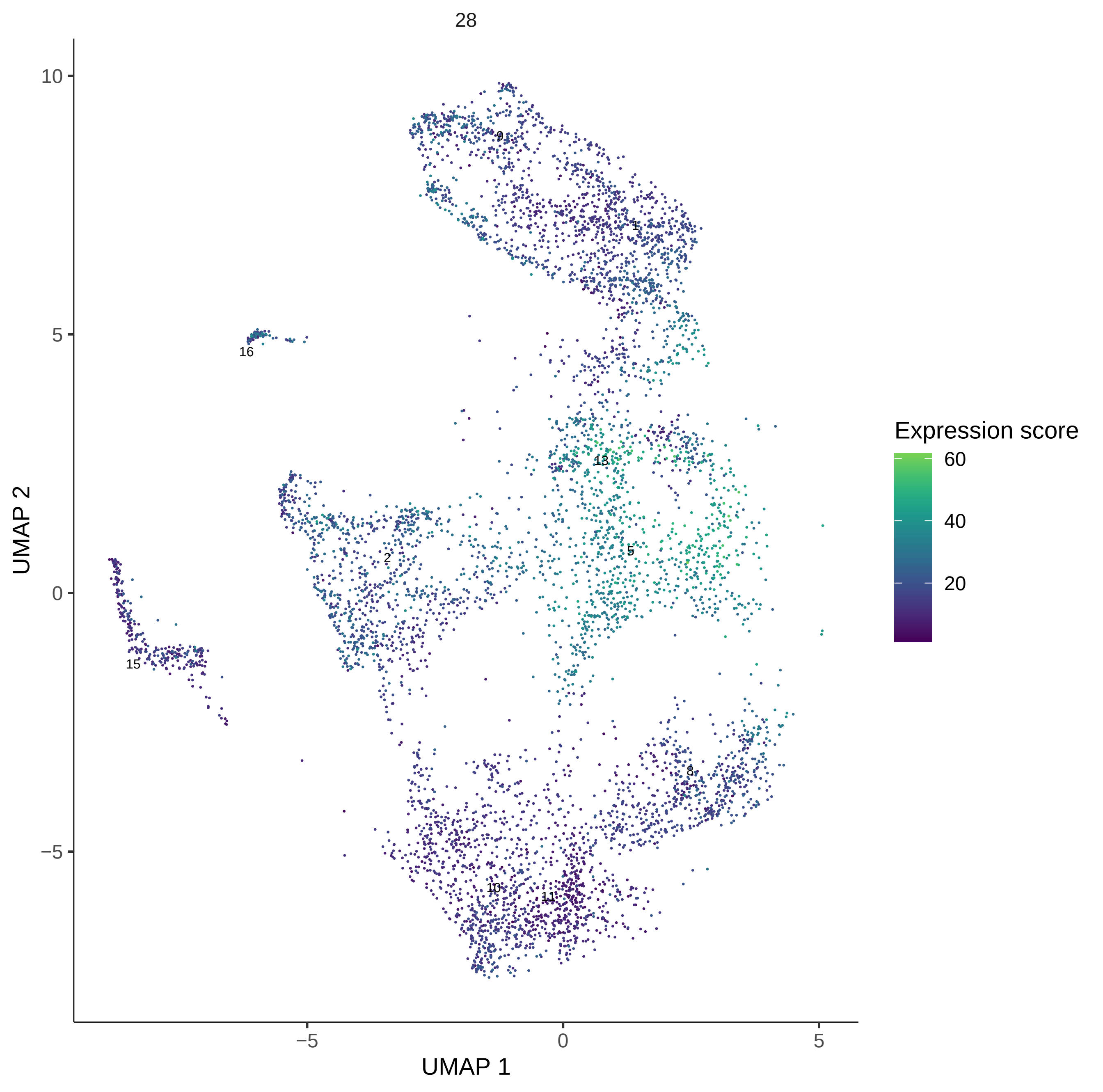

28
UMAP 2
UMAP 1
10
5
0
−5
−5
0
5
9
1
16
13
2
5
15
8
10
11
Expression score
60
40
20

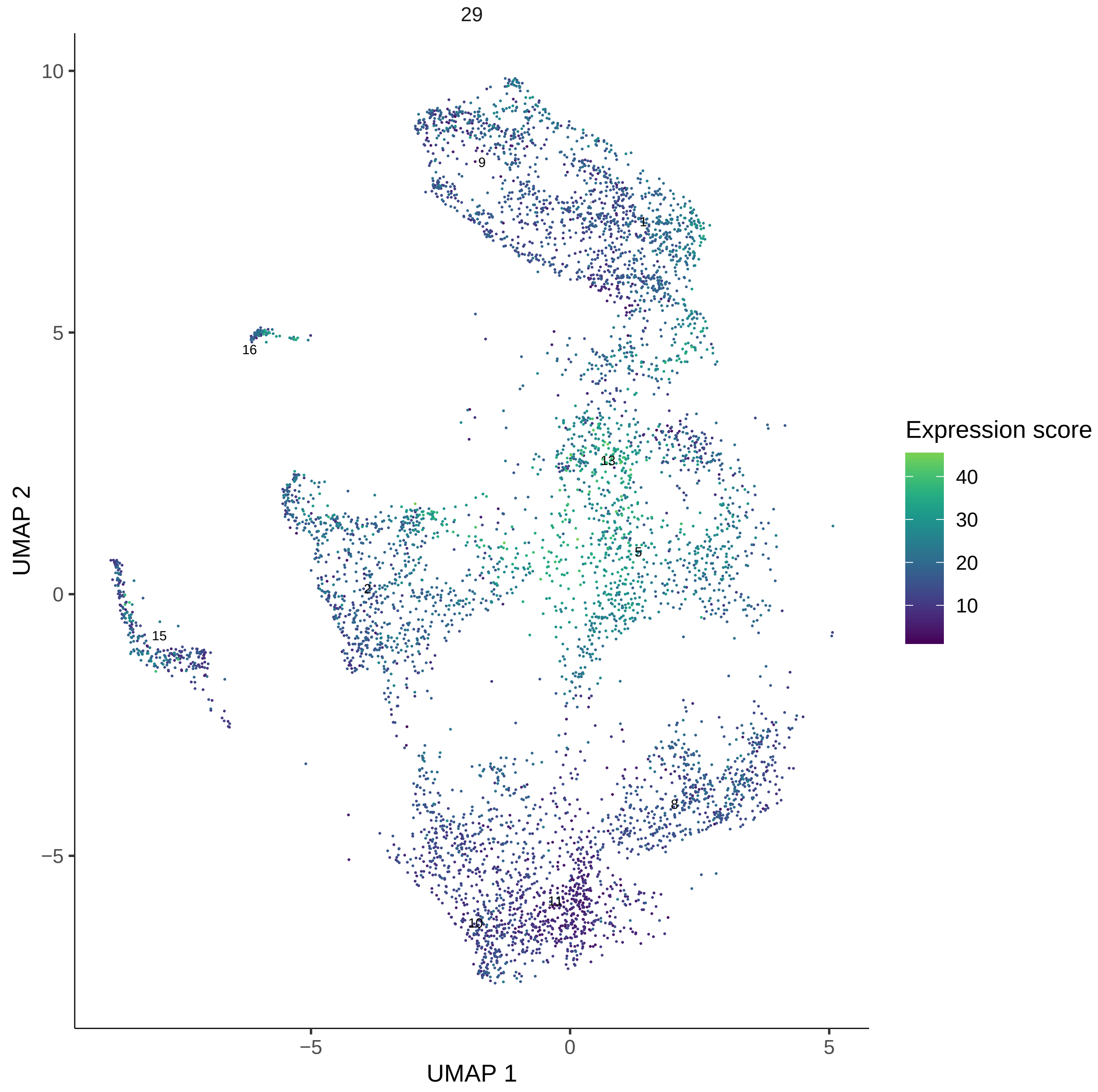

29
UMAP 2
UMAP 1
10
5
0
−5
−5
0
5
9
1
16
13
5
2
15
8
11
10
Expression score
40
30
20
10

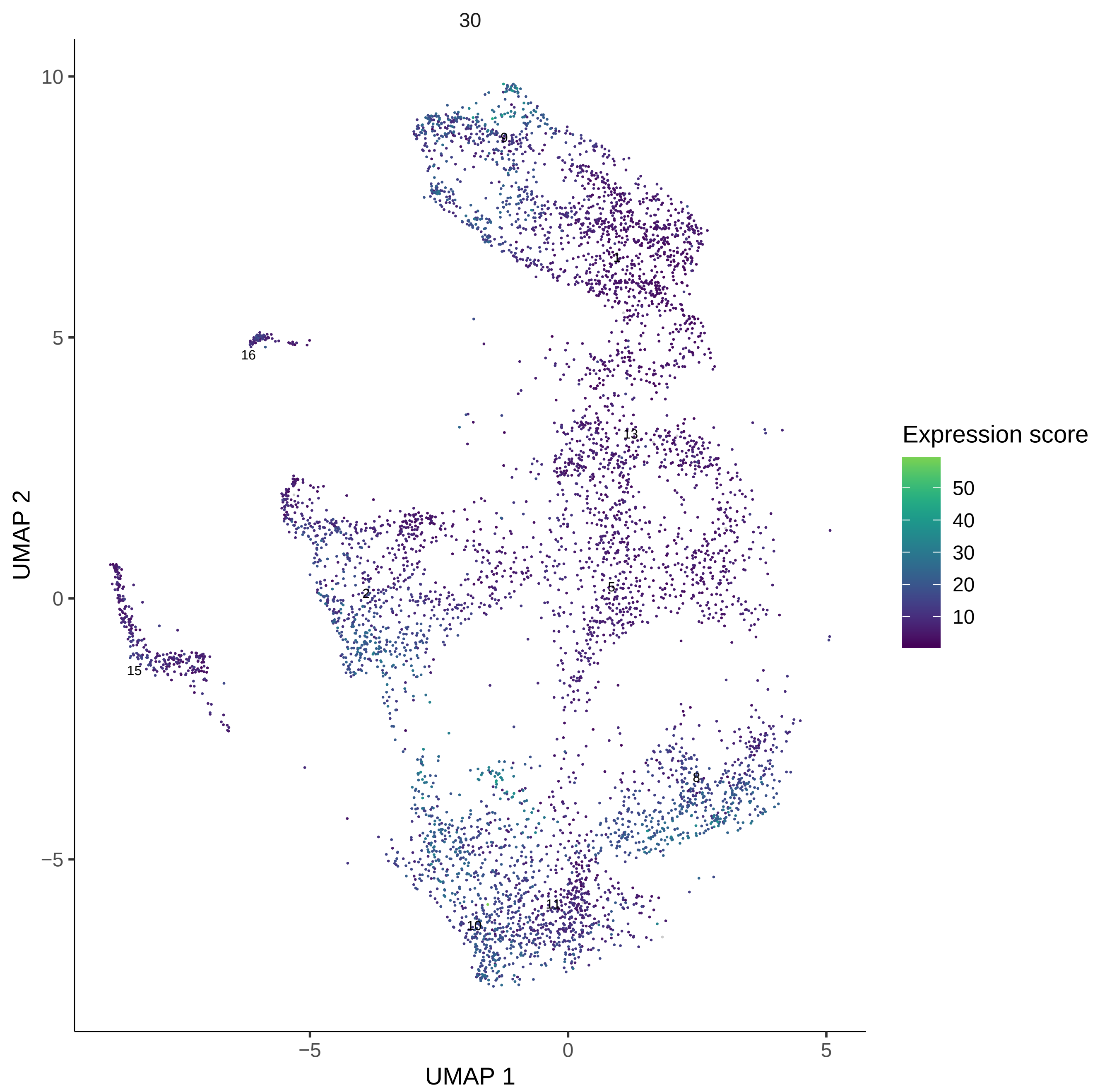
30
UMAP 2
UMAP 1
10
5
0
−5
−5
0
5
9
1
16
13
2
5
15
8
11
10
Expression score
50
40
30
20
10

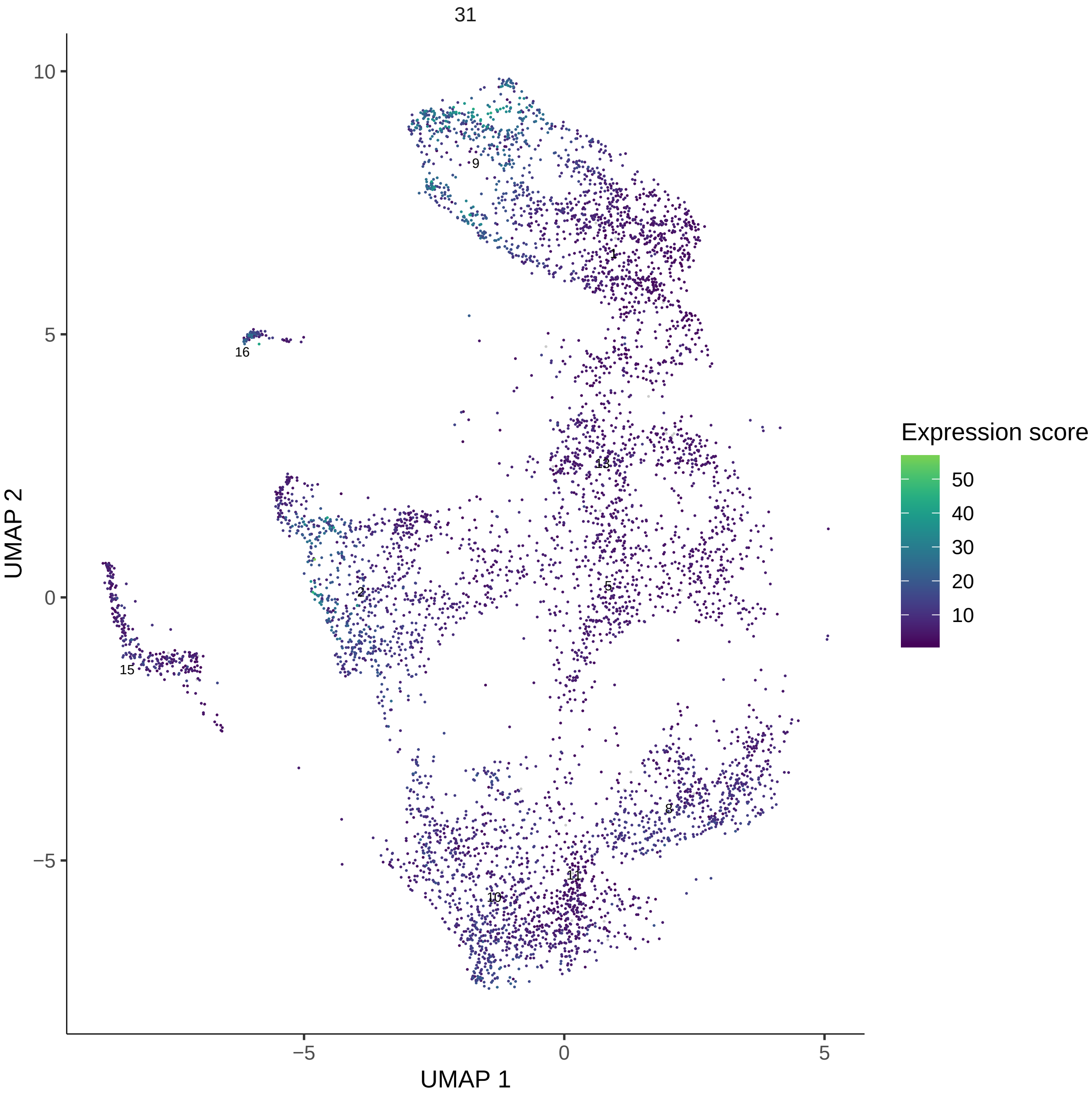

31
10
5
0
−5
−5
0
5
UMAP 1
UMAP 2
9
1
16
13
2
5
15
8
11
10
Expression score
50
40
30
20
10

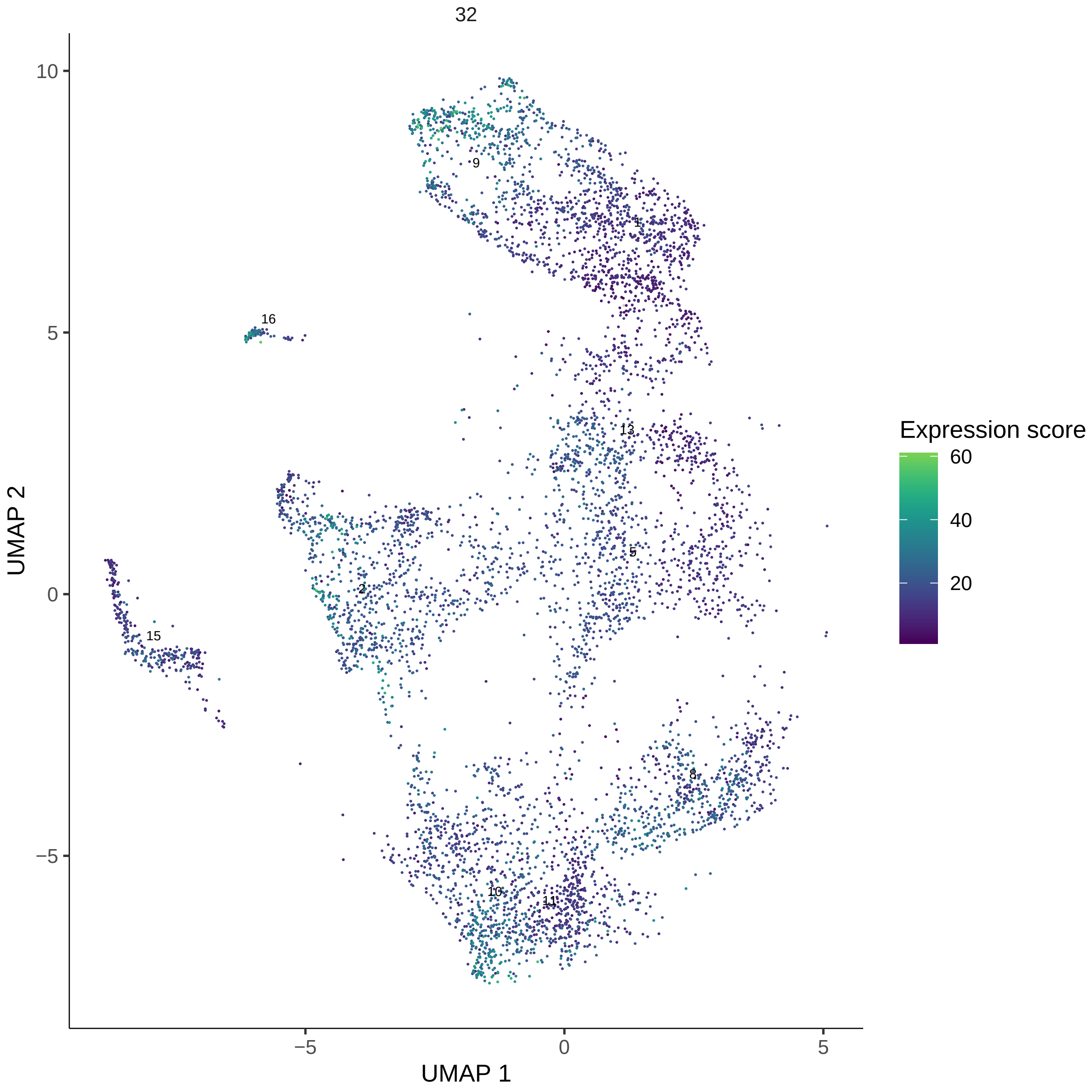
32
UMAP 1
UMAP 2
Expression score
60
40
20
9
1
16
13
5
2
15
8
10
11
−5
0
5
10

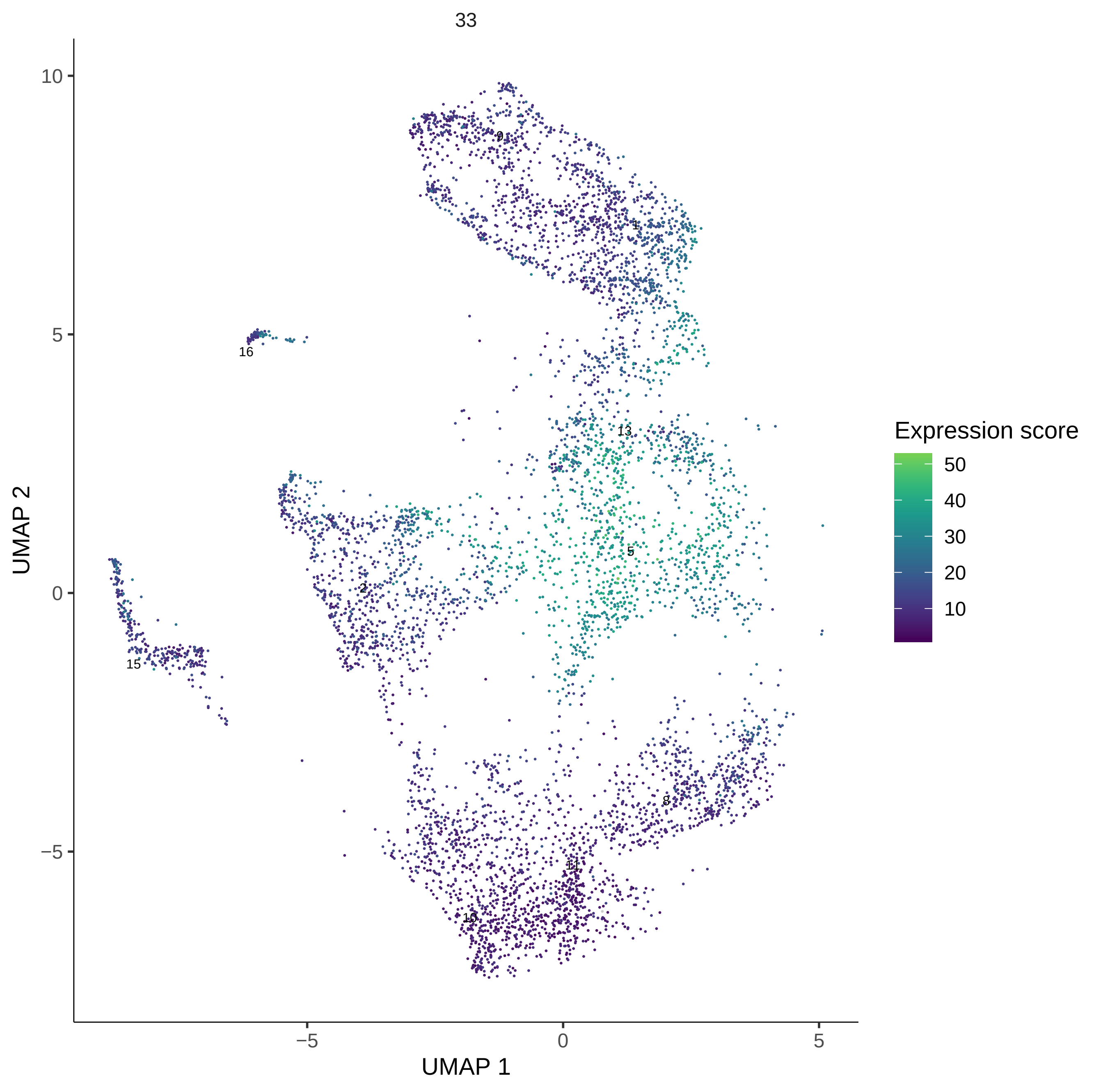
33
UMAP 1
UMAP 2
Expression score
50
40
30
20
10
9
1
16
13
5
2
15
8
11
10
−5
0
5

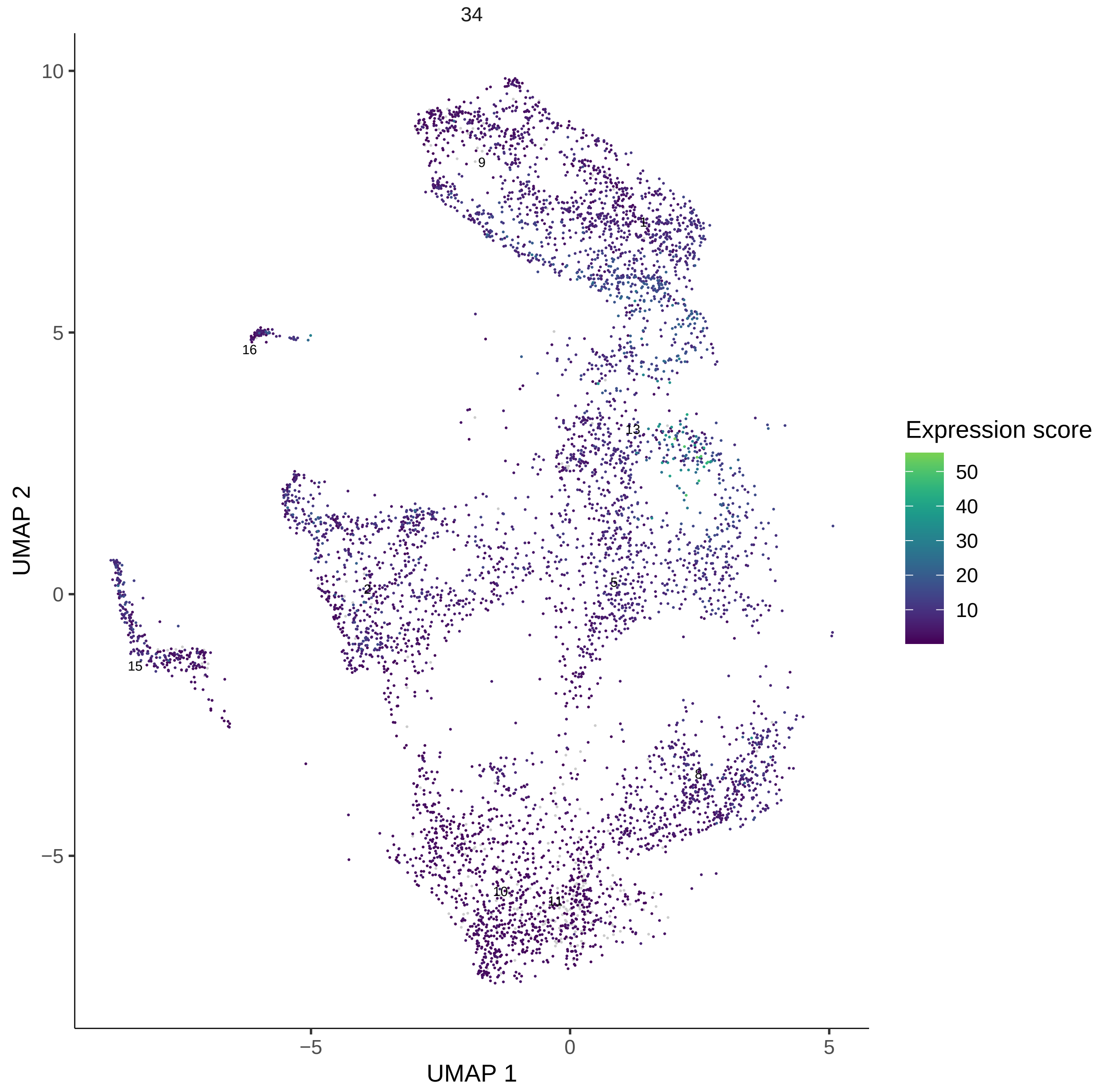
34
UMAP 1
UMAP 2
10
5
0
−5
5
0
−5
9
1
16
13
2
5
15
8
10
11
Expression score
50
40
30
20
10

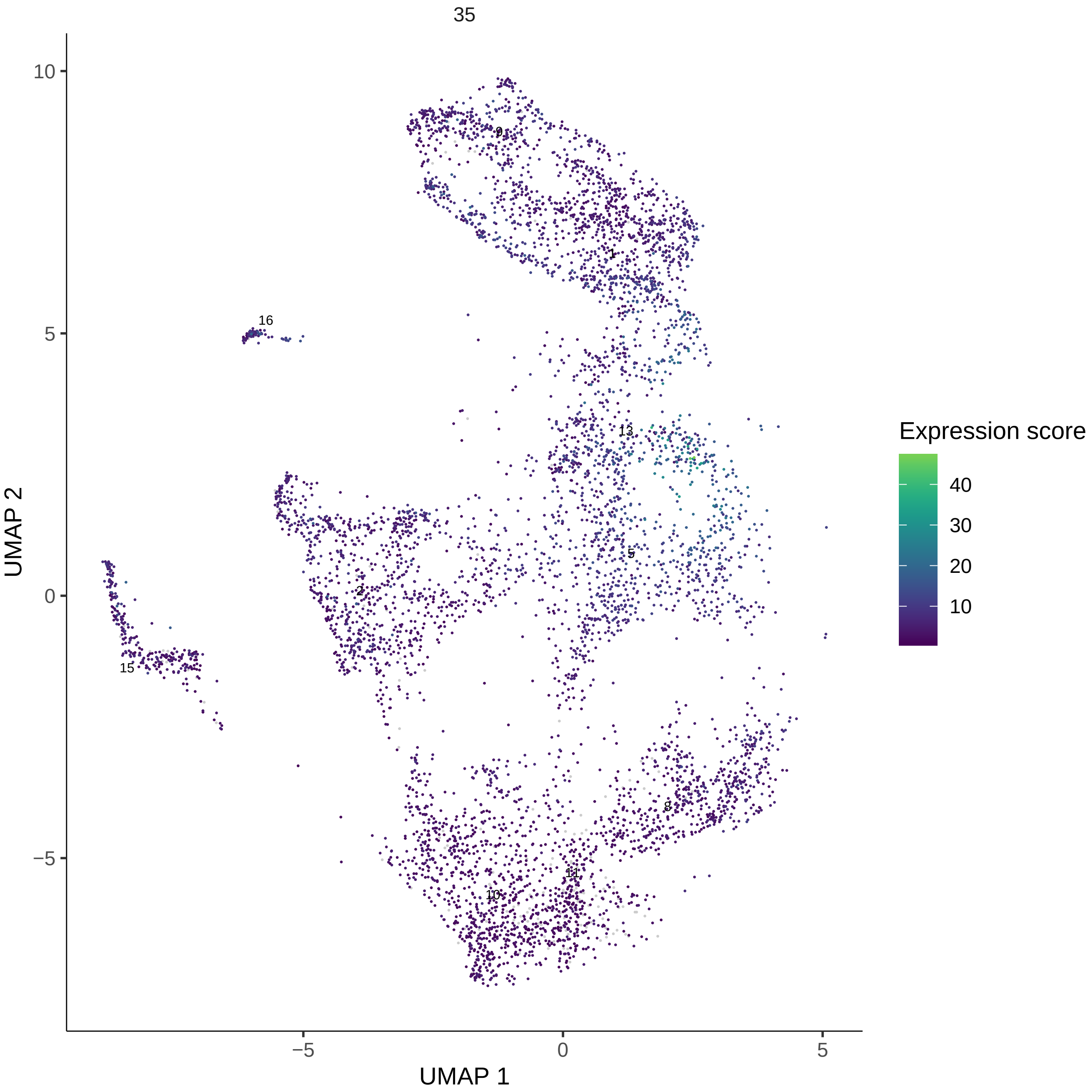
35
Expression score
40
30
20
10
UMAP 1
UMAP 2
10
5
0
−5
−5
0
5
9
1
16
13
5
2
15
8
11
10

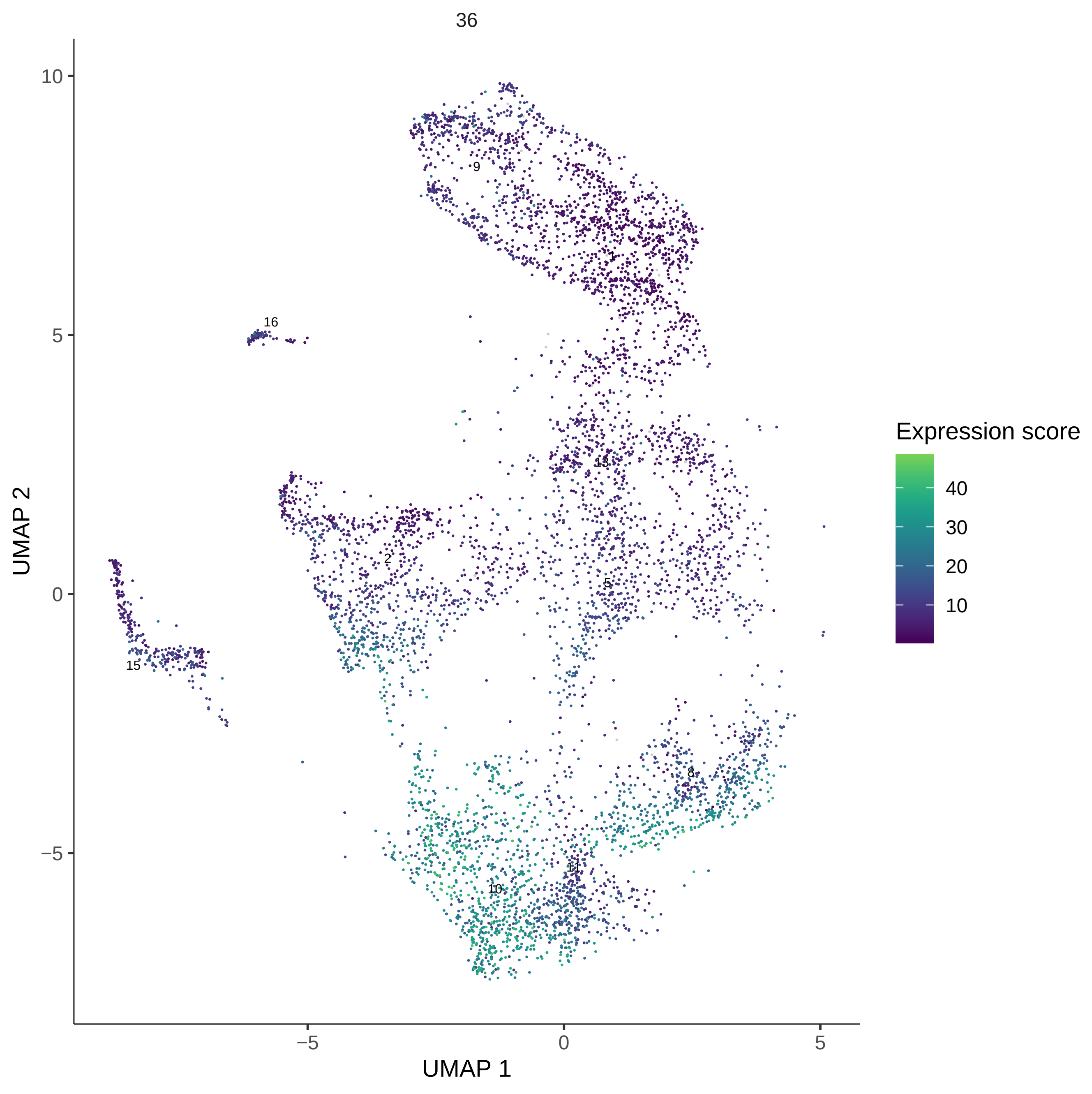
36
UMAP 1
UMAP 2
Expression score
40
30
20
10
10
5
0
−5
−5
0
5
9
1
16
13
2
5
15
8
11
10

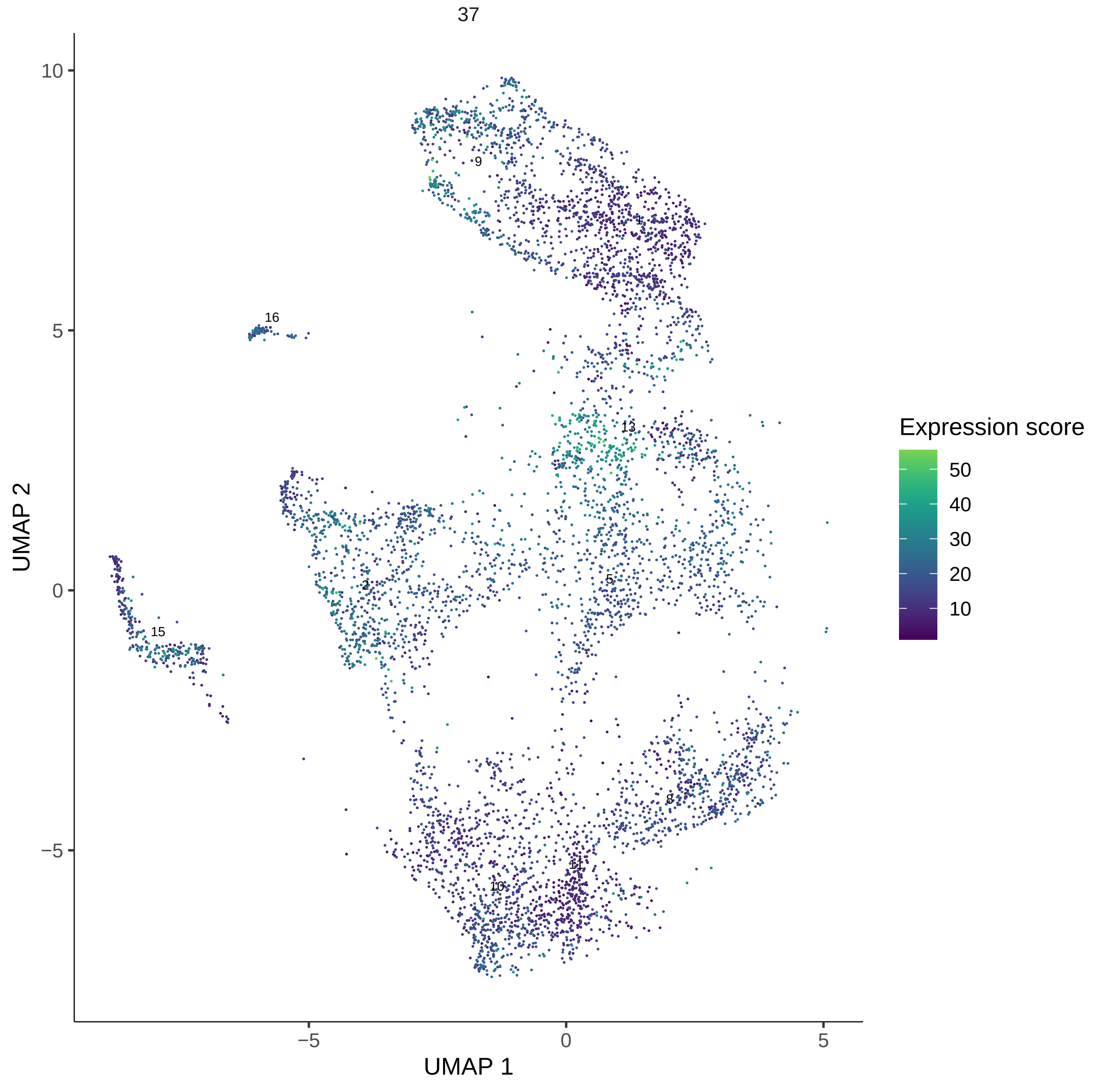

37
UMAP 2
UMAP 1
10
5
0
−5
−5
0
5
9
1
16
13
2
5
15
8
11
10
Expression score
50
40
30
20
10

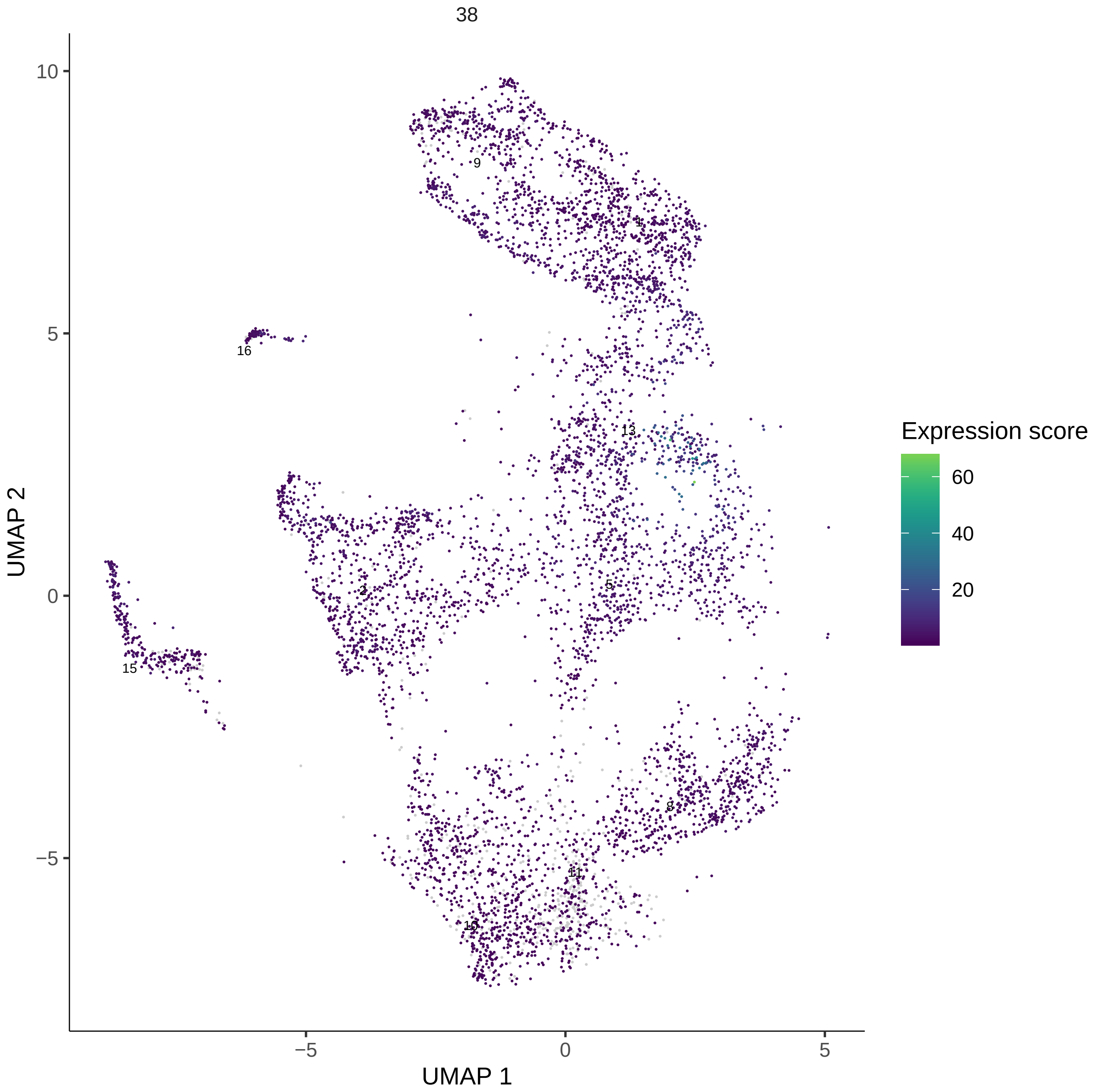
38
10
5
0
−5
UMAP 2
−5
0
5
UMAP 1
9
1
16
13
2
5
15
8
11
10
Expression score
60
40
20

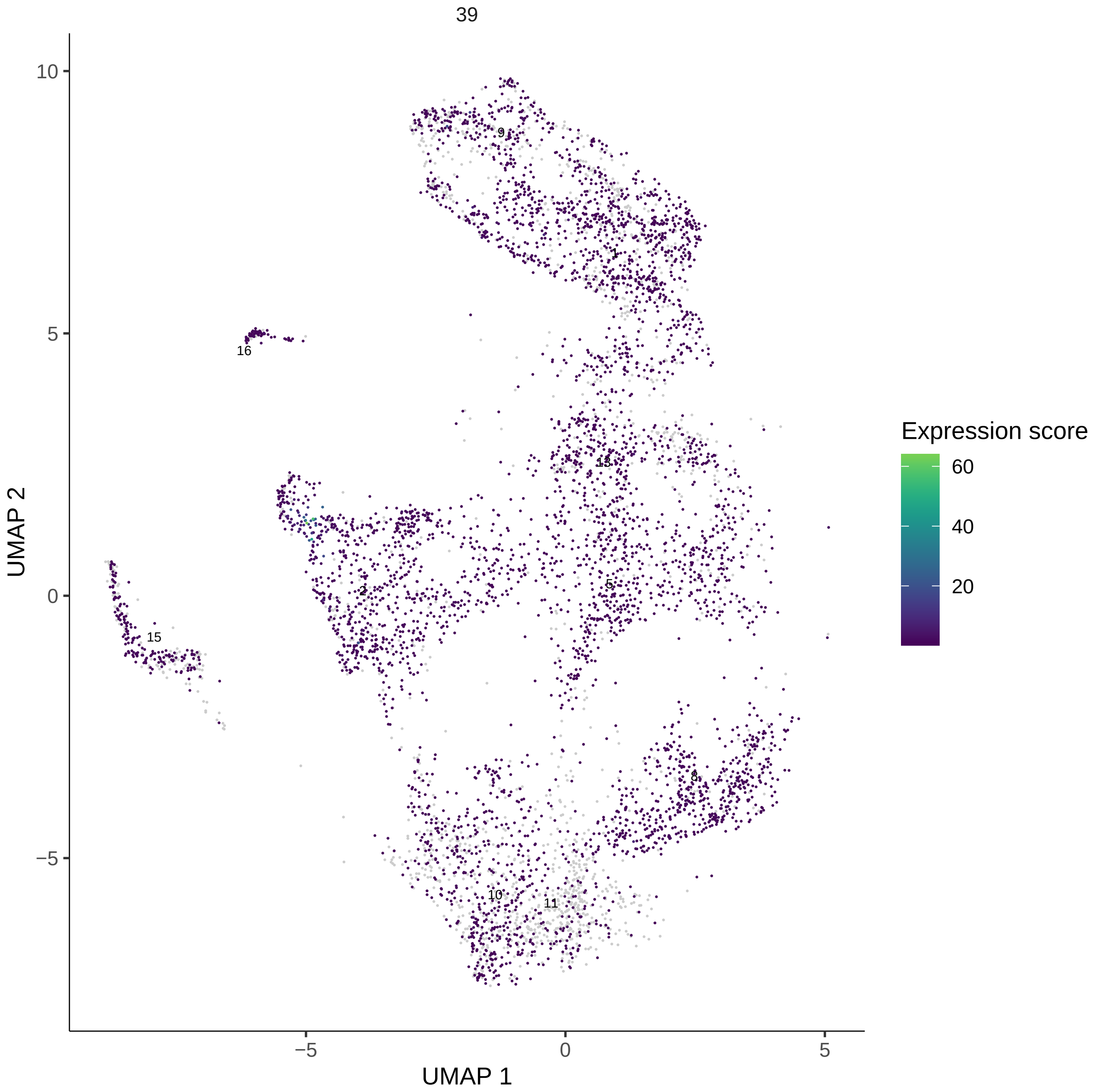
39
UMAP 2
UMAP 1
10
5
0
−5
−5
0
5
9
1
16
13
15
2
5
8
10
11
Expression score
60
40
20

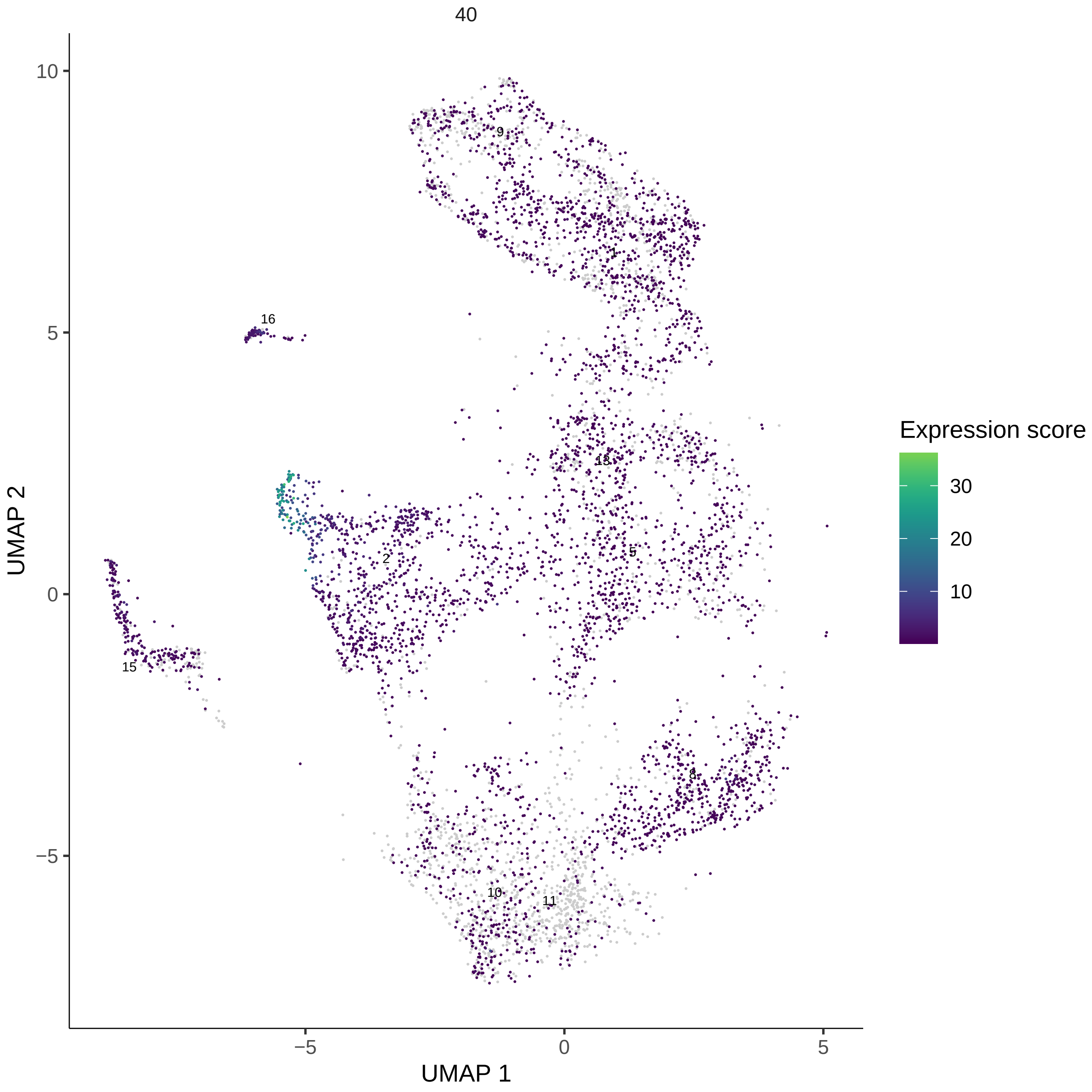
40
UMAP 2
UMAP 1
10
5
0
−5
−5
0
5
Expression score
30
20
10
9
1
16
13
2
5
15
8
10
11

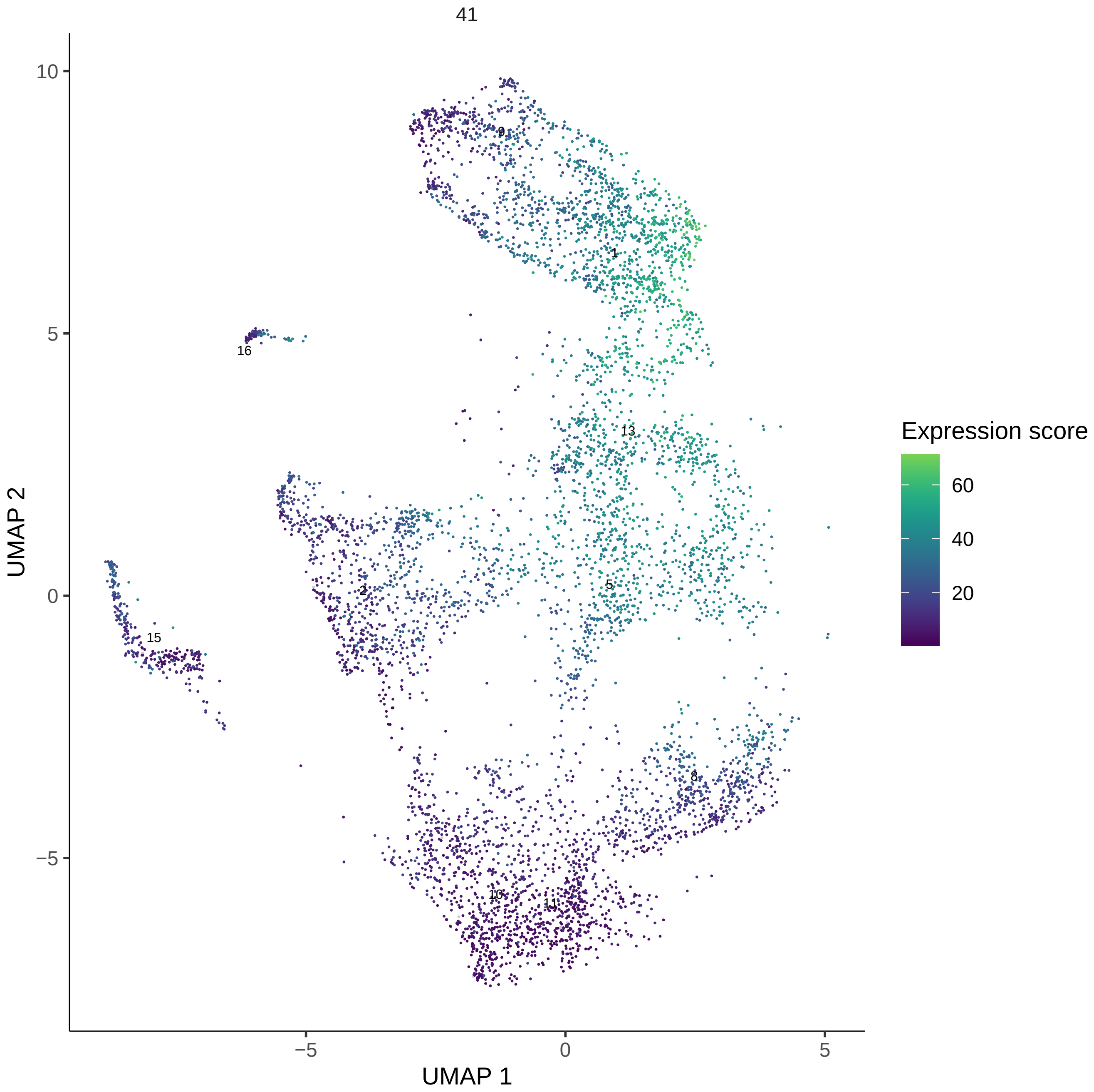
41
UMAP 1
UMAP 2
Expression score
60
40
20
10
5
0
−5
9
1
16
13
15
2
5
8
10
11

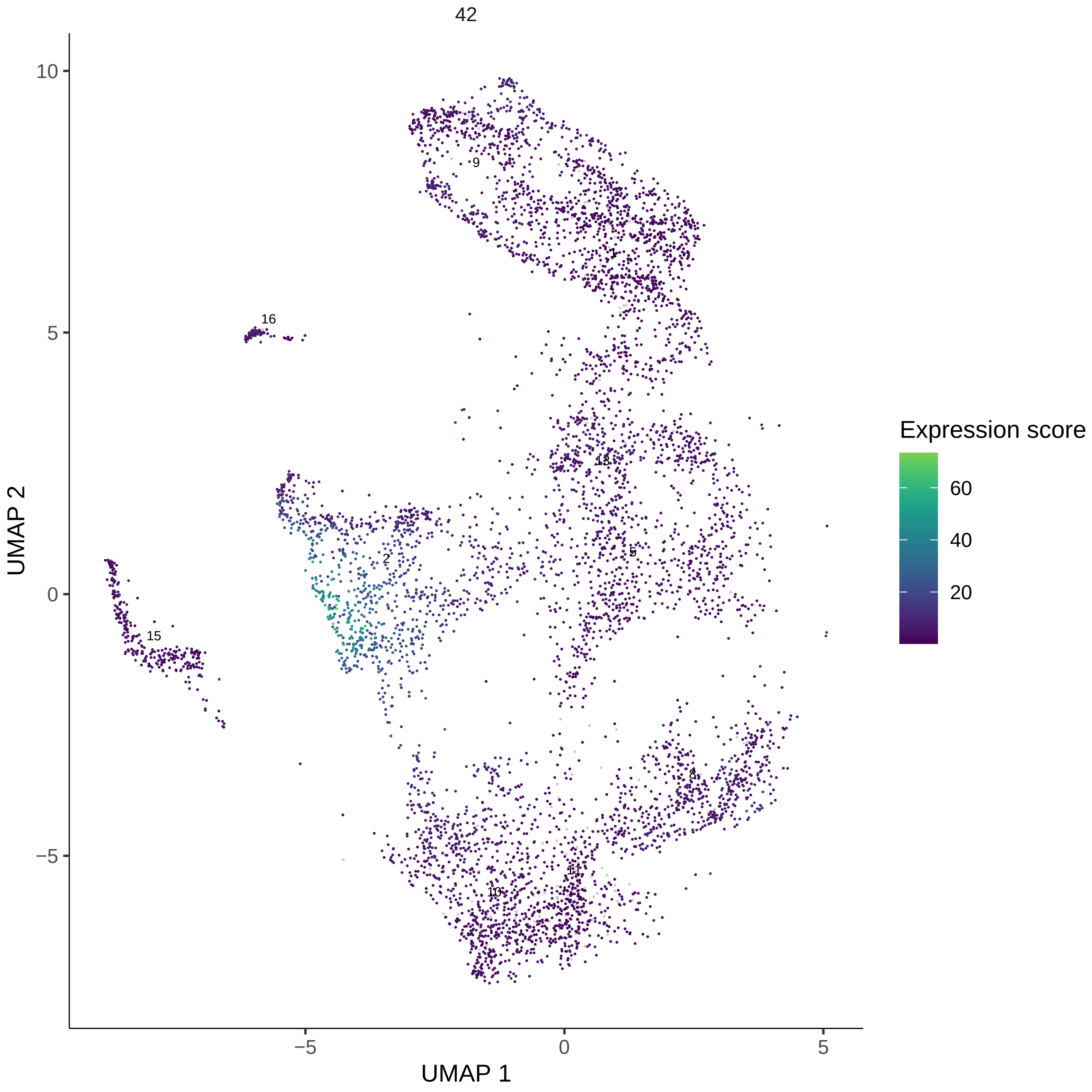

42
UMAP 2
UMAP 1
10
5
0
−5
−5
0
5
9
1
16
13
2
5
15
8
11
10
Expression score
60
40
20

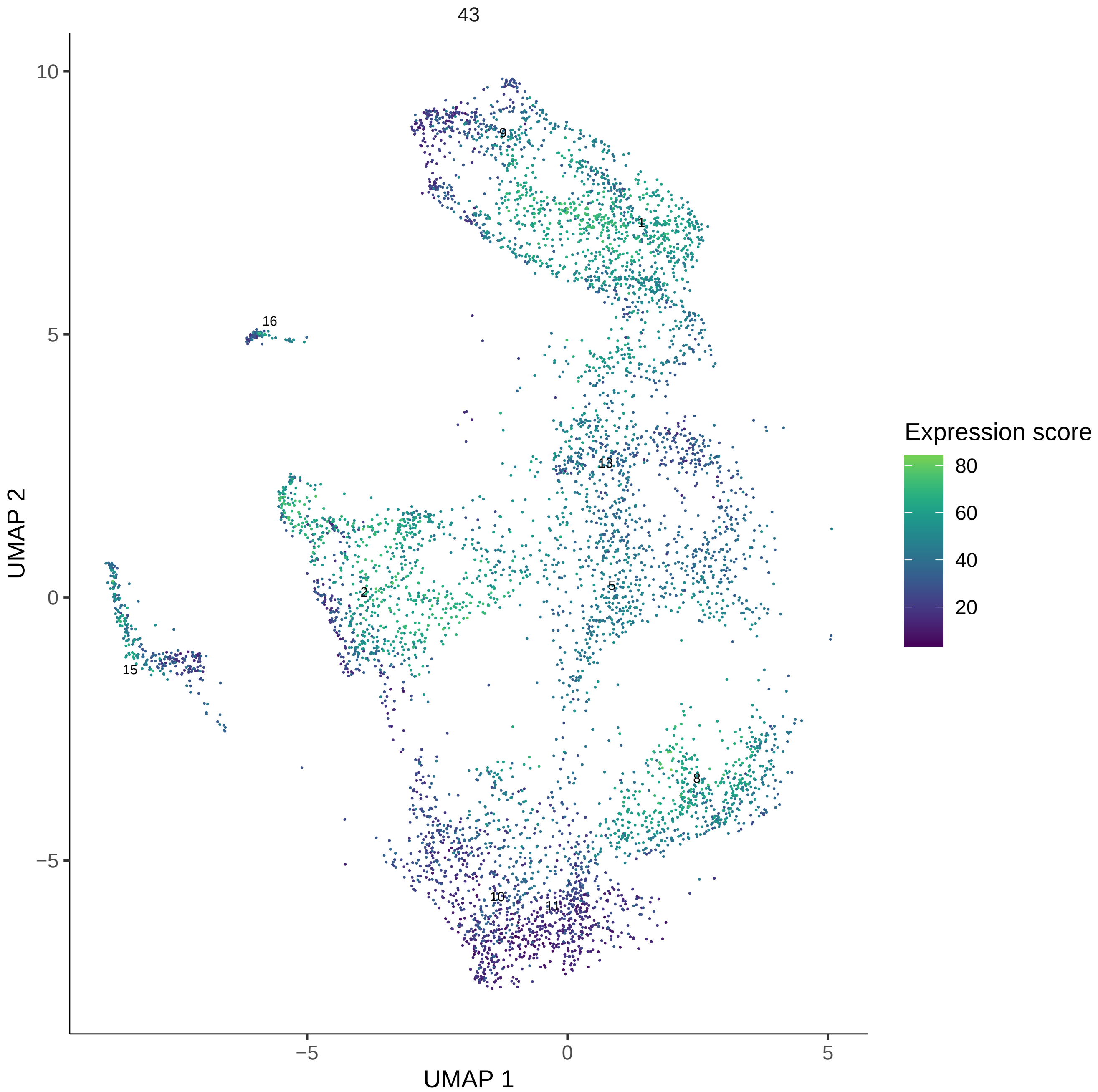

43
UMAP 1
UMAP 2
Expression score
80
60
40
20
10
5
0
−5
5
0
−5
9
1
16
13
2
5
15
8
10
11

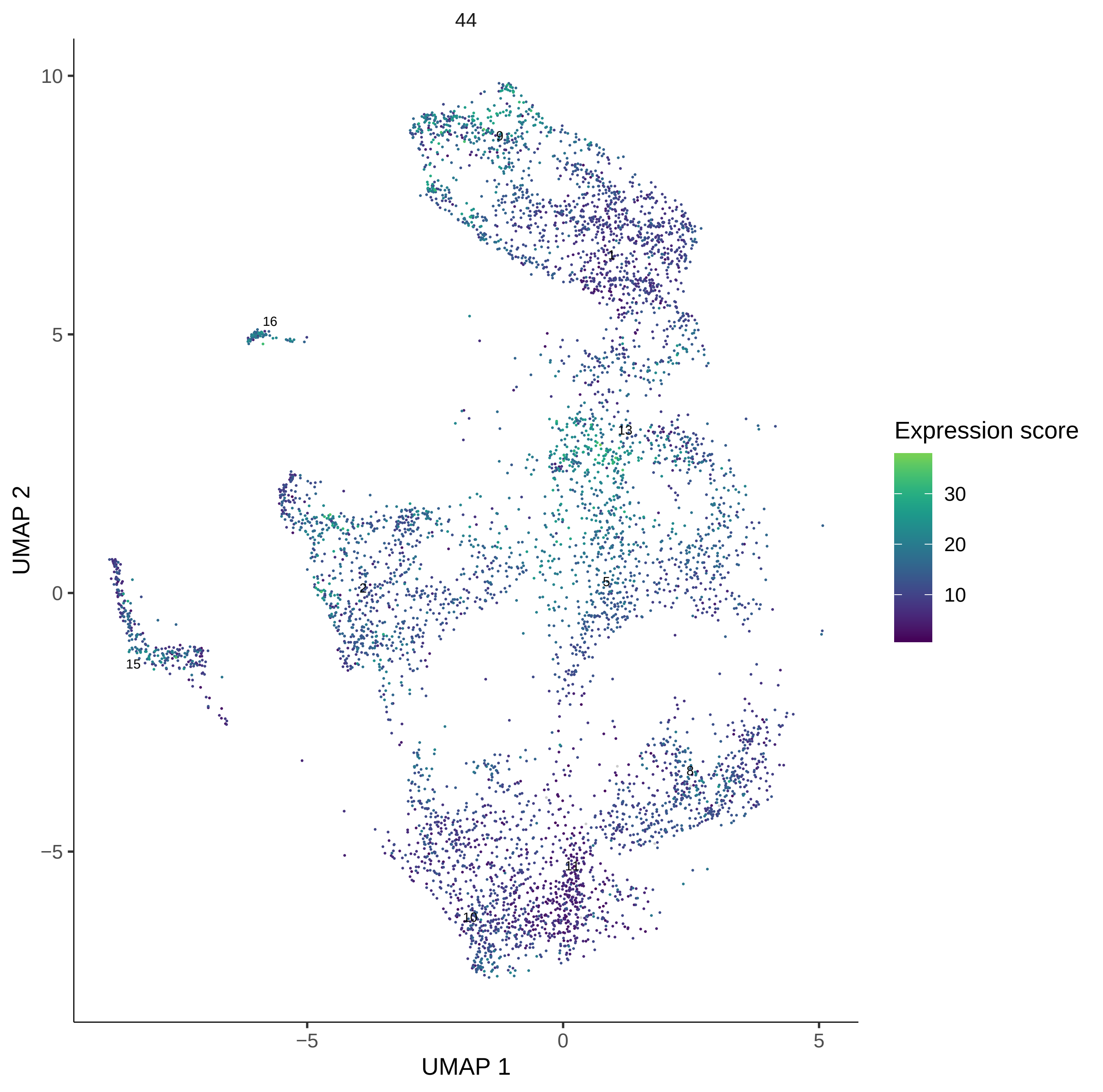

44
UMAP 2
UMAP 1
10
5
0
−5
−5
0
5
9
1
16
13
2
5
15
8
11
10
Expression score
30
20
10

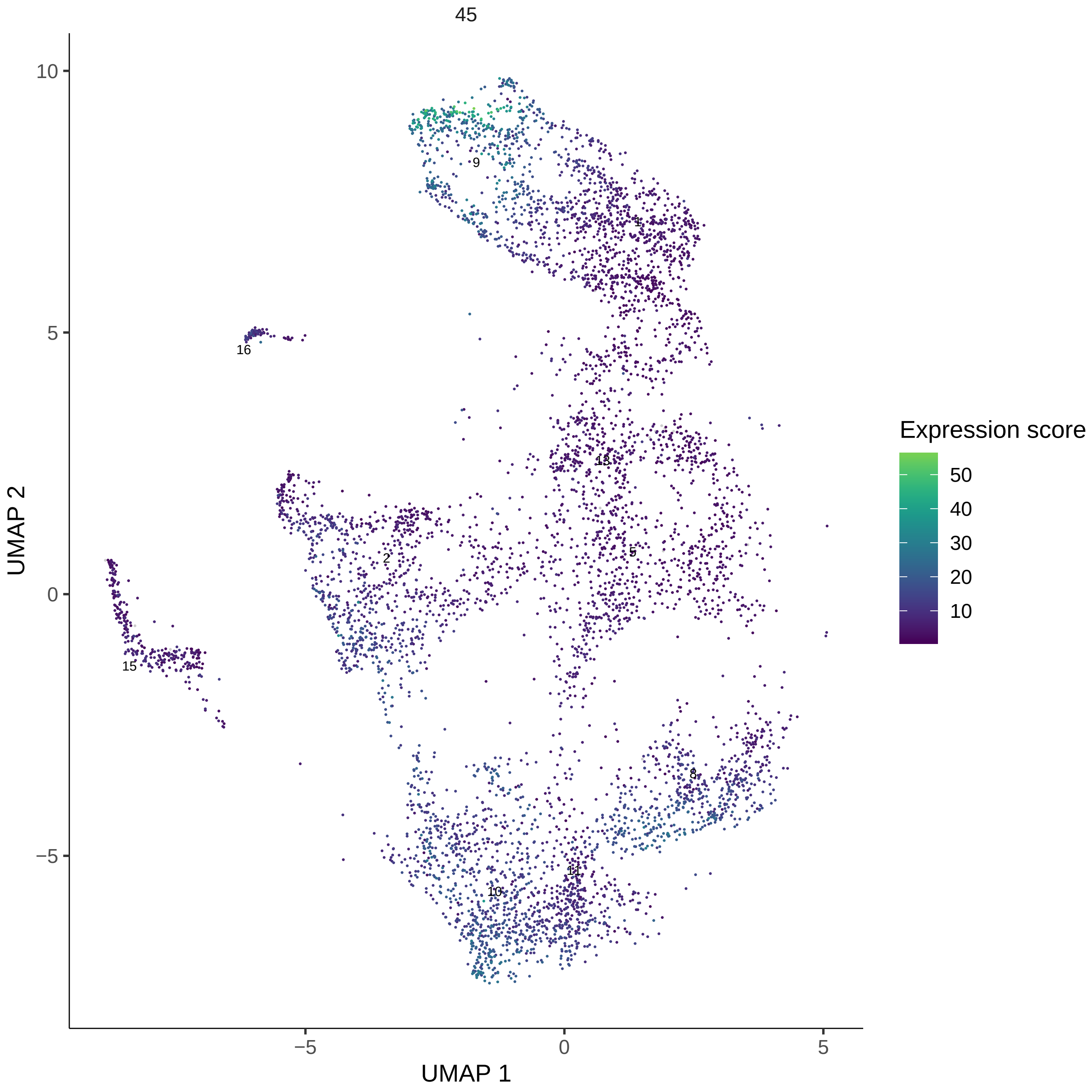

45
10
5
0
−5
−5
0
5
UMAP 1
UMAP 2
9
1
16
13
2
5
15
8
11
10
Expression score
50
40
30
20
10

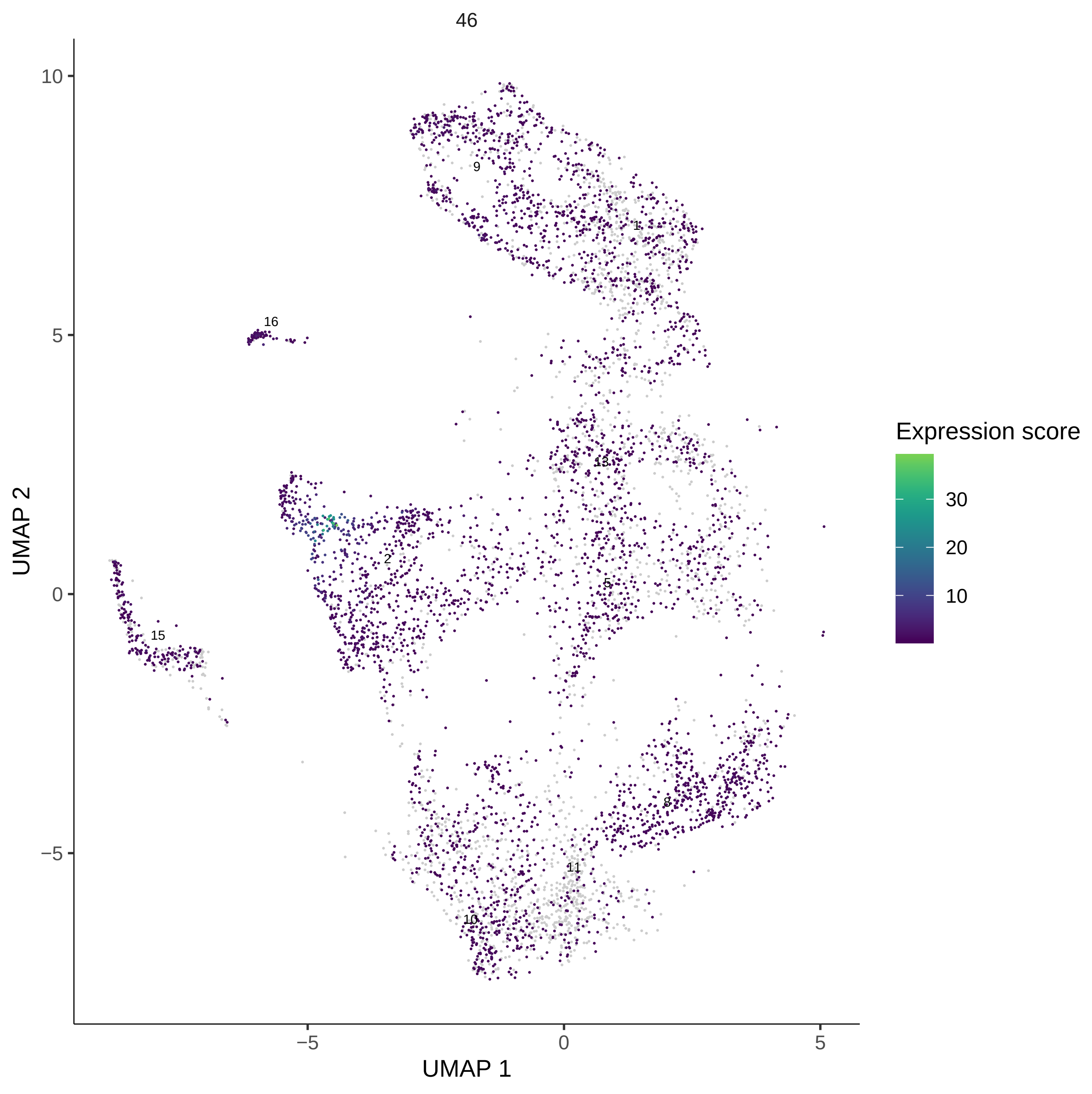

46
UMAP 2
UMAP 1
10
5
0
−5
−5
0
5
9
1
16
13
2
5
15
8
11
10
Expression score
30
20
10

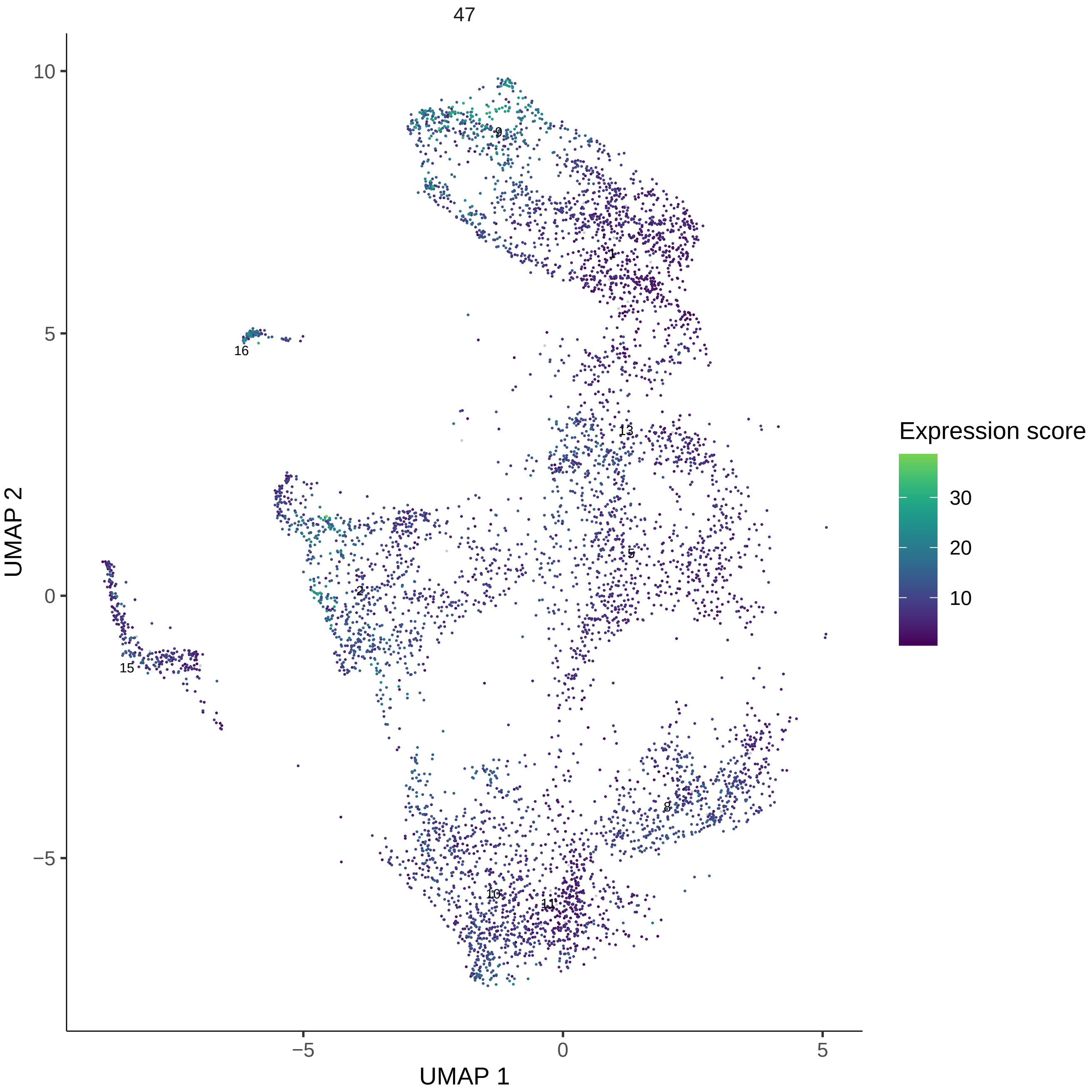
47
10
5
0
−5
−5
0
5
UMAP 1
UMAP 2
9
1
16
13
2
5
15
8
10
11
Expression score
30
20
10

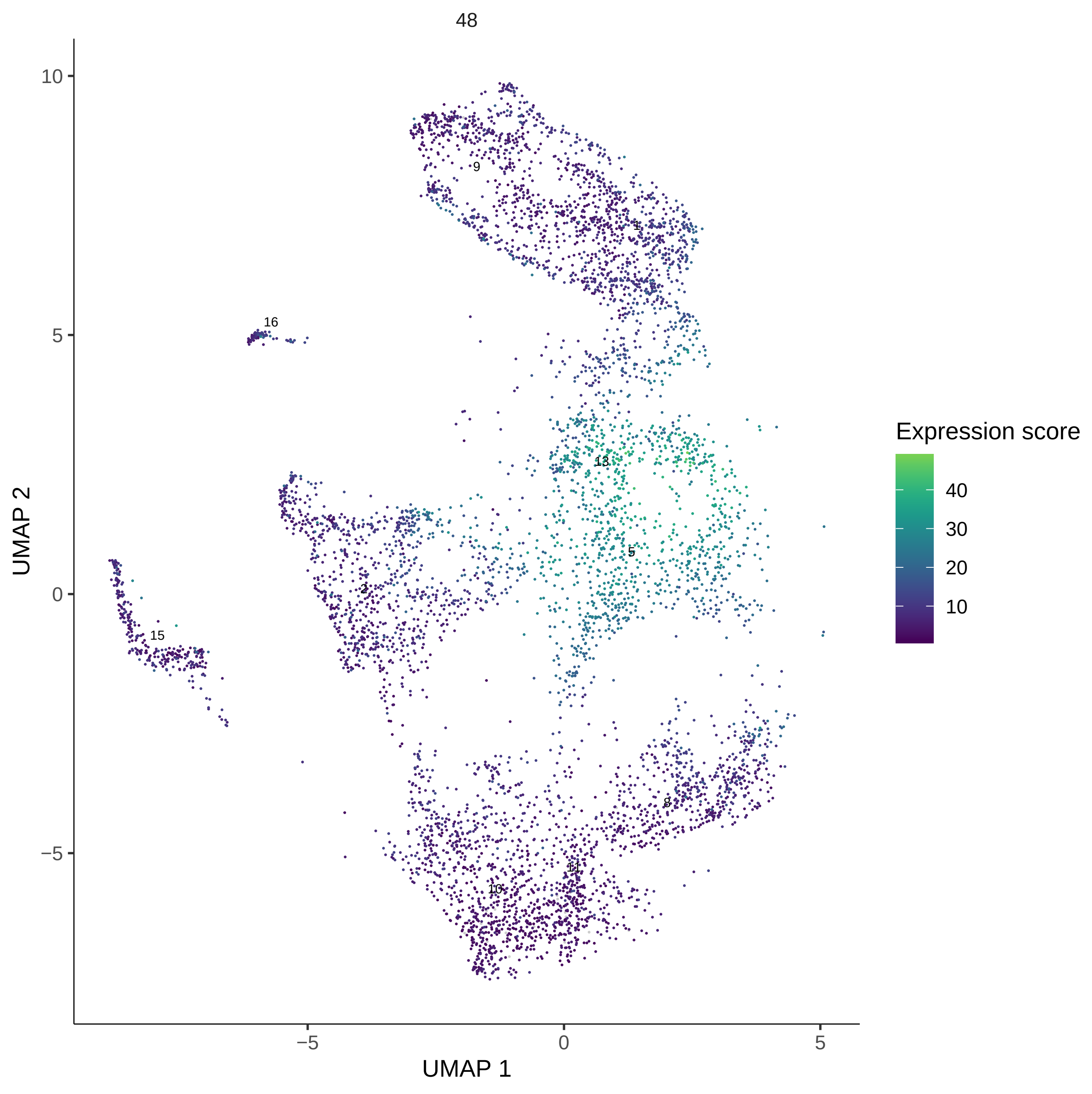
48
UMAP 1
UMAP 2
Expression score
40
30
20
10
9
1
16
13
5
2
15
8
11
10
10
5
0
−5
−5
0
5

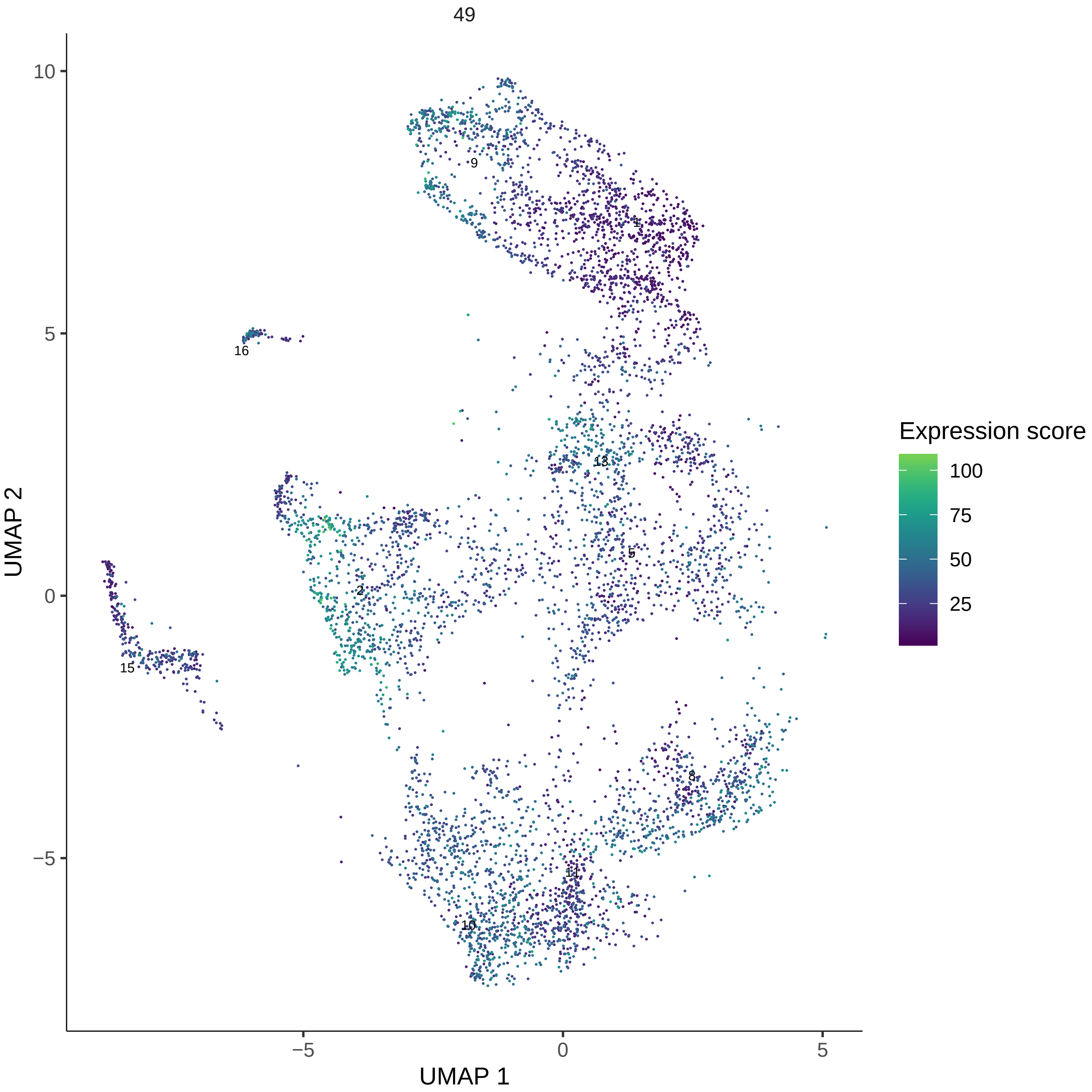
49
Expression score
100
75
50
25
UMAP 2
UMAP 1
10
5
0
−5
9
1
16
13
5
2
15
8
11
10

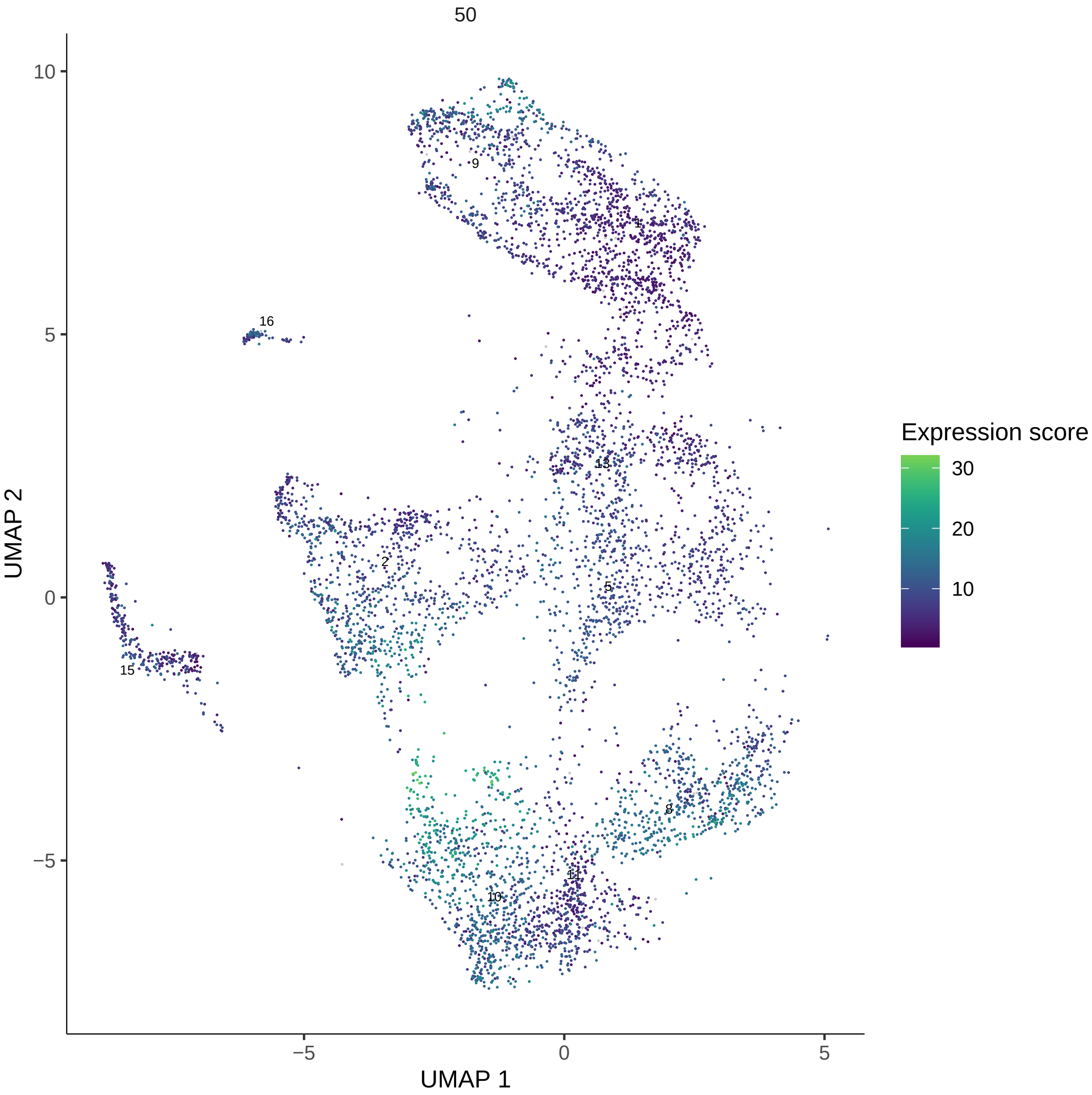

50
UMAP 2
UMAP 1
10
5
0
−5
−5
0
5
9
1
16
13
2
5
15
8
11
10
Expression score
30
20
10

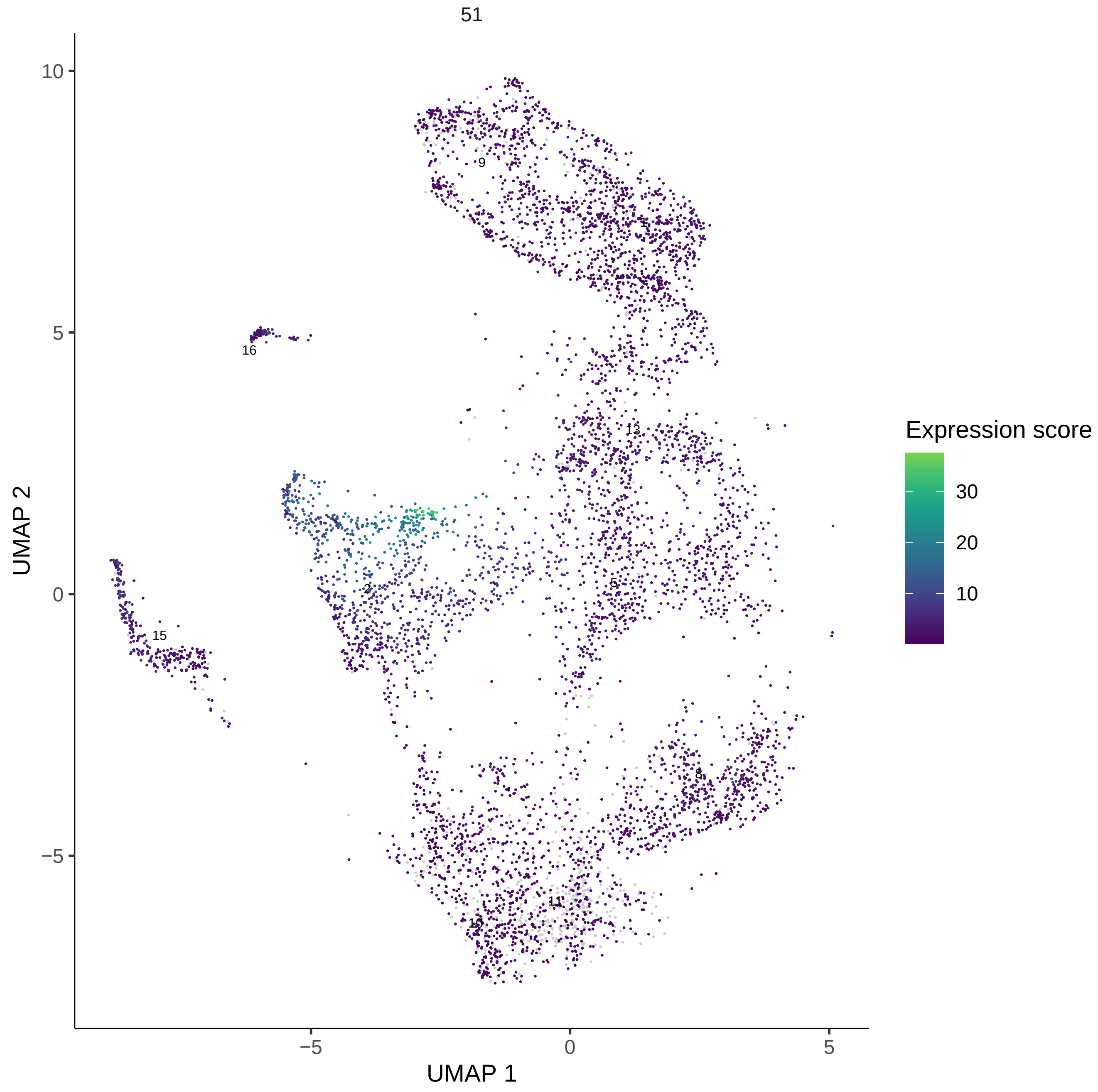

51
UMAP 2
UMAP 1
10
5
0
−5
−5
0
5
9
1
16
13
2
5
15
8
11
10
Expression score
30
20
10

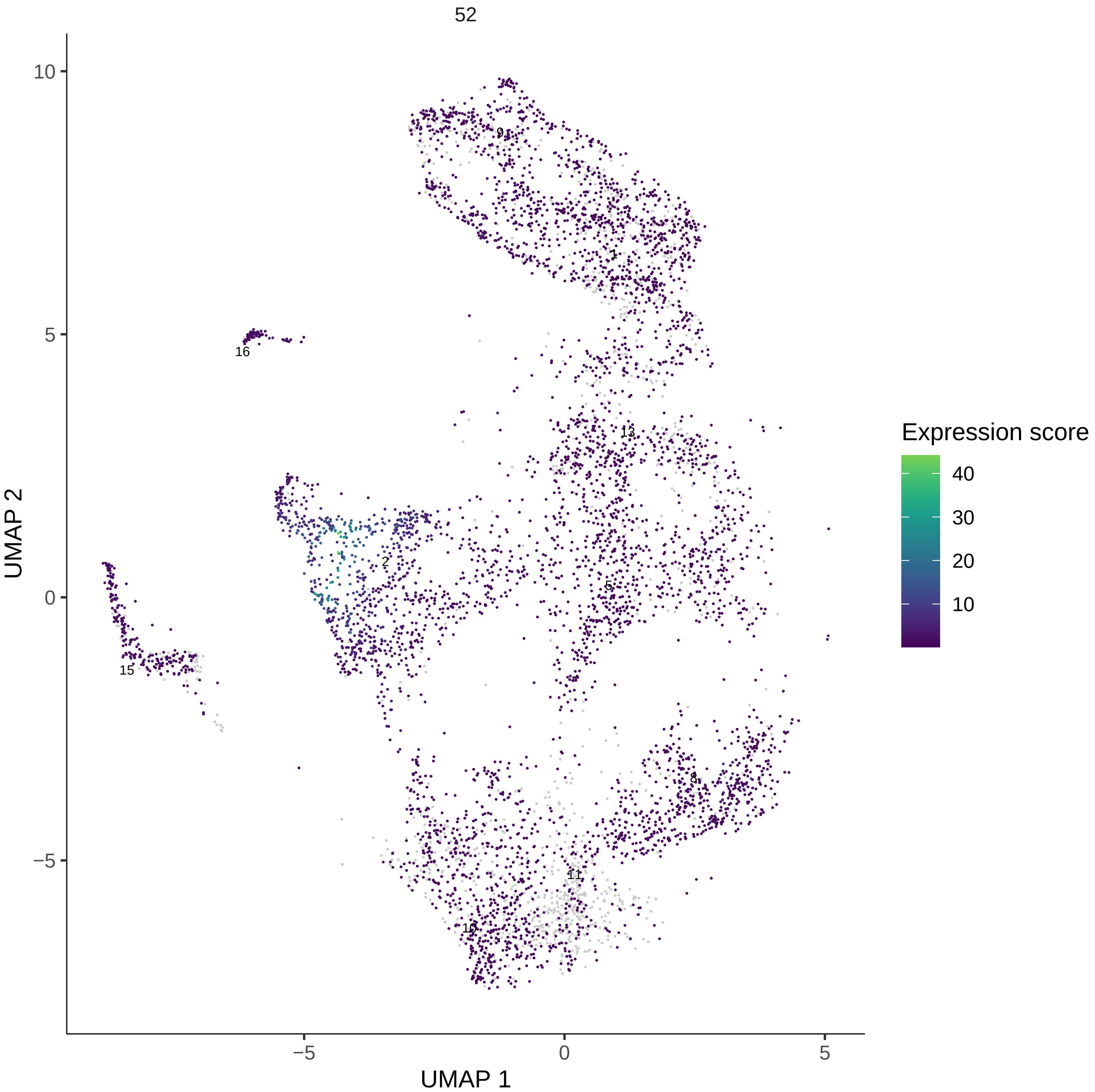
52
UMAP 1
UMAP 2
Expression score
40
30
20
10
−5
0
5
10
9
1
16
13
2
5
15
8
11
10

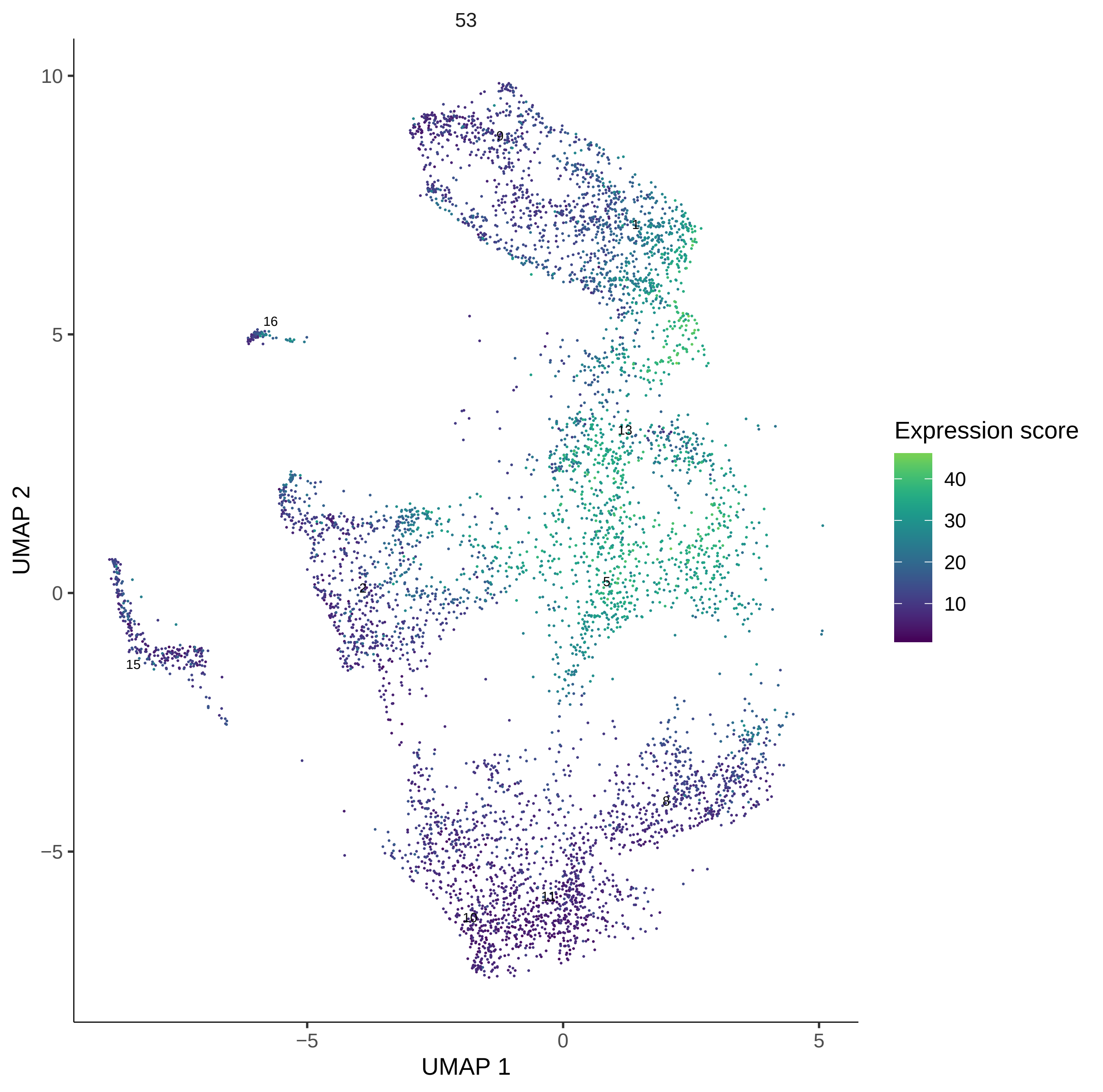
53
UMAP 1
UMAP 2
Expression score
40
30
20
10
−5
0
5
10
9
1
16
13
2
5
15
8
11
10

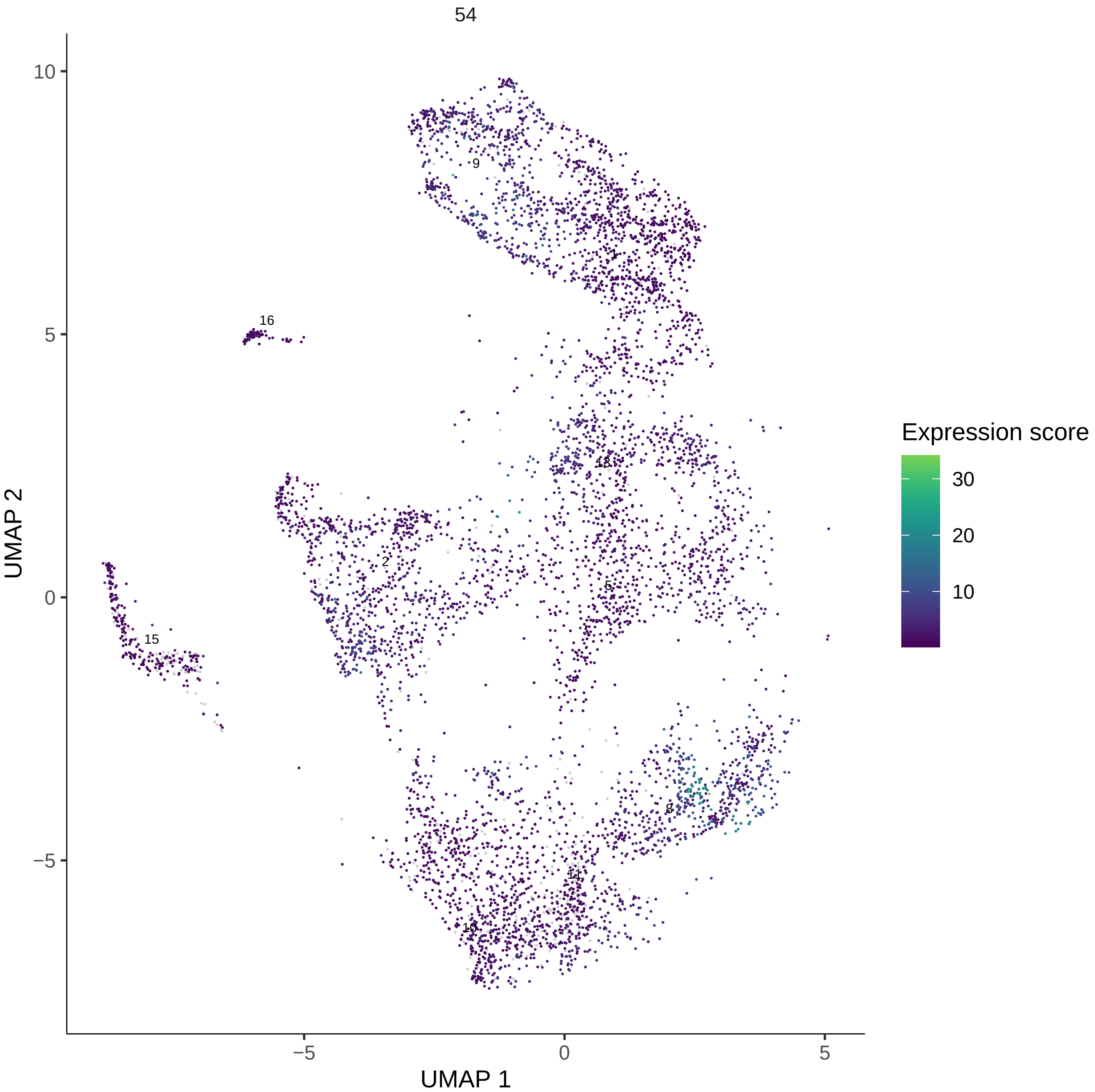

54
Expression score
30
20
10
UMAP 1
UMAP 2
10
5
0
−5
−5
0
5
9
1
16
13
2
5
15
8
11
10

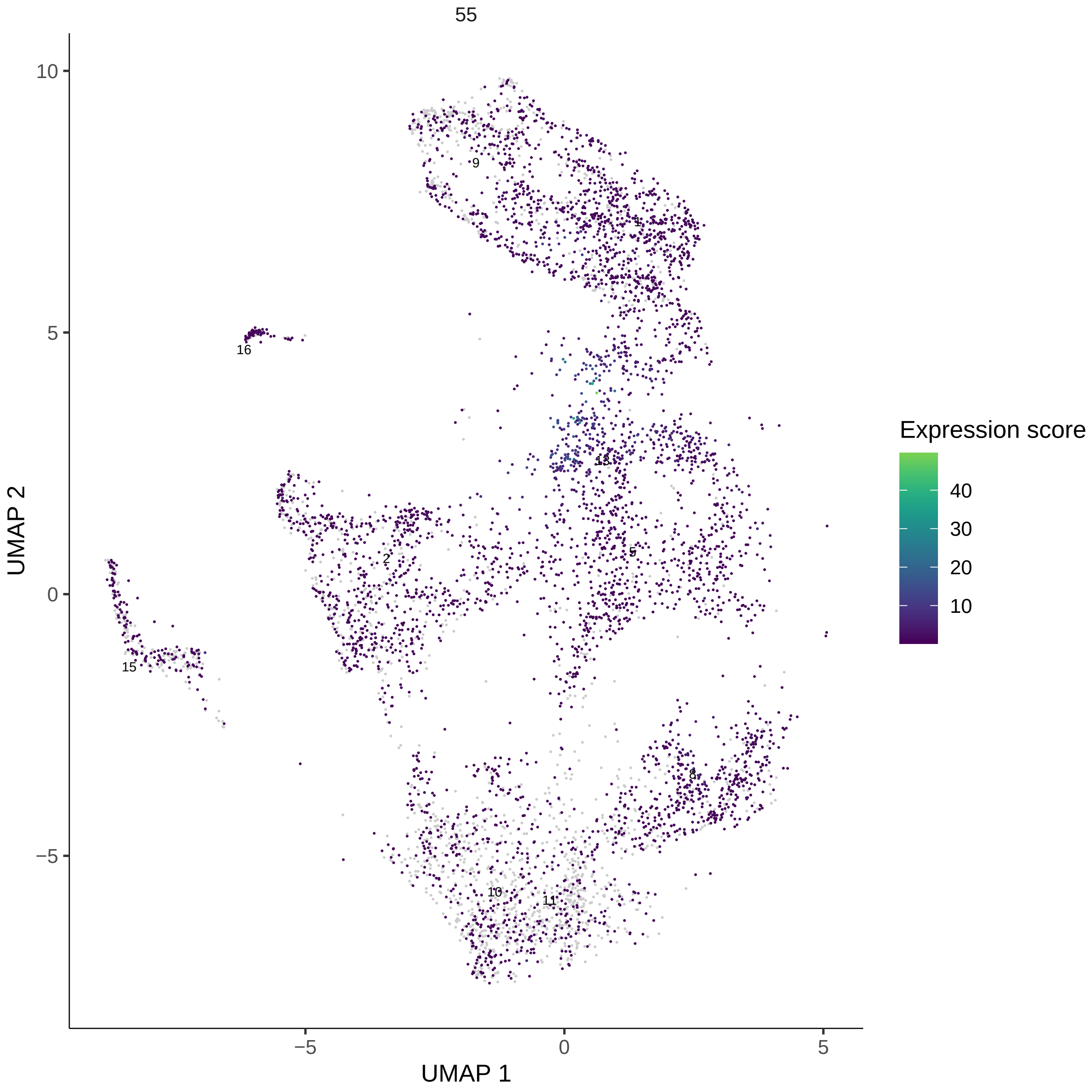
55
UMAP 1
UMAP 2
Expression score
40
30
20
10
9
1
16
13
2
5
15
8
10
11
−5
0
5
10

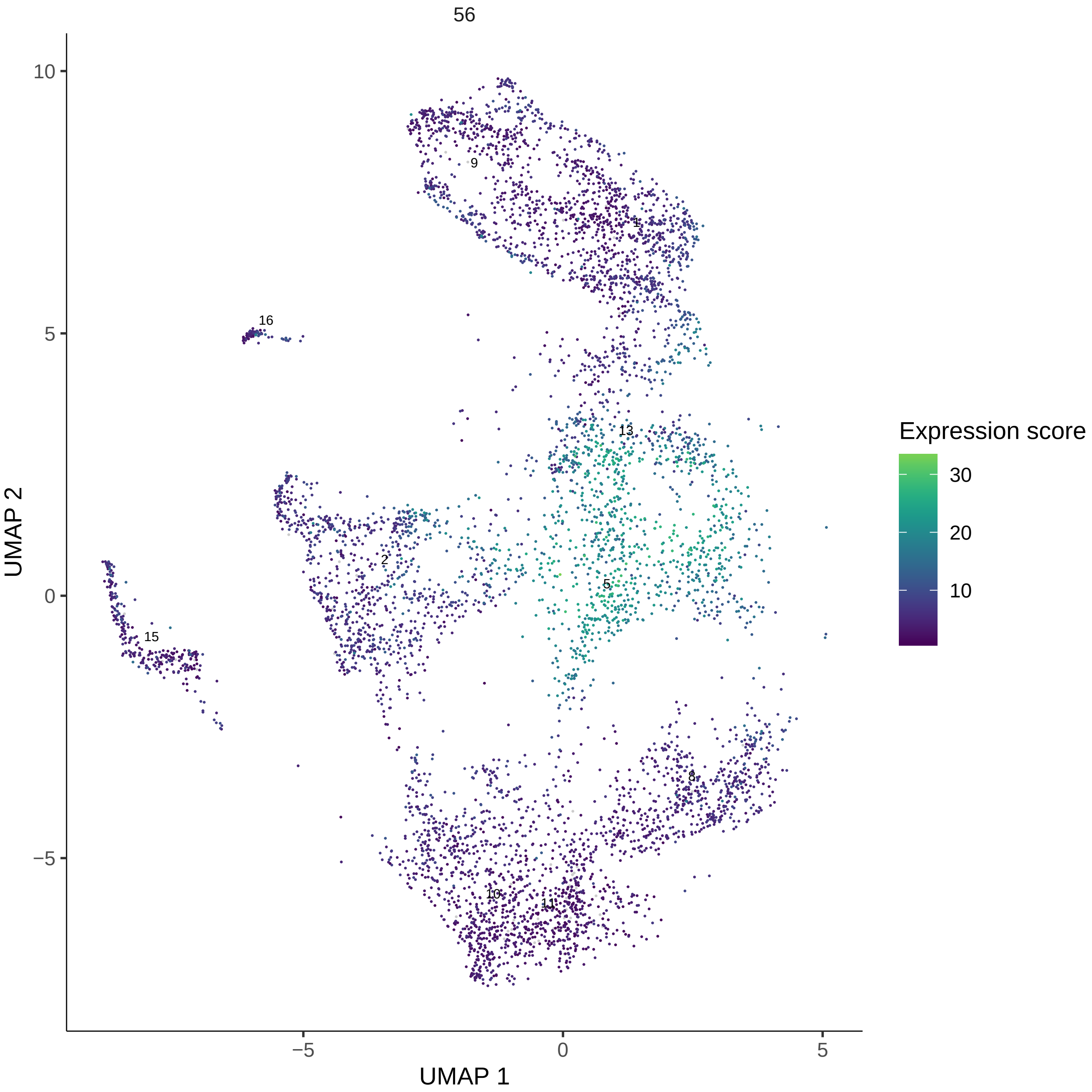

56
Expression score
30
20
10
UMAP 2
UMAP 1
10
5
0
−5
−5
0
5
9
1
16
13
2
5
15
8
10
11

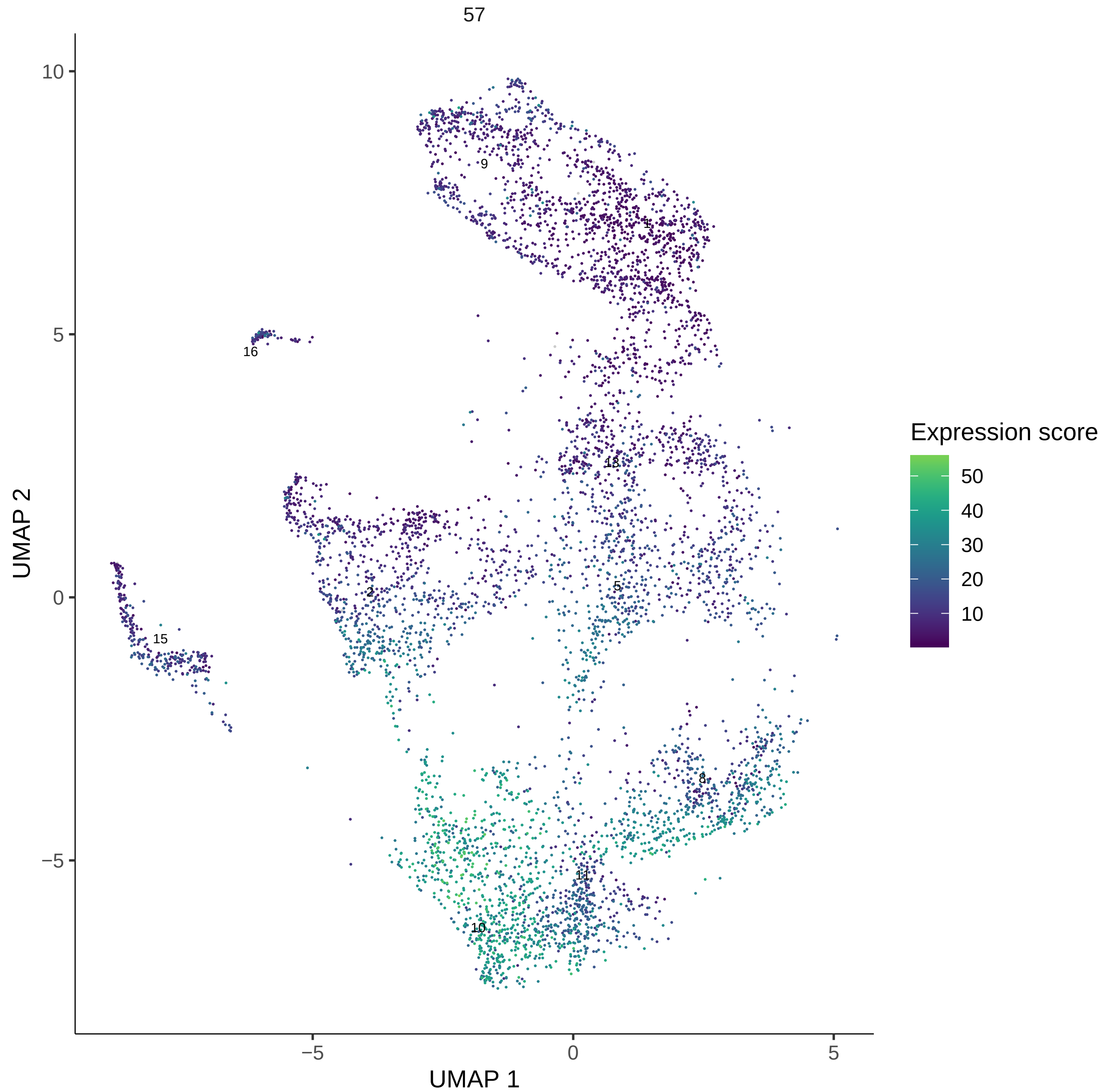

57
Expression score
50
40
30
20
10
UMAP 2
UMAP 1
10
5
0
−5
−5
0
5
9
1
16
13
2
5
15
8
11
10

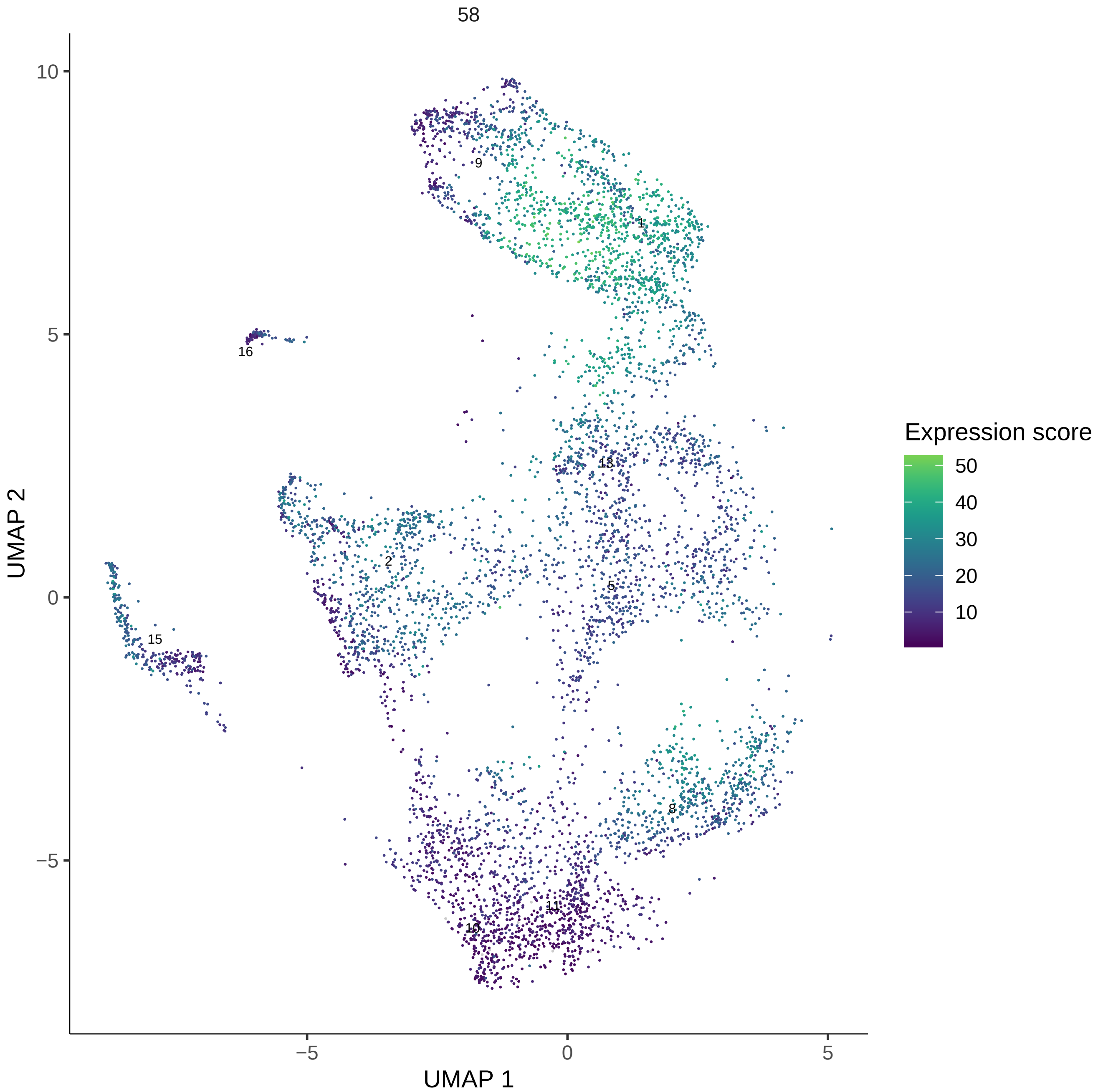

58
UMAP 1
UMAP 2
Expression score
50
40
30
20
10
10
5
0
−5
−5
0
5
9
1
16
13
2
5
15
8
11
10

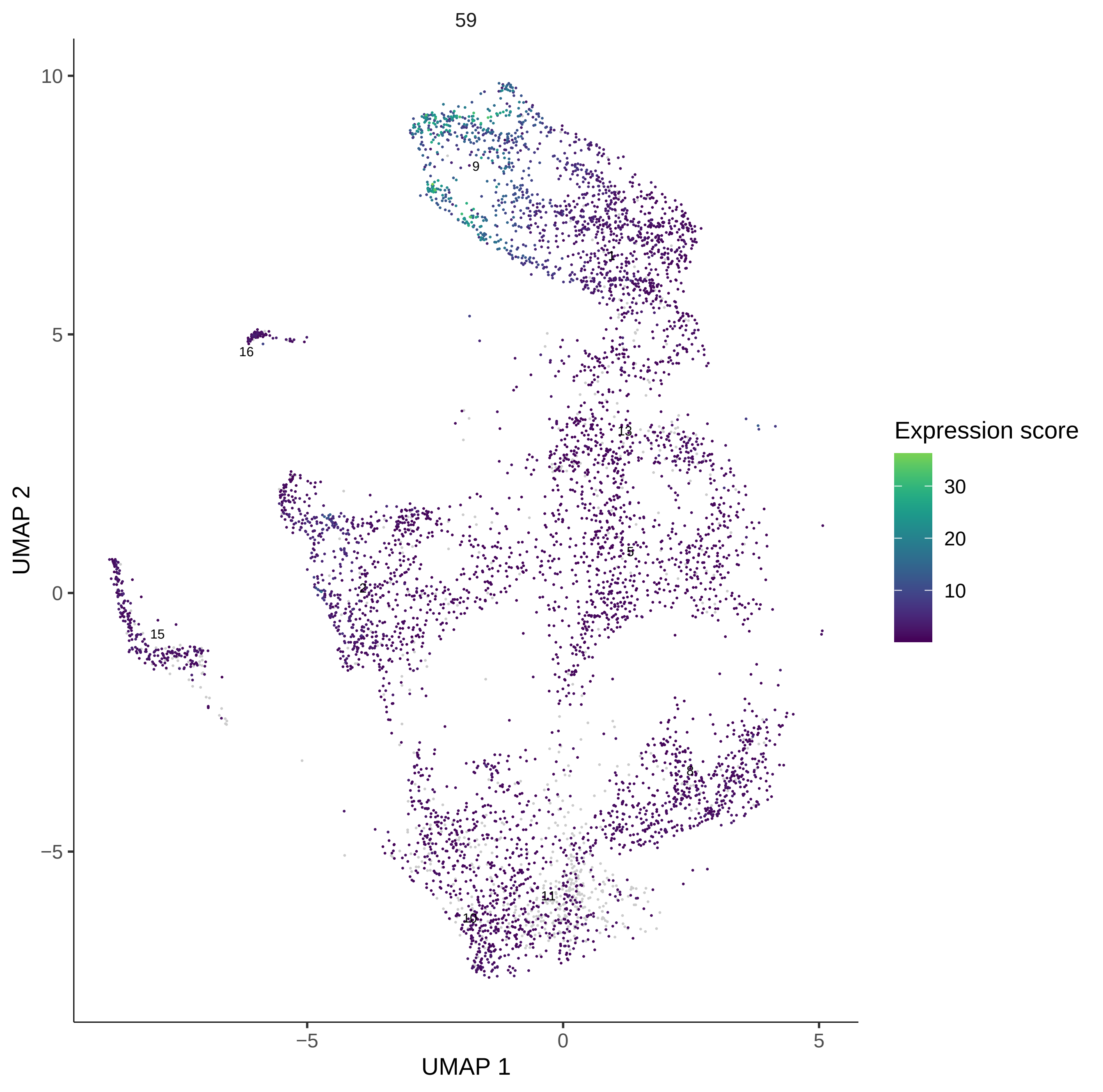

59
UMAP 2
UMAP 1
10
5
0
−5
−5
0
5
9
1
16
13
15
2
5
8
11
10
Expression score
30
20
10

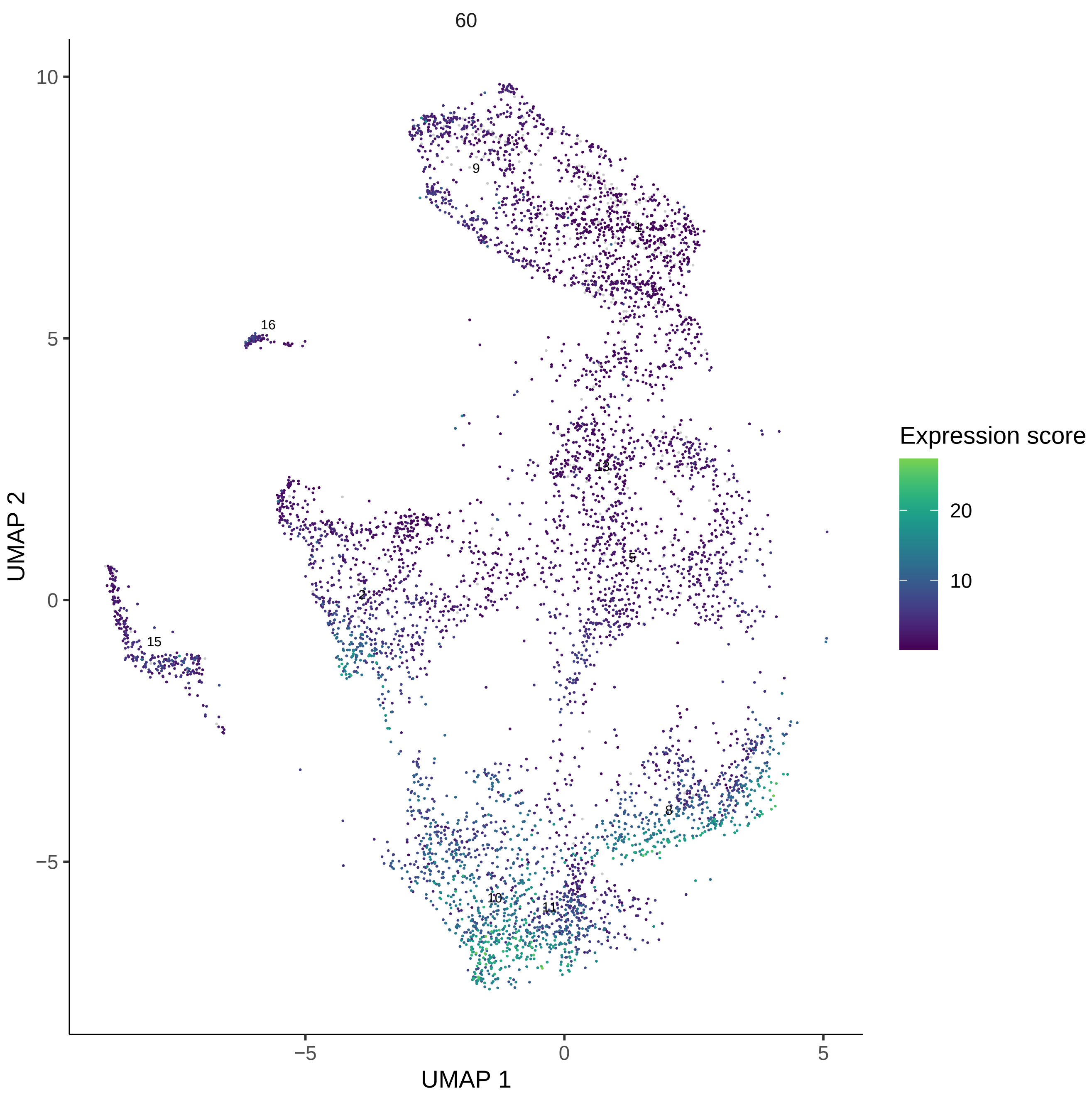
60
UMAP 1
UMAP 2
Expression score
20
10
10
5
0
−5
−5
0
5
9
1
16
13
5
2
15
8
10
11

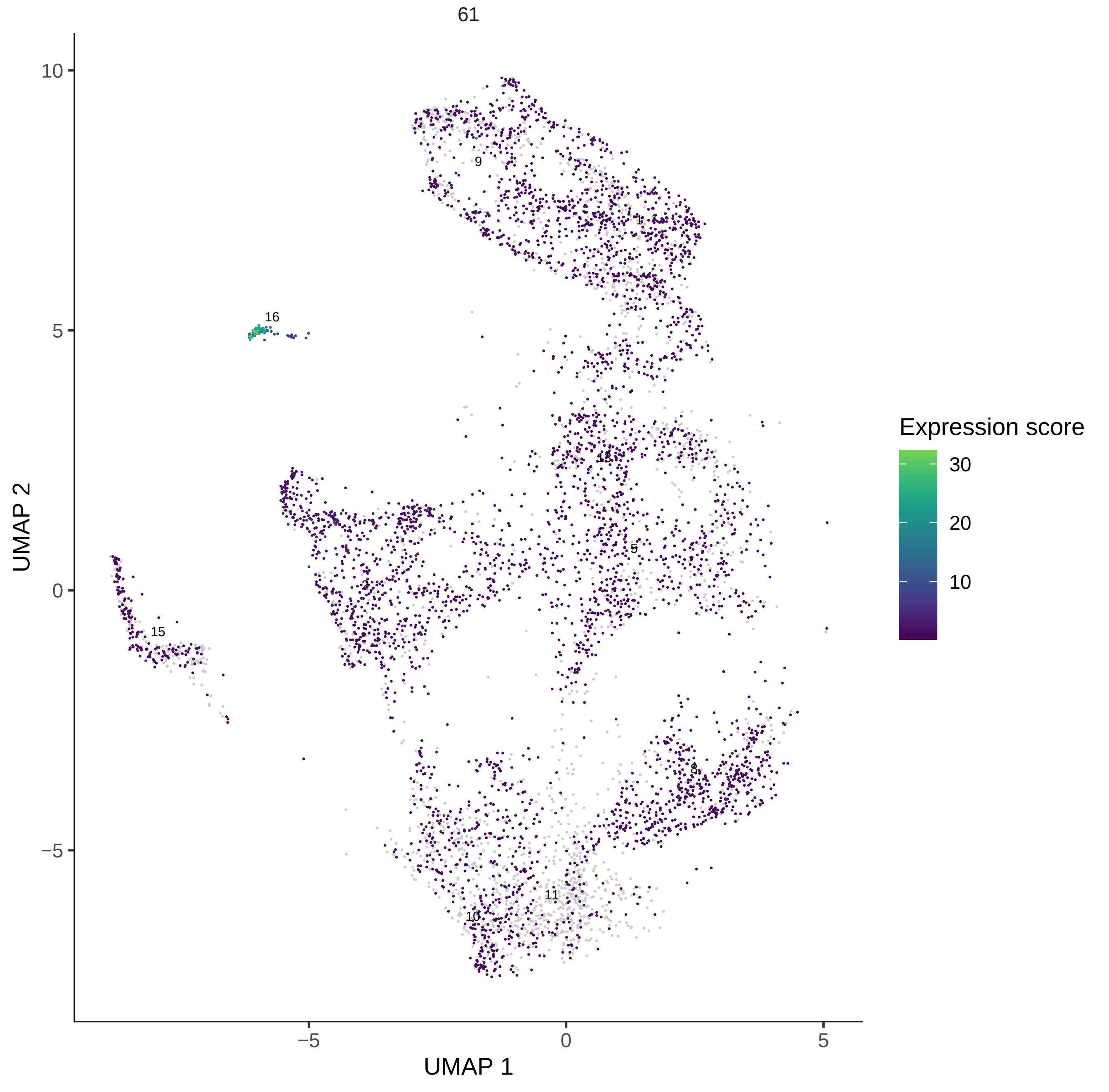
61
UMAP 2
UMAP 1
10
5
0
−5
−5
0
5
9
1
16
13
5
2
15
8
11
10
Expression score
30
20
10

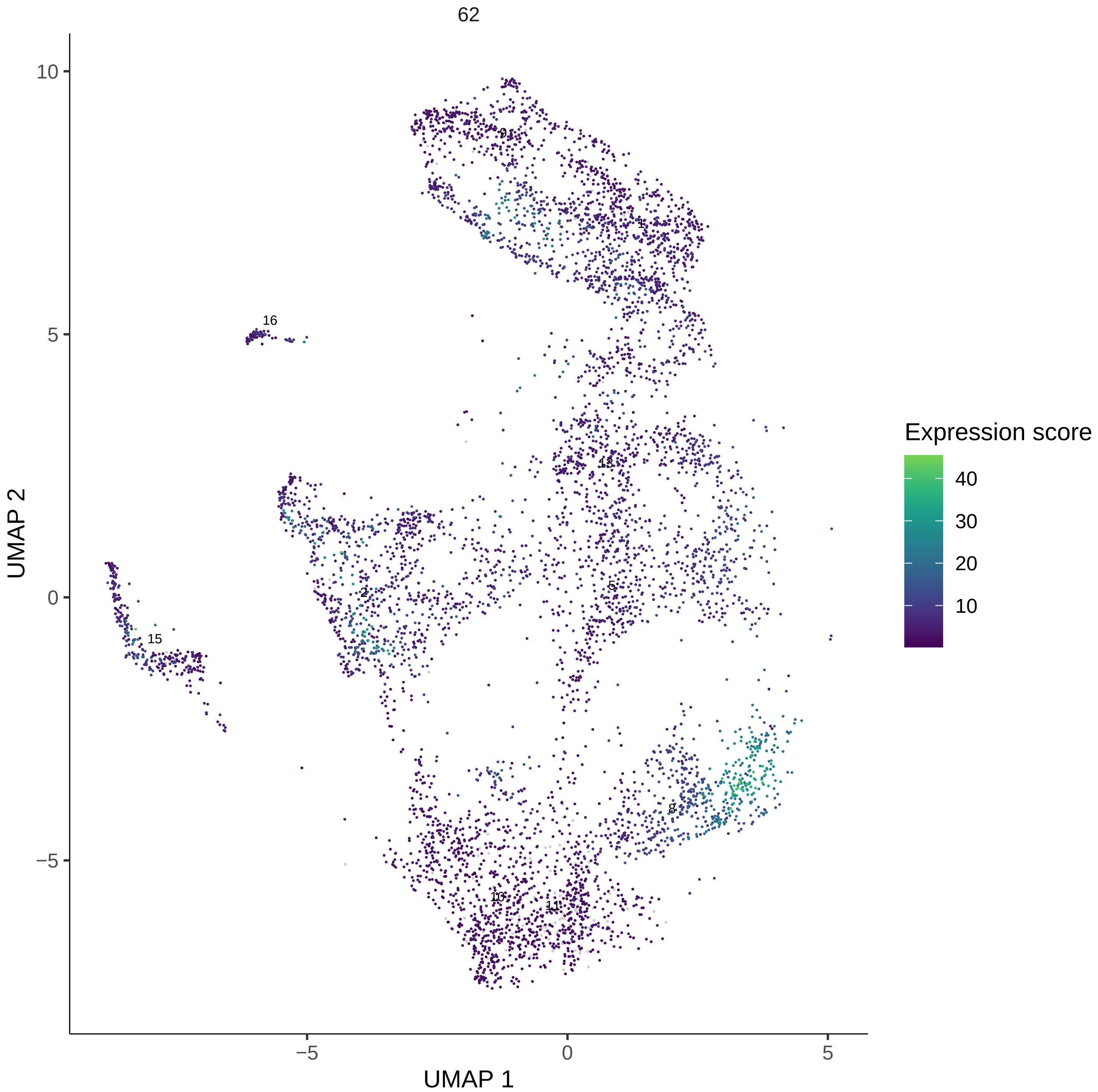

62
UMAP 2
UMAP 1
10
5
0
−5
−5
0
5
9
1
16
13
2
5
15
8
10
11
Expression score
40
30
20
10

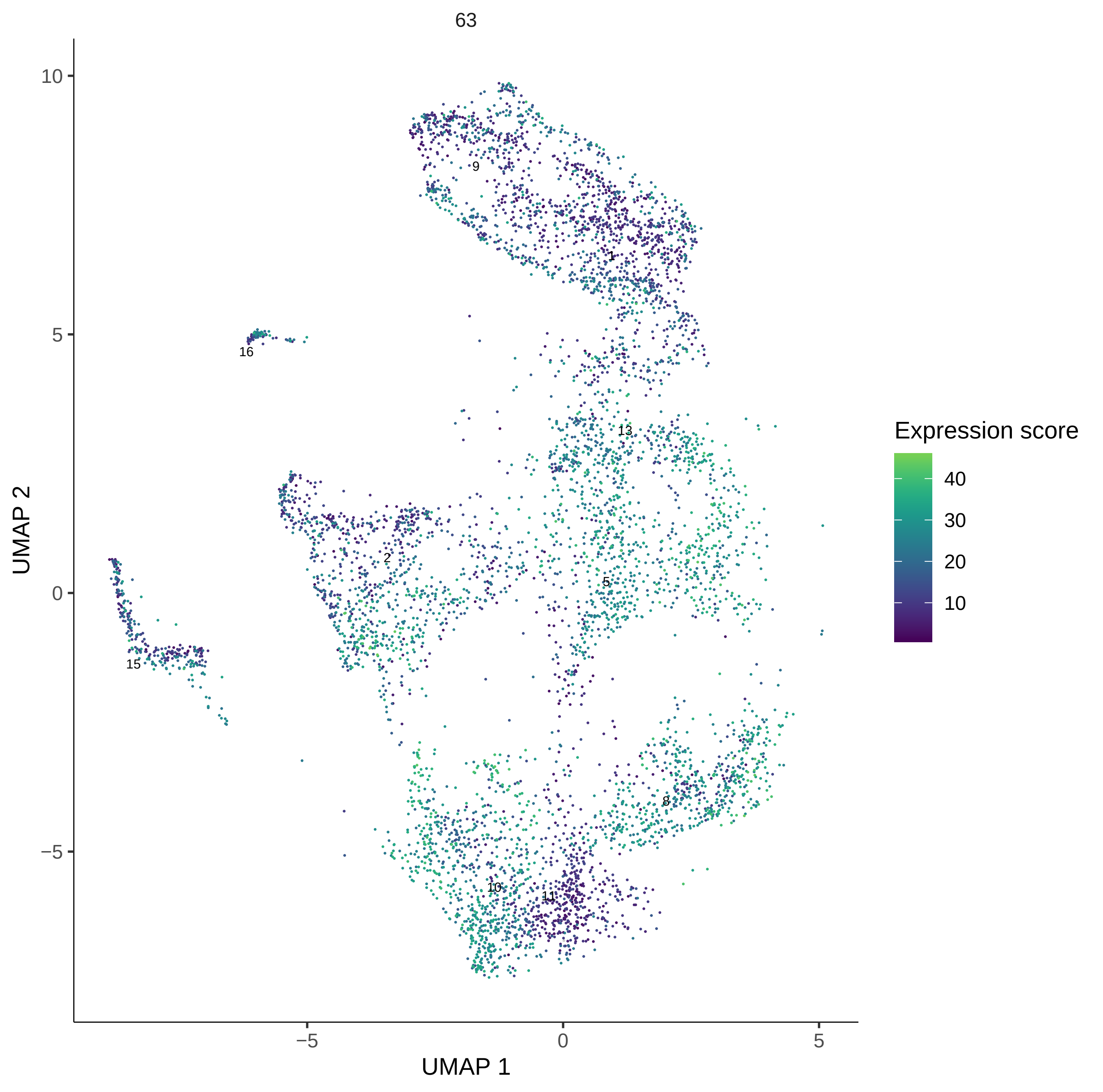

63
UMAP 2
UMAP 1
10
5
0
−5
−5
0
5
9
1
16
13
2
5
15
8
10
11
Expression score
40
30
20
10

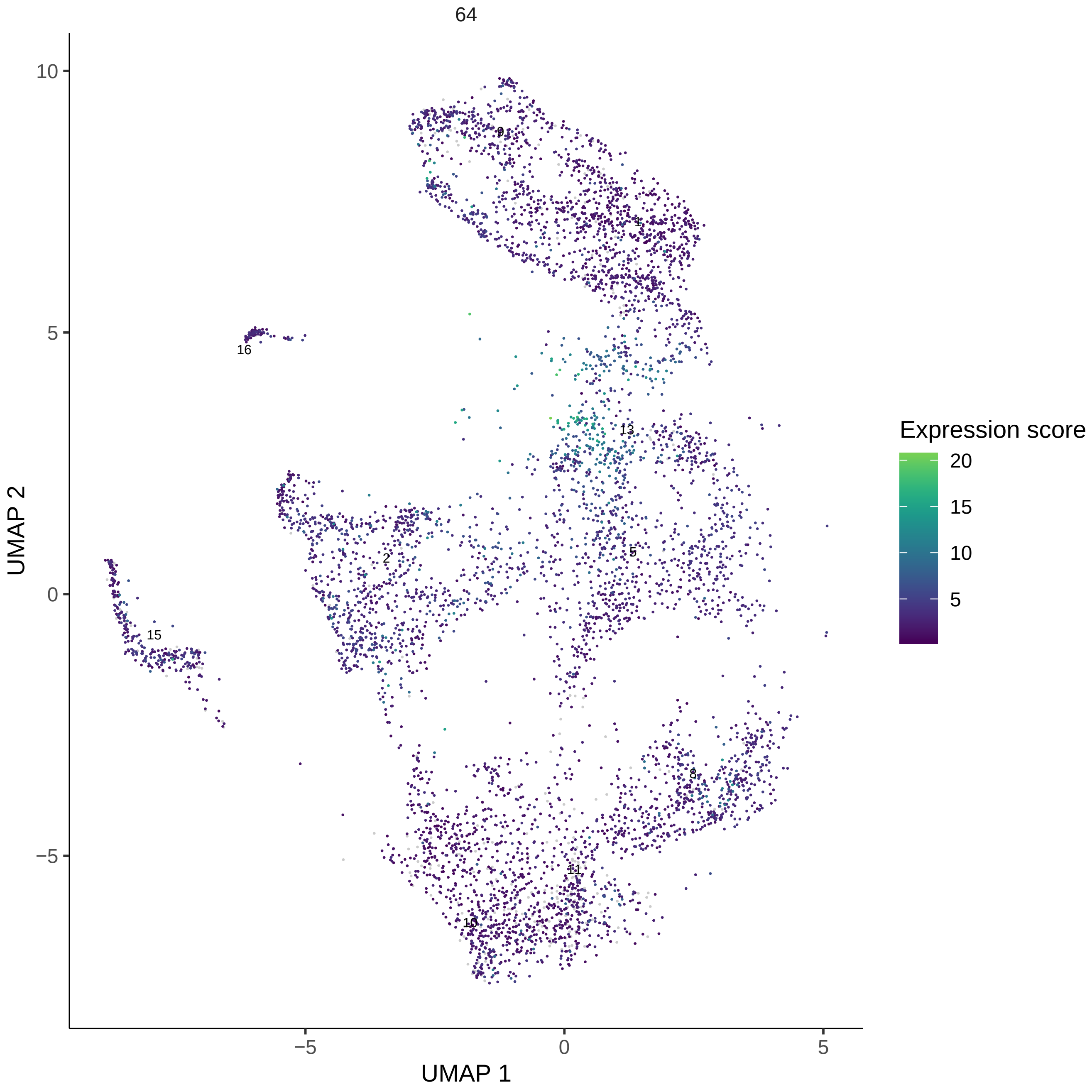

64
10
5
0
−5
−5
0
5
UMAP 1
UMAP 2
9
1
16
13
2
5
15
8
11
10
Expression score
20
15
10
5

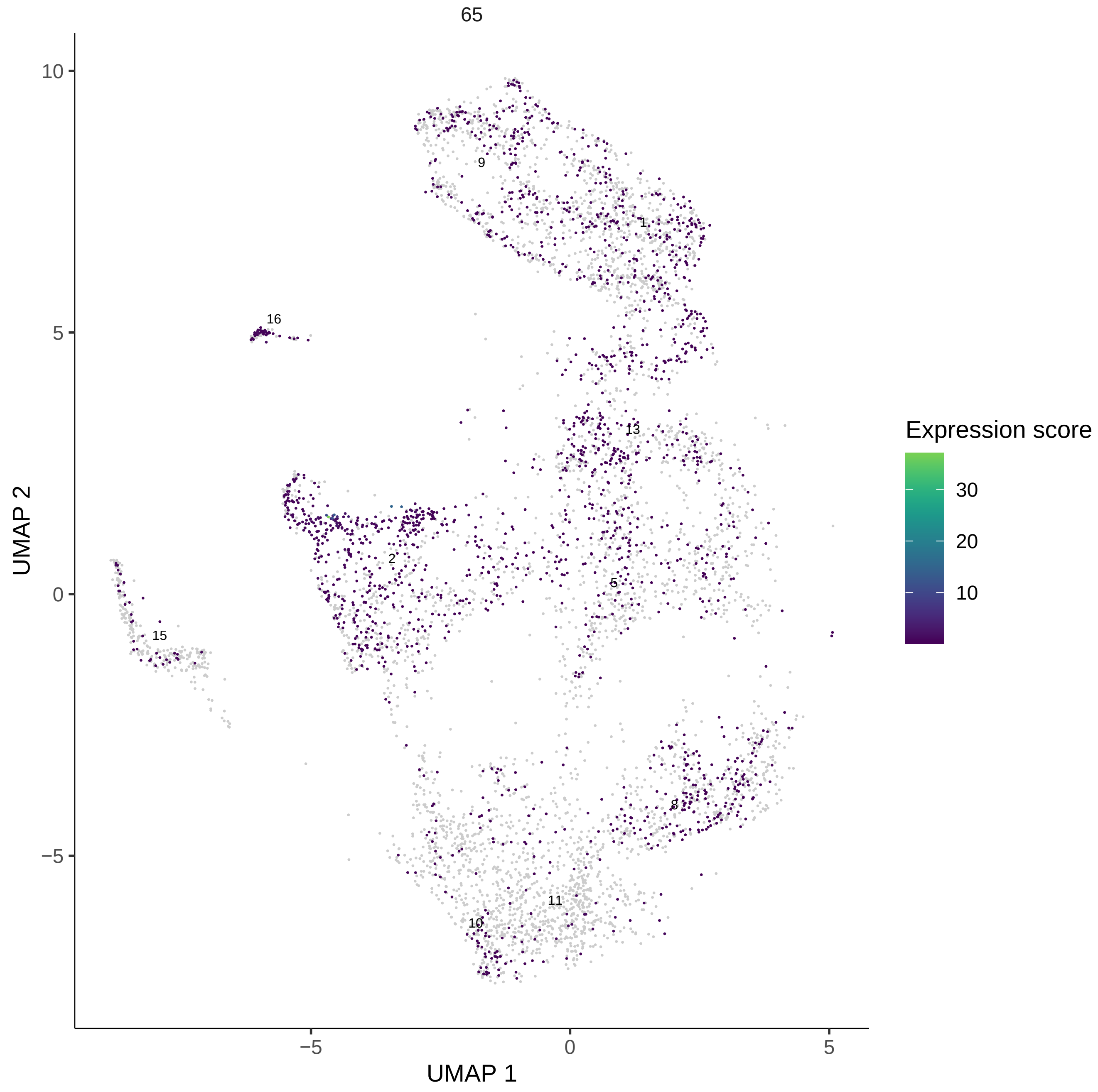

65
UMAP 2
UMAP 1
10
5
0
−5
−5
0
5
9
1
16
13
2
5
15
8
11
10
Expression score
30
20
10

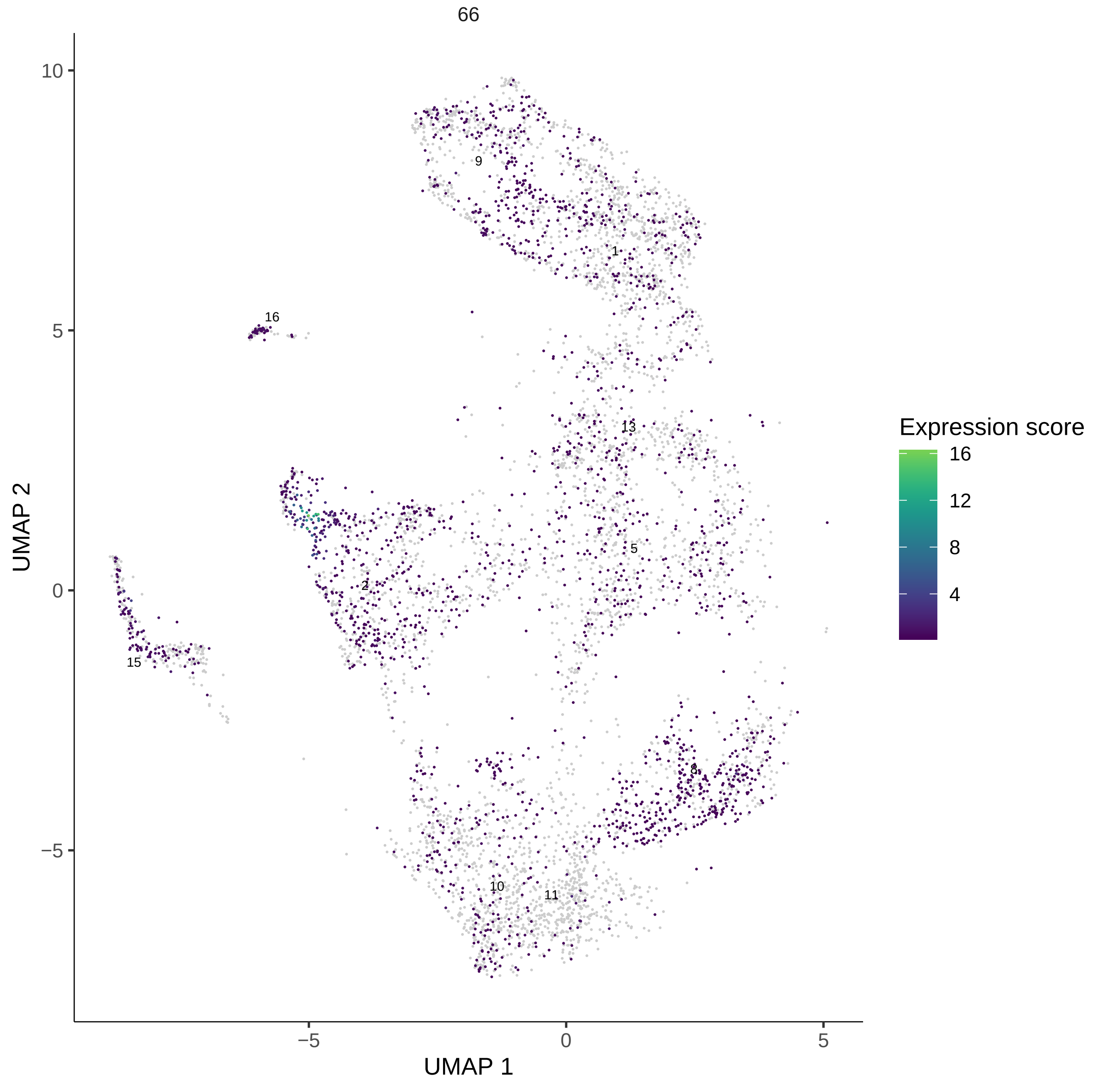
66
10
5
0
−5
UMAP 2
−5
0
5
UMAP 1
9
1
16
13
5
2
15
8
10
11
Expression score
16
12
8
4

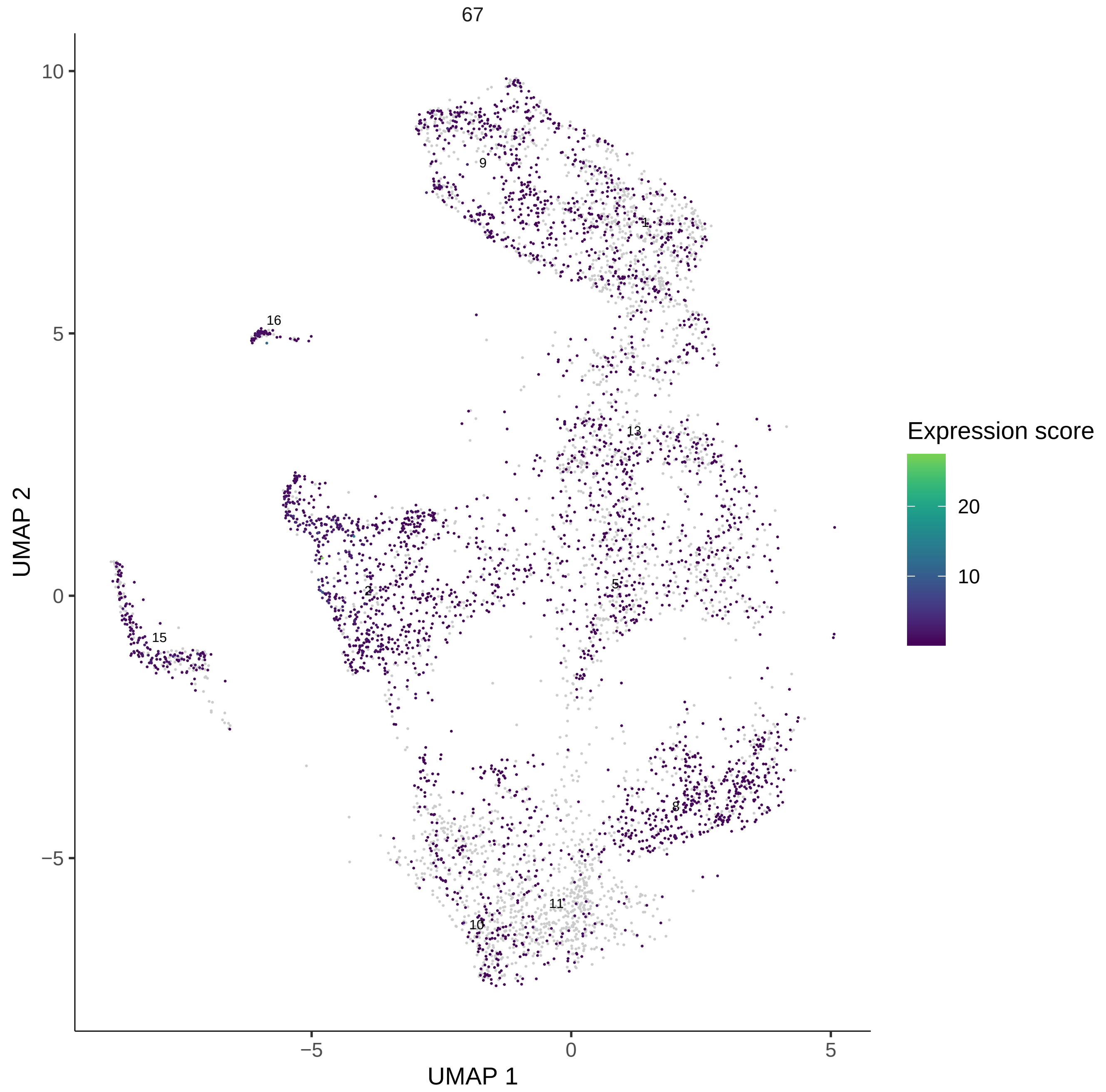
67
10
5
0
−5
−5
0
5
UMAP 1
UMAP 2
9
1
16
13
2
5
15
8
11
10
Expression score
20
10

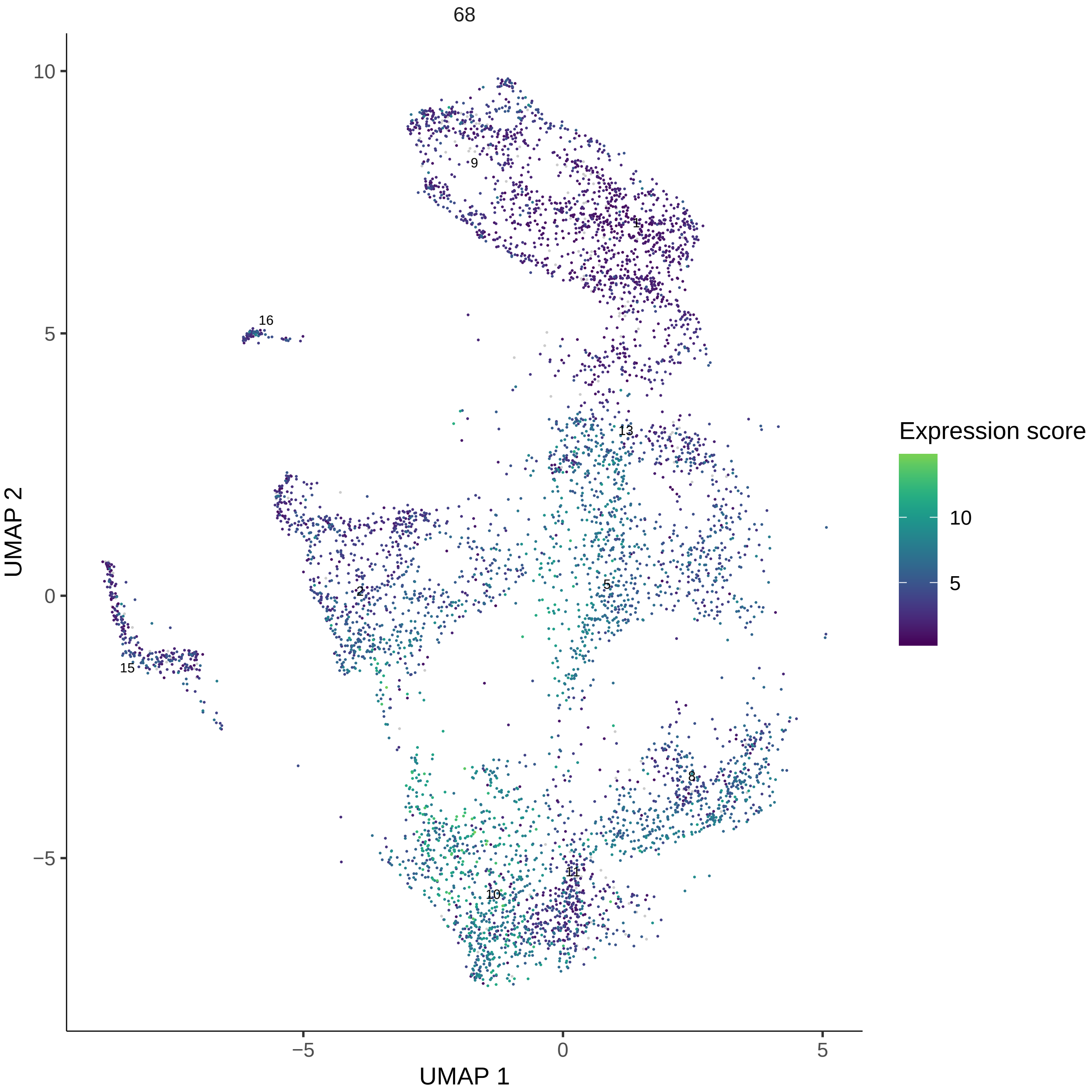

68
UMAP 2
UMAP 1
10
5
0
−5
−5
0
5
9
1
16
13
2
5
15
8
11
10
Expression score
10
5

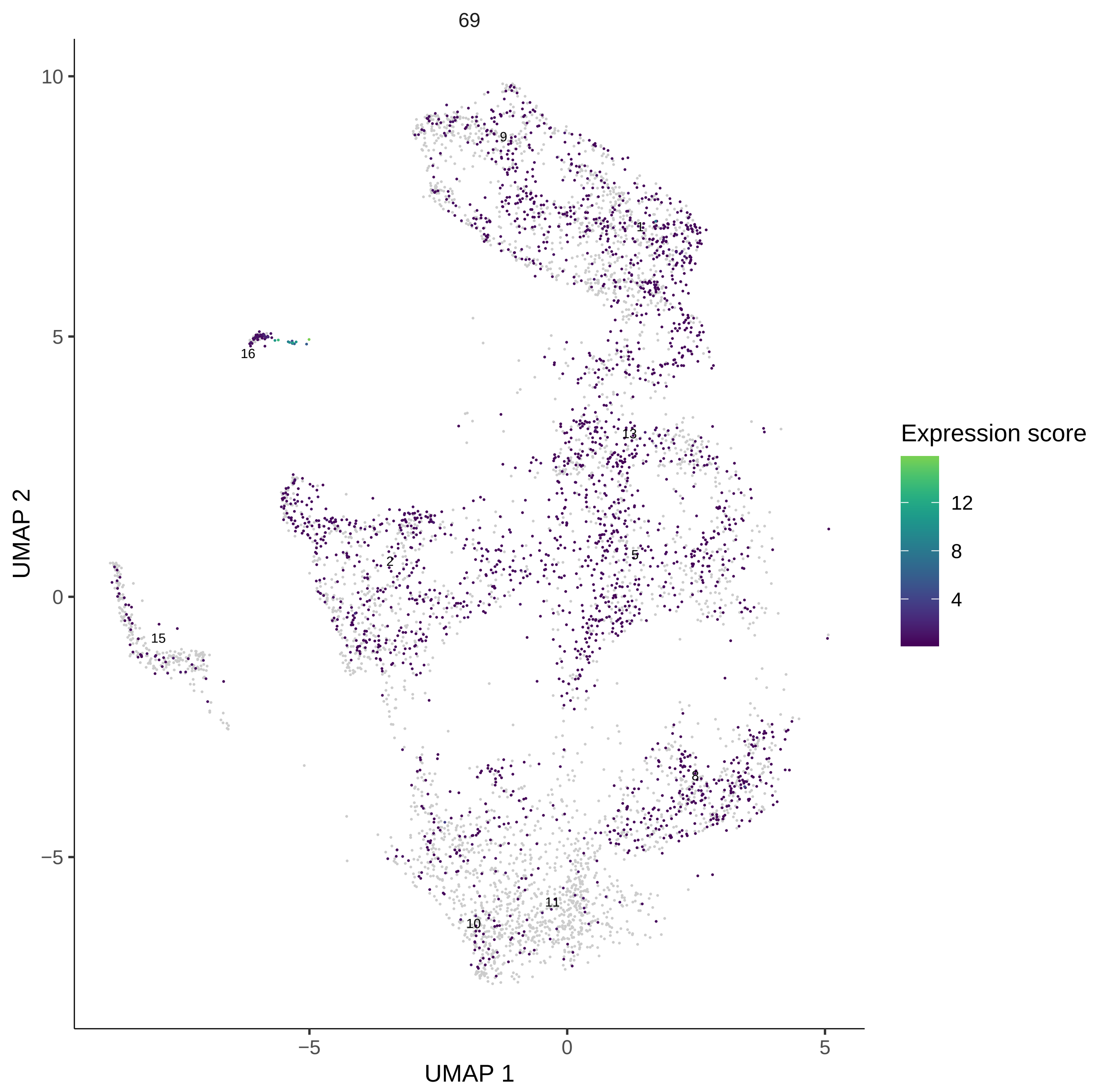
69
UMAP 2
UMAP 1
10
5
0
−5
−5
0
5
9
1
16
13
2
5
15
8
11
10
Expression score
12
8
4

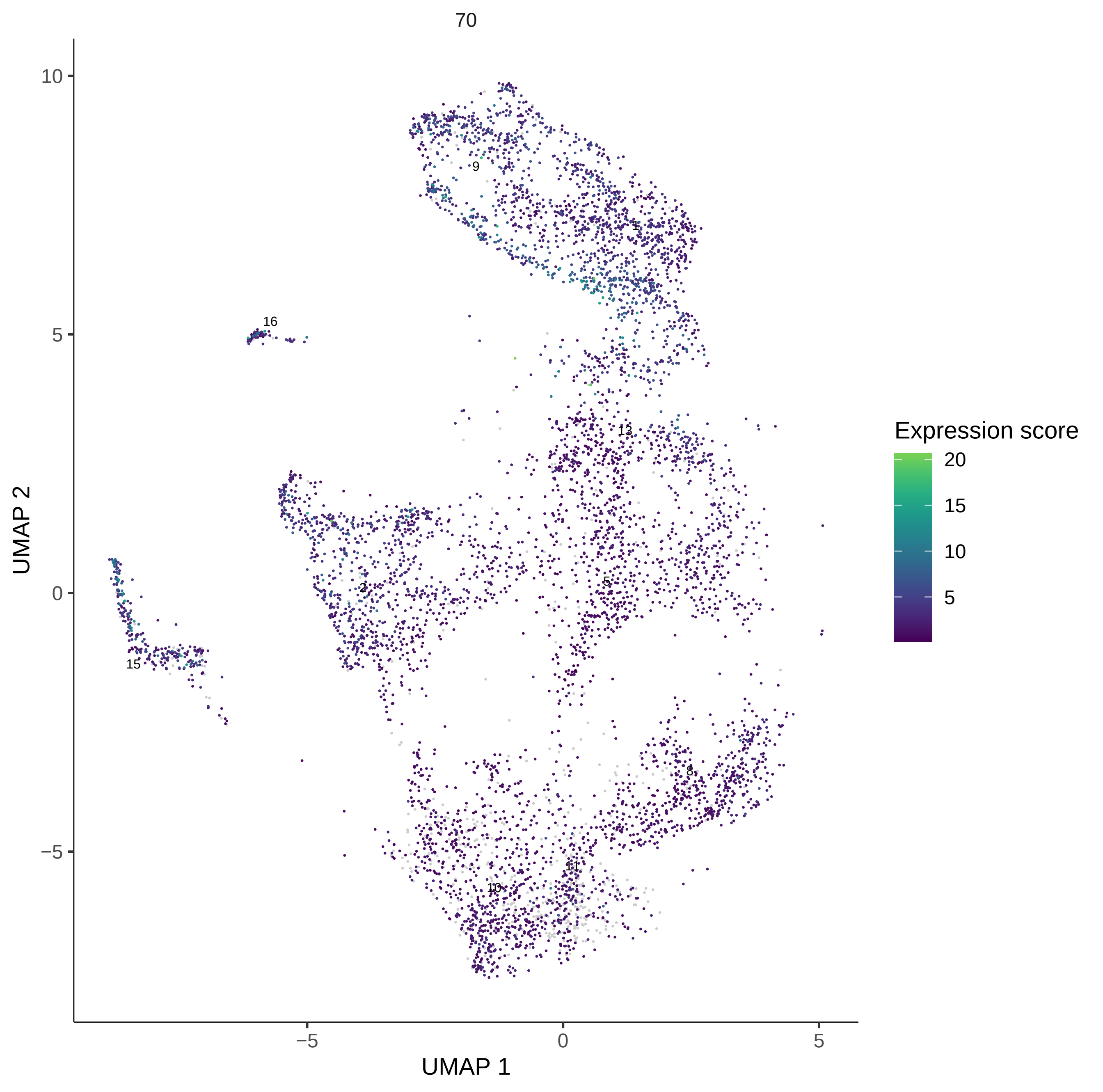
70
UMAP 2
UMAP 1
10
5
0
−5
−5
0
5
9
1
16
13
2
5
15
8
11
10
Expression score
20
15
10
5

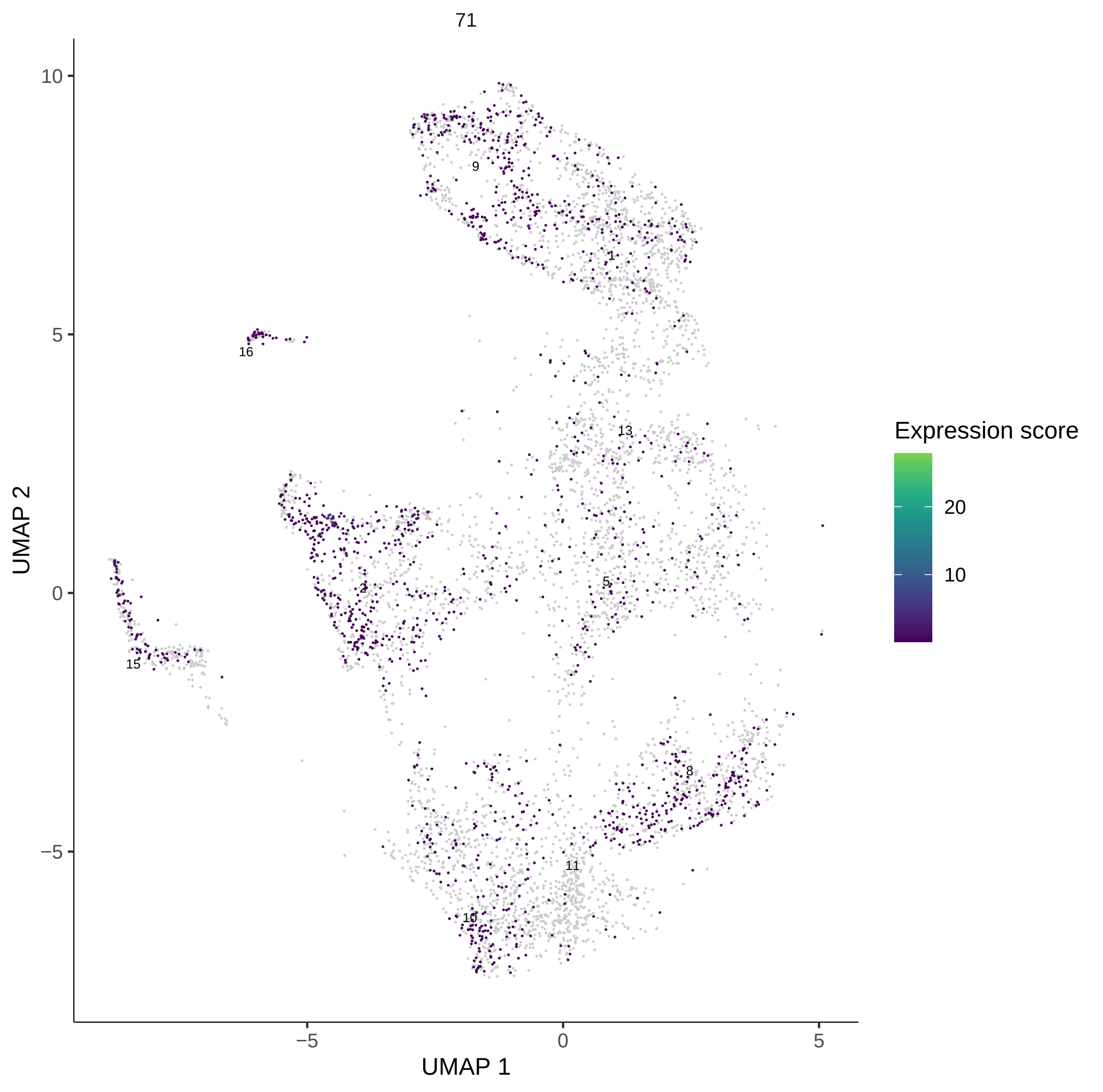
71
UMAP 2
UMAP 1
10
5
0
−5
−5
0
5
9
1
16
13
2
5
15
8
11
10
Expression score
20
10

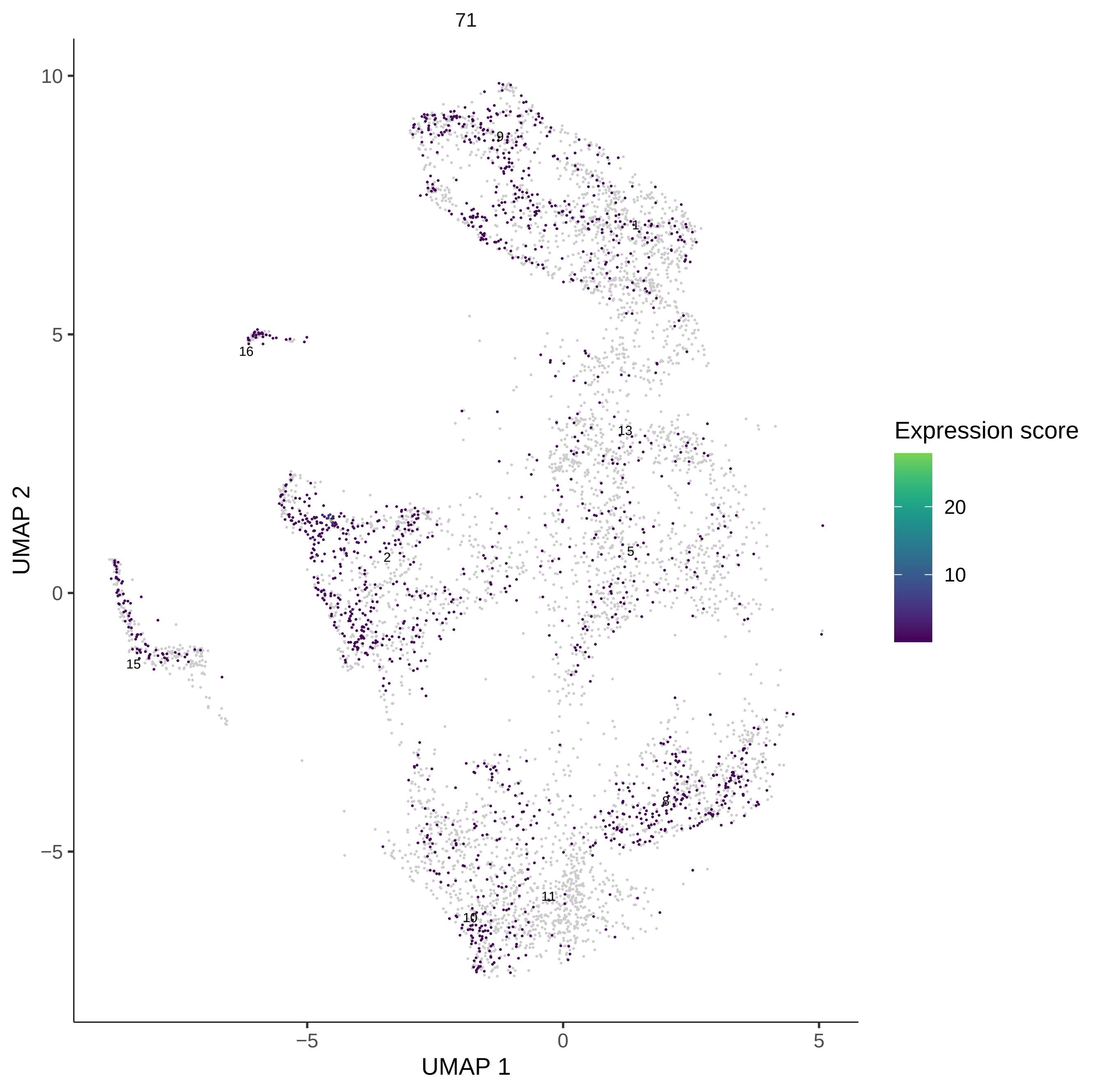

71
UMAP 2
UMAP 1
10
5
0
−5
−5
0
5
9
1
16
13
2
5
15
8
11
10
Expression score
20
10
